# Stability of Two-quartet G-quadruplexes and Their Dimers in Atomistic Simulations

**DOI:** 10.1101/820852

**Authors:** Barira Islam, Petr Stadlbauer, Michaela Vorlíčková, Jean-Louis Mergny, Michal Otyepka, Jiří Šponer

**Author notes:** **Corresponding Author:** Prof. Jiri Sponer.

## Abstract

G-quadruplexes (GQs) are four-stranded non-canonical DNA and RNA architectures that can be formed by guanine-rich sequences. The stability of GQs increases with the number of G-quartets and three G-quartets generally form stable GQs. However, the stability of two-quartet GQs is an open issue. To understand the intrinsic stability of two-quartet GQ stems, we have carried out a series of unbiased molecular dynamics (MD) simulations (∼505 µs in total) of two- and four-quartet DNA and RNA GQs, with attention paid mainly to parallel-stranded arrangements. We used AMBER DNA parmOL15 and RNA parmOL3 force fields and tested different ion and water models. DNA two-quartet parallel-stranded GQs unfolded in all the simulations while the equivalent RNA GQ was stable in most of the simulations. GQs composed of two stacked units of two-quartet GQs were stable for both DNA and RNA. The simulations suggest that a minimum of three quartets are needed to form an intrinsically stable all-*anti* parallel-stranded DNA GQ. Parallel two-quartet DNA GQ may exist if substantially stabilized by another molecule or structural element, including multimerisation. On the other hand, we predict that isolated RNA two-quartet parallel GQs may form, albeit being weakly stable. We also show that ionic parameters and water models should be chosen with caution because some parameter combinations can cause spurious instability of GQ stems. Some in-so-far unnoticed limitations of force-field description of multiple ions inside the GQs are discussed, which compromise capability of simulations to fully capture the effect of increase of the number of quartets on the GQ stability.

## INTRODUCTION

DNA and RNA can form diverse stable secondary structures depending on their sequence and environment.^1, 2^ Guanine-rich sequences can adopt G-quadruplex (GQ) structures by guanine-guanine Hoogsteen base pairing.^3^ Potential GQ-forming sequences have been found at telomeres, gene promoters, immunoglobulin heavy chain switch regions, hypervariable repeats and in genomically unstable and recombination-prone regions.^4^ The abundance of GQ-forming sequences at functional and gene controlling sites suggest their potential role in gene expression.^5^ Therefore, there is considerable interest in studying and modulating the formation of GQs as a means for controlling gene expression and other cellular processes.^6^

GQs are formed by stacking of two or more planar guanine quartets, creating the G-stem. The O6 atoms of all the guanines face to form a central electron-rich channel within the GQ where cations bind and coordinate with the O6 atoms. This coordination is essential for GQ stability.^7^ Bases not involved in G-stem belong to bulges, flanking nucleotides or participate in the loops linking the G-strands of the GQs.^3^

The GQs can be monomolecular, bimolecular or tetramolecular depending on the number of DNA or RNA molecules that form the GQ.^8, 9^ The topologies of GQs are highly variable as the G-strand orientation can be parallel or antiparallel and based on that GQs can fold into a parallel, antiparallel or hybrid topology.^10, 11^ The guanines in the DNA GQs can adopt *syn* or *anti* orientation.^10, 12, 13^ In the RNA GQs, the 2□ -hydroxyl group in ribose allows for more interactions with the bases, cations and water molecules thereby increasing their stability, as common in RNA molecules.^14-19^ It also restricts the orientation of the guanine bases favouring the *anti*-orientation. Therefore, the majority of the RNA GQs have been observed in the parallel-stranded topology with all bases in the *anti*-orientation.^15-18^

The loops further add to the diversity as the GQs can have propeller, lateral or diagonal loops.^8, 20-27^ The parallel-stranded GQ have propeller (double-chain reversal) loops as they are necessary to orient the following and preceding G-strands in a parallel arrangement.^8^ The antiparallel and hybrid GQs can have various combinations of the three loop types.^20-27^ The overall GQ topology dictates the loop combination as well as the allowable *syn*-*anti* patterns of the guanines. Na^+^ ions tend to coordinate within the plane while K^+^ ions coordinate between the planes of the G-quartets.^28, 29^ The cation-quadruplex interactions can influence kinetic and equilibrium differences between the different topologies.^28, 30, 31^

Stability of GQs is affected by the number of G-quartets in the stem.^14, 32-34^ It is commonly assumed that increasing the number of G-quartets usually stabilizes the GQ structure.^12, 14, 34, 35^ In DNA, stable GQ structures generally involve three quartets or more, although stable antiparallel two-quartet GQs with loop alignments or stacked triads have also been observed.^21, 36 27^ In such GQs, additional interactions formed by the loops or triads above and/or below the G-stems contribute to their stabilization.^21, 27, 36^ The solution structure of a two-quartet stacked dimer of RNA GQs has been solved, arguing that stabilization is provided by the dimerization of the two GQ units.^37^ Various experiments have suggested increased stability of RNA GQs over their DNA counterparts.^14, 38^ On the other hand, it was shown that substitution of even one guanine in the three-quartet telomere DNA quadruplex by abasic site or by any non-G base prevents its parallel folding under dehydrating conditions which otherwise favour parallel-stranded arrangement.^39, 40^ This could indicate low stability of two-quartet parallel-stranded DNA GQs. However, the intrinsic stability of isolated two-quartet G-stems cannot be fully clarified by the experiments, due to difficulty to separate the G-stem stability from the influence of additional interactions as well as due to structural resolution limits of the experiments. Still, understanding the stability of two-quartet G-stems is important. They may for example transiently occur during folding processes of larger GQs, in the initial fast phase of folding into the early distribution of GQs as well as in later transitions between different GQ folds towards the thermodynamics equilibrium.^41^ Thus, two-quartet GQs can have an impact on the folding processes. They also may provide regulatory functions due to possible equilibrium between the folded and unfolded state at physiological temperatures. In addition, the existence of these two-quartet GQs is of fundamental importance for all bioinformatics searches as their inclusion greatly increases the number of putative GQ-forming sequences in the genome.^42, 43^

The goal of this study was to assess the stability of the smallest two-quartet G-stems, with emphasis on the parallel arrangement. We have carried out a series of unbiased explicit-solvent molecular dynamics (MD) simulations (total of 505 μs). The two-quartet DNA and RNA G-stems are studied in isolation as well as in the context of GQ dimers. MD methodology, when wisely applied, can provide useful insights into various aspects of GQ structural dynamics and folding.^44^ While the contemporary MD is not robust enough to quantitatively evaluate the absolute stability of the GQs in a thermodynamic sense, the simulations should fairly well reflect different relative structural stabilities of GQ arrangements. It requires that the simulation time scale is sufficient to see structural perturbations and unfolding in the simulations, *i*.*e*., to assess the lifetimes (1/k_off_) of the studied arrangements.

We suggest that the all-*anti* parallel-stranded two-quartet DNA G-stem is not stable *per se*, and explain the reason for this behaviour. Nevertheless, such G-stems may still be involved in the formation of larger GQ structures. We also suggest that the RNA two-quartet G-stem is more stable. Simulations were further done for two-quartet DNA antiparallel G-stem to explain the difference in stability of various two-quartet G-stems.

To obtain additional insights into the balance of forces in the G-stems and the consequences of the force-field approximations, we have also carried out simulations of tetrameric parallel-stranded all-*anti* DNA three and four-quartet G-stem. We compared the performance of several water models and ionic parameters. Strikingly, our simulations show that several water-ion combinations lead to visible destabilization of G-stems, especially when using the more advanced four-site water models. Together with our previous study on three-quartet G-stems,^29^ these results illustrate the complexity of the choice of water models and ionic parameters in simulations of nucleic acids; it appears that no combination could be generally recommended as the best one. In addition, our simulations indirectly suggest that the pair-additive molecular mechanics (MM) description generally does not properly capture the stabilizing effect due to increasing the number of G-quartets in the stem.

## METHODS

### Starting structures

We took the 28-mer two-quartet stacked DNA GQ (PDB id 2N3M^45^, first model; Figure 1a), which is a solution structure of a synthetic construct d[(TGG)_3_TTGTTG(TGG)_4_T] that exhibits anti-proliferative activity *in vitro*. It consists of a dimer of two-quartet parallel-stranded GQ units stacked by their 3□-5□ interface and connected by a linker nucleotide. The simulations were carried out with the whole 2N3M structure, 2N3M with no bond between G15 and T16 of first and second GQ unit and also for its individual two-quartet GQ units (Figure 1a, 1b and 1c). The individual GQ units are called first and second GQ starting from the 5′-end of 2N3M. The model of parallel-stranded DNA G-stem d[GG]_4_ was made using the first two quartets of PDB structure 1KF1 (Figure 1d). The model of parallel-stranded DNA G-stem d[GG]_4_ with all 5′-guanines in *syn* orientation (Figure 1e) was made using the first two quartets of PDB structure 3TVB.^46^ The antiparallel DNA G-stem d[GG]_4_ was obtained from the solution structure of the thrombin binding aptamer (TBA) sequence d[(GGTTGGTGTGGTTGG)] GQ (PDB id 148D, first frame; Figure 1f).^21^

**Figure 1.**
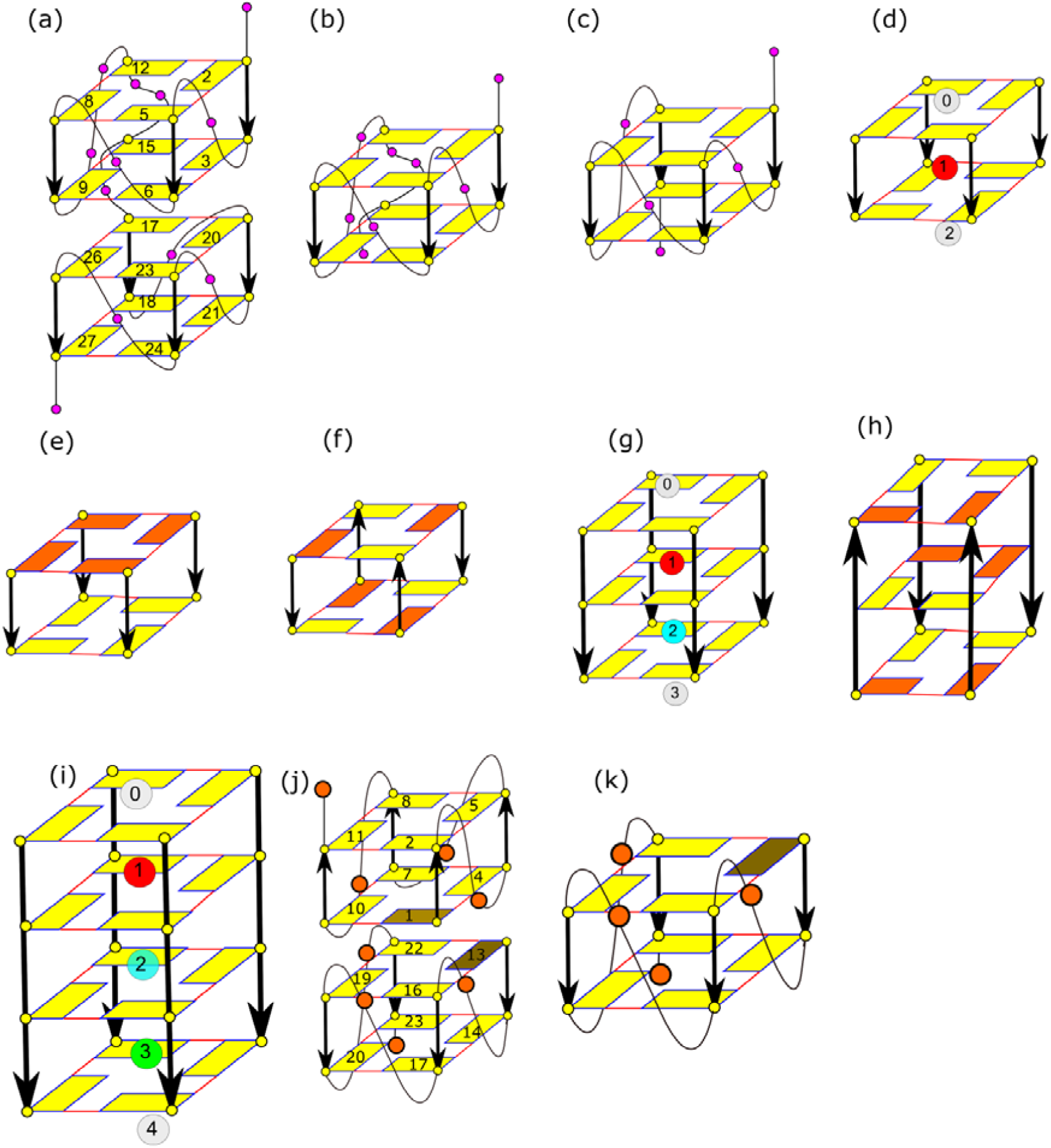
Starting structures used in the simulations. (a) 2N3M GQ with nucleotide numbering of guanines, (b) two-quartet GQ extracted from upper half of 2N3M, (c) two-quartet GQ extracted from lower half of 2N3M, (d) all *anti* parallel-stranded d[GG]_4_ G-stem, (e) parallel-stranded d[GG]_4_ with 5′-guanines in *syn* orientation, (f) antiparallel d[GG]_4_ extracted from 148D, (g) parallel-stranded d[GGG]_4_ from 1KF1, (h) antiparallel d[GGG]_4_ from 143D and (i) parallel-stranded d[GGGG]_4_ from 4R44. The RNA simulations were carried out using the dimer of (j) 2RQJ with nucleotide numbering of guanines and (k) monomer of 2RQJ GQ. The *anti* and *syn* guanines are shown in yellow and orange, respectively. The 5′-end of RNA structures is shown in tan. The loops are shown in thin lines while dark arrows show the backbone of G-stem. The yellow, magenta and orange circles show guanine, thymine and adenine bases, respectively. The solid red lines represent the Hoogsteen base pairing. Cation binding sites numbering of two-quartet, three-quartet and four-quartet GQs are marked in the panels (d), (g) and (i), respectively. Supporting Information Figure S1 shows the atomistic structures.

The coordinates for parallel- and antiparallel-stranded three-quartet system d[GGG]_4_ were obtained from the human telomeric DNA quadruplex (PDB ids 1KF1, chain A and 143D, model 1; Figure 1g and 1h).^8, 22^ For four-quartet tetramolecular DNA G-stem d[GGGG]_4_ simulations, we took the coordinates of guanine core from the X-ray structure of the d(TGGGGT) sequence (PDB id 4R44, chains A-D; Figure 1i).^47^ It is a tetramolecular GQ with three K^+^ in the GQ as the stabilizing ions; the structure is virtually identical to other available d[TGGGGT]_4_ structures.^48^ The coordinates for the dimer and monomer of two-quartet RNA GQ were taken from a solution structure of a dimer of two-RNA GQ units formed by the sequence 5′-r(GGA)_4_-3′ stacked through the 5′-5′ interface (PDB id 2RQJ, first model; Figure 1j).^37^ The head-to-head stacking means that quartets of the two subunits are of opposite directionality. For simulations of the monomer, we took the first GQ unit (Figure 1k). The hypothetical antiparallel RNA G-stem r[GG]_4_ was prepared directly from the DNA variant d[GG]_4_ (extracted from TBA, see above) by changing deoxyribose into ribose. Atomistic representations of the starting structures are shown in Figure S1 in Supporting Information.

### Comment on potential end effects caused by truncation

Simulations of nucleic acids are known to suffer from a not fully correct description of the helix termini, which leads to likely excessive end-fraying of double helices.^49, 50^ A similar concern can also be voiced about our simulations where we introduce some truncations of structures which lead to quartets containing chain termini. This should be taken into consideration when interpreting the results. However, terminal G-quarters in GQ simulations are more stable than terminal base pairs in duplex simulations, due to the stiffness of GQs and richer network of interactions. The “end fraying” issue for GQs is therefore less severe compared to duplex simulations. In addition, as our goal was to assess the intrinsic structural stability of G-stems in isolation, these simulations represent a key part of the simulation set. These structures may be experimentally relevant when considering tetramolecular complexes which do not involve any loop. Such intermolecular complexes may have anti-HIV activity for example as shown in ref.^51^ Understanding the stability of the core/stem is therefore of interest *per se*. We did not attempt adding (or keeping) 5□-terminal or 3□-terminal flanking nucleotides in the simulations of the truncated structures as these could be susceptible to excessive fluctuations and could introduce more uncertainty into the simulations compared to the presently studied systems. Note also that flanking bases in experimental studies of tetrameric quadruplexes are generally introduced to prevent multimerization rather than to stabilize the GQs *per se*.^14, 33, 34^

### MD simulations

The GQs were solvated in a truncated octahedral box with a minimal distance of 10 Å of solute from the box border, mostly in either SPC/E^52^ or OPC^53^ water model; some simulations were carried out in TIP4P-D water model.^54^ The simulations were carried out in 0.15 M excess NaCl or KCl, using mostly SPC/E specific (for SPC/E water model) or TIP4Pew specific (for OPC and TIP4P-D) Joung and Cheatham (JC) parameters.^55^ A minority of simulations was performed with Amber-adapted CHARMM22 parameters^54, 56, 57^ (for TIP4P-D), SPC/E specific (for SPC/E) and HFE or IOD OPC specific (for OPC) 12-6 LJ parameters developed by Li, Merz and coworkers (LM).^58^ The channel cations, when intended, were manually placed between the quartets and in the cavity between the stacked GQs. System preparation was done in xleap module of AMBER16.^59^ The list of structures and specific conditions of each simulation are presented in Table 1 and Tables S1 and S2 in the Supporting Information.

**Table 1.**
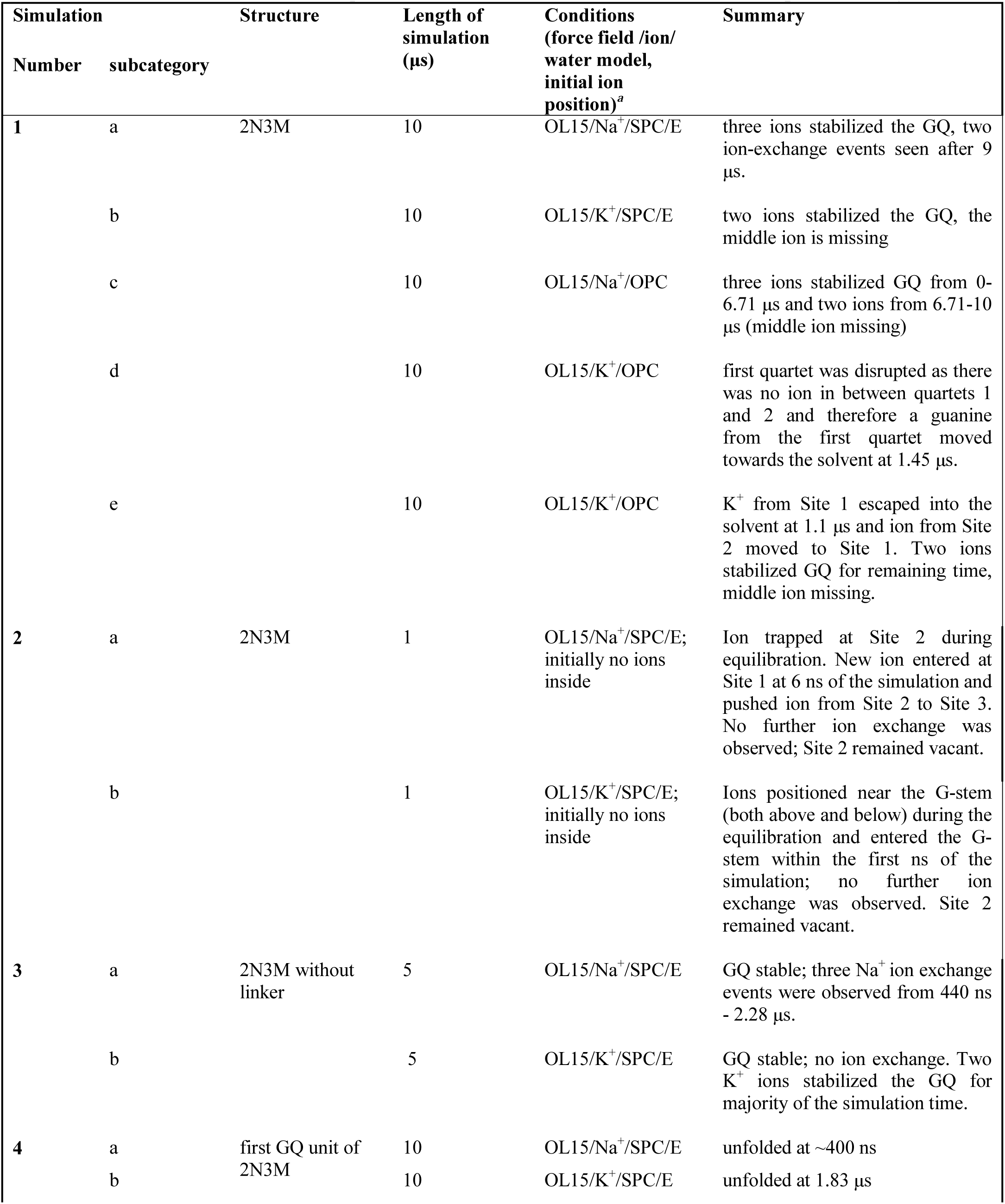
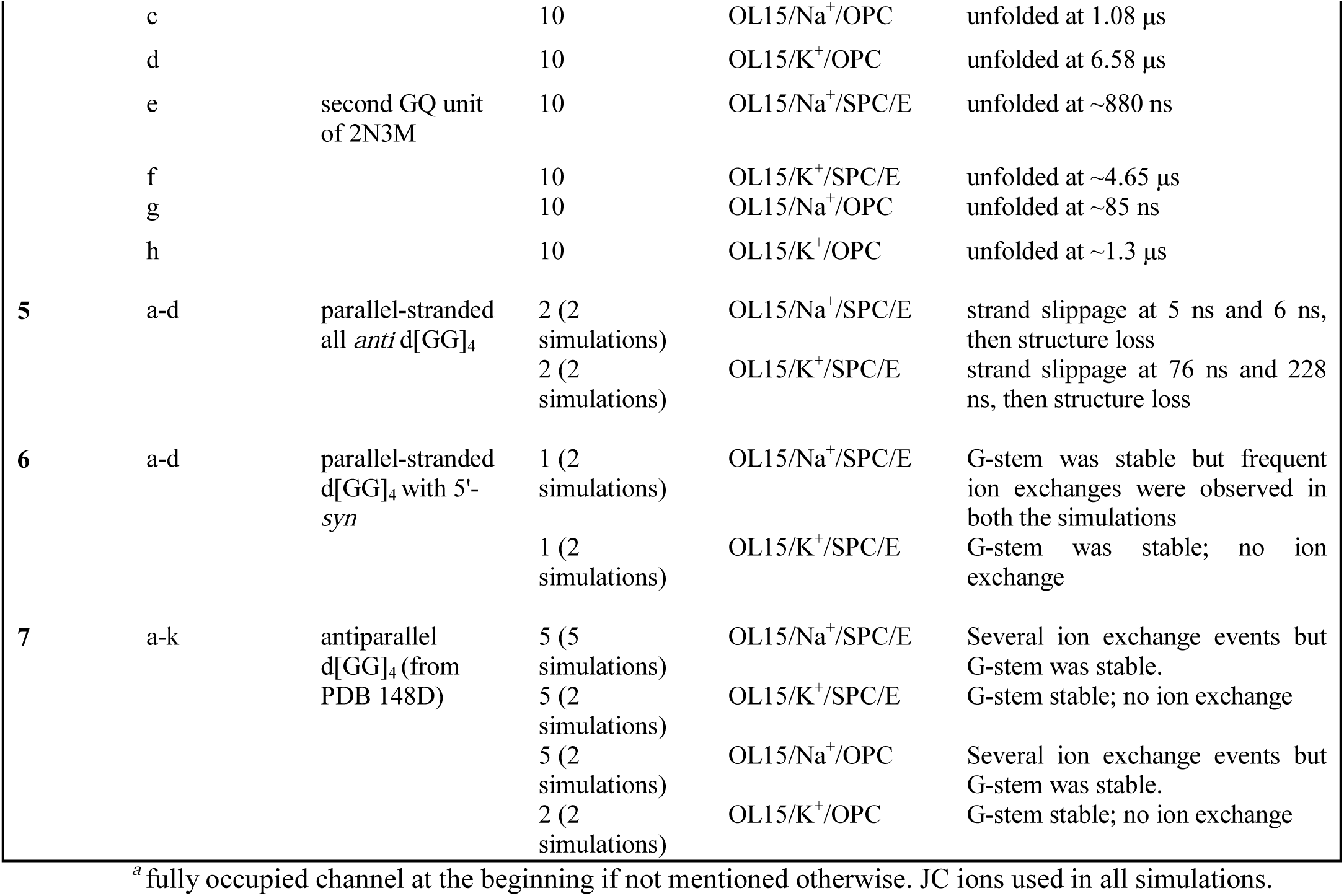
List of 2N3M and two-quartet DNA GQ simulations carried out in the present study

We used OL15^60^ and OL3 (known also as χ_OL3_)^61^ AMBER force field for DNA and RNA, respectively. These force-field versions include also earlier essential DNA^62-64^ and RNA^62^ modifications of the original AMBER Cornell *et al*. nucleic acids force-field;^65^ for more details see refs.^44, 66^

The starting structures were equilibrated using standard protocols. The system was first minimized with 500 steps of steepest descent followed by 500 steps of conjugate gradient minimization with 25 kcal mol^-1^ A^-2^ position restraints on DNA (or RNA) atoms. It was then heated from 0 to 300 K during 100 ps with constant volume and position restraints of 25 kcal mol^-1^ A^-2^. Minimization with 5 kcal mol^-1^ A^-2^ restraints followed, using 500 steps of steepest descent method and 500 steps of conjugate gradient. The restraints of 5 kcal mol^-1^ A^-2^ were maintained on DNA (or RNA) atoms and the system was equilibrated for 50 ps at constant temperature of 300 K and pressure of 1 atm. An analogous series of alternating minimizations and equilibrations followed using decreasing position restraints of 4, 3, 2 and 1 kcal mol^-1^A^-2^ consecutively. The final equilibration was carried out with position restraints of 0.5 kcal mol^-1^A^-2^ and starting velocities from the previous equilibration, followed by a short free MD simulation of 50 ps. Temperature and pressure coupling time constant during equilibration was set to 0.2 ps and for the last MD phase, it was set to 5 ps.

The simulations were performed with the PMEMD CUDA version of AMBER16.^59^ The electrostatic interactions were calculated using the Particle mesh Ewald method.^67^ The cut-off distance for nonbonded interactions was set to 9 Å. Covalent bonds involving hydrogen atoms were constrained using the SHAKE algorithm with a tolerance of 0.0001 Å.^68^ Hydrogen mass repartitioning of solute atoms was utilized allowing for an integration time step of 4 fs.^69^ Berendsen weak coupling thermostat and barostat were used to maintain constant temperature and pressure of 300 K and 1 atm, respectively.^70^ Langevin thermostat with a friction coefficient of 2 ps^-1^ was used in a few simulations. The final production runs without restraints were carried out for continuous length specified in Table 1 and Tables S1 and S2. Analyses of trajectories were performed using cpptraj^71^ module of AMBER and VMD^72^ program was used for visualization. An in-house script was used to identify cations residing in the binding sites of the GQs. Details of the script are provided in the Supporting Information. Gnuplot (Version 5) and Grace (Version 5.1.22) programs were used to produce the graphs.

### Justification of the sampling strategy

We have carried out 120 individual simulations. Rather than trying to repeat a small number of equivalent simulations multiple times to get standard statistics of events, we performed an as diverse set of simulations as possible. The diversity includes both the simulated systems and the water/ion parameters. We suggest that although a few individual simulations in the dataset may exhibit some rare behaviour (though we have no indication of that), all the reported trends consistently emerge from the overall set of simulations and are backed up by multiple simulations. This includes both the analysis of relative structural stabilities of different GQ systems as well as the assessment of properties of different water/ion parameter combinations. In other words, simulations of different GQ systems give a consistent relative ranking of different parameter combinations and simulations with different parameter combinations give a consistent assessment of relative stabilities of different simulated GQ systems.

## RESULTS

### Dimer of DNA GQs is stable in simulations (Simulations 1a - 1e)

We first performed simulations of the complete d[(TGG)_3_TTGTTG(TGG)_4_T] 2N3M structure. It forms a dimer of two-quartet parallel-stranded DNA GQs stacked by their 3′ / 5′ faces (Figure 1a). The residues G2:G5:G8:G12 and G3:G6:G9:G15 form the guanine quartets of the first GQ unit. T4 and T7 form single nucleotide propeller loops and T10 and T11 form a dinucleotide propeller loop. The residues T13 and T14 form a bulge between the two quartets, while T16 links the two GQ units. In the second GQ unit, G17:G20:G23:G26 and G18:G21:G24:G27 form the guanine quartets and T19, T22 and T25 form single-nucleotide propeller loops. Three cation binding sites in between the subsequent quartets are described further as Sites 1, 2 and 3 starting from the 5’-end. Additional cation binding sites were observed just above the 5′-quartet and 3′- quartet and these are described as Sites 0 and 4, respectively. We carried out 10 μs long simulations of 2N3M in SPC/E and OPC water models with both Na^+^ and K^+^ initiated with three ions inside the GQ (see Figure S2 for the RMSD developments).

The structure of 2N3M was maintained in the 10 μs SPC/E simulations with both Na^+^ and K^+^ (Simulations 1a and 1b). The Na^+^ ions were present in the three sites within the GQ (Figure 2a). Two ion exchange events were observed at 9.25 and 9.55 μs. In both events, Na^+^ ion entered from the solvent through the bottom of the stem (the fourth quartet) into Site 3 (Figure S3a). The approach of new cation to the bottom quartet was followed by a linear concerted upward movement of internal ions leading to escape of Na^+^ ion through the first quartet into the solvent and movement of the approaching cation into the channel. In the Simulation 1b, the K^+^ ion from Site 1 moved up at 1.2 μs and stayed on the top of the first quartet for further 300 ns and then escaped into the solvent. K^+^ from Site 2 moved to Site 1. Subsequently, Site 2 remained vacant (Figure S3b). Transient ion sites were formed just above and below the GQ (Figure 2b).

**Figure 2.**
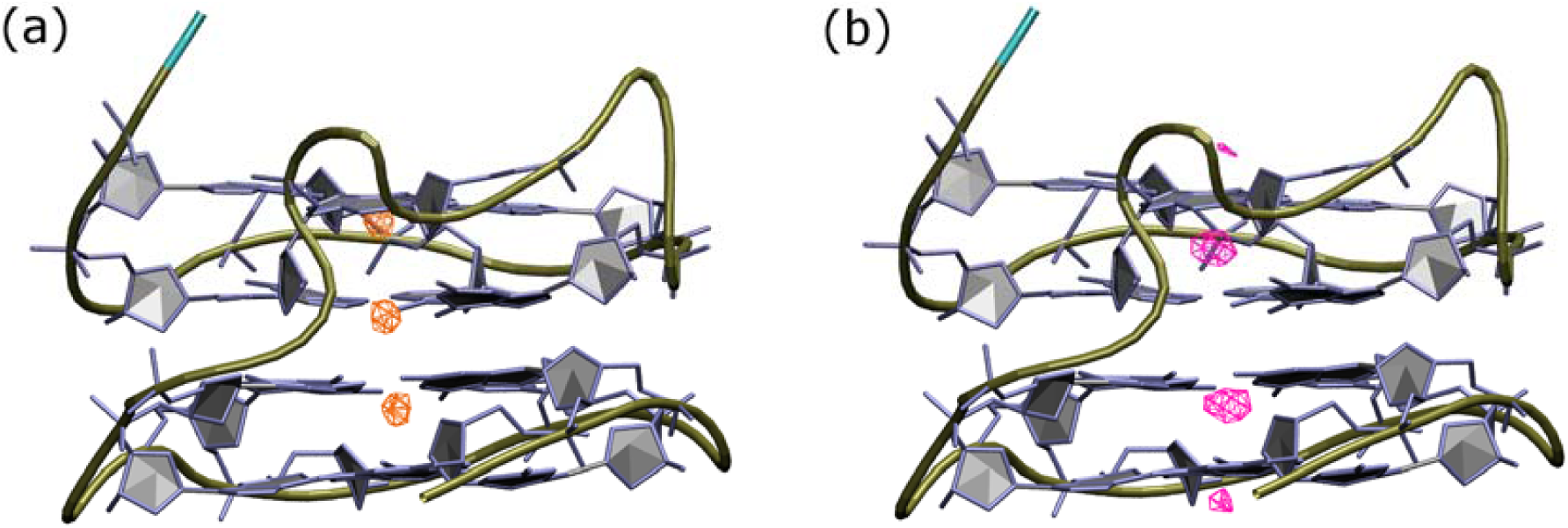
Cation binding sites (densities) observed in the MD simulations of 2N3M. (a) Three Na^+^ ions stabilized the GQ in the Simulation 1a while (b) two K^+^ ions stabilized the GQ for the majority of the Simulation 1b. The Na^+^ and K^+^ binding sites are shown in orange and pink, respectively. The backbone of the GQ is shown in tan tube. The guanine nucleotides of GQ are shown in silver ribbons and blue lines. The backbone of 5′-end of the GQ is shown in cyan. In the simulation in K^+^ ions, transient cation binding sites were developed above and below the GQ. A 2500 Å^-3^ threshold was used for density representation.

The 2N3M structure was also maintained in the simulation carried out in the Na^+^ ions with OPC water model (Simulation 1c). An ion exchange event was observed at 80 ns as a new Na^+^ ion entered from the bottom and occupied the Site 3 causing linear movement of cations within the GQ expelling the cation from the top quartet. The second ion exchange event was observed at 6.71 μs as a new Na^+^ ion entered into the Site 1 concurrently with the linear movement of cations and escape of a cation through the bottom quartet. Within 70 ns of this event, Na^+^ ion escaped from Site 3 and Na^+^ ion from Site 2 moved to occupy Site 3. Then only two Na^+^ ions remained in the GQ (Figure S3c).

In one OPC simulation carried out with K^+^ (Simulation 1d) the ion from Site 1 escaped into the solvent at 350 ns which was accompanied by a perturbation of the first quartet (Figure S3d). There was no concerted linear movement of K^+^ ions. At 1.45 μs, the first quartet was disrupted as G5 oriented to the solvent and the top quartet had only three guanines remaining (Figure S2f). The Site 1 remained vacant during the 10 μs long simulation. We carried out an additional 10 μs long simulation of 2N3M in OPC/K^+^ condition (Simulation 1e). The K^+^ ion from Site 1 moved to align in-plane with the first quartet within few ns of the start of the simulation and escaped into the solvent at 1.1 μs. This was followed by a subsequent movement of K^+^ ion from Site 2 to Site 1. The GQ was stable and Site 2 remained vacant till the simulation end (Figures S2e and S3e). These results show several things. In the simulations with K^+^, the cation tends to leave the ion-binding cavity between the two stacked GQs, leaving this site empty. Consistently with literature data,^29, 73^ observed ion exchanges with Na^+^ are typically concerted, *i*.*e*., an ion approach from the bulk to the channel entrance facilitates ion expulsion on the other side of GQ stem accompanied with a concerted shift of all ions. Loss of one quartet in the K^+^ OPC simulation, although not confirmed by the second simulation, is consistent with our recent preliminary tests showing, for model d[GGG]_4_ systems, that the OPC water model may under-stabilize the G-stems.^29^ In the present case, we observe a perturbation of GQ core of an experimentally determined structure, which is exceptionally rare in AMBER GQ simulations. ^74-78^

### Cations can quickly enter the two-quartet dimer through both top and bottom of GQ (Simulations 2a and 2b)

We also carried out 1 μs long simulations of 2N3M in the SPC/E water model but initially with no internally bound ions. In the simulation with Na^+^ ions in the bulk, an ion was trapped at Site 2 during equilibration. A new ion entered Site 1 after 6 ns of the start of the simulation and pushed ion from Site 2 to Site 3. No further ion exchange was observed. In the K^+^ simulation, two K^+^ ions positioned near both ends of the G-stem during the equilibration. They entered the G-stem within the first ns and no further ion exchange was observed. While Sites 1 and 3 were occupied, Site 2 remained ion-free (Figure S4a and S4b). The G-stem was stable in both simulations.

### The linker is not required for the stability of GQ dimer (Simulations 3a-3b)

We carried out Na^+^ and K^+^ SPC/E simulations of 2N3M upon cutting the bond between the G15 and T16, with standard 3′-OH and 5′-OH chain termini and with initially three bound ions. The GQ dimer stacking was stable. As for the simulations of the complete 2N3M structure, Na^+^ ions were more dynamic than K^+^ (Figure S4c).^29, 74-76, 79^ In Simulation 3a, three Na^+^ ion-exchange events were observed between 440 ns and 2.28 μs (Figure S4c). These ion exchanges were linked to the entry of a cation from the top or bottom quartet concurrent with a concerted movement leading to the escape of the ion from the top quartet. All three sites remained occupied (Figure S4c). In Simulation 3b, the K^+^ ion initially resided just below the fourth quartet rather than in Site 3. At 1.08 μs, this ion escaped into the solvent followed by the movement of ion from Site 2 into Site 3 (Figure S4d) and Site 2 remained unoccupied. The behaviour of cations was thus similar as in the complete 2N3M GQ.

### DNA two-quartet monomer GQs are unstable in simulations (Simulations 4a - 4h)

We carried out simulations of both the GQ units. The two-quartet GQs were unstable and unfolded in both the SPC/E and OPC water models between 80 ns - 7 μs (Table 1). Several unfolding pathways were observed. This may be partly caused by the differences in the ions and water models, though most likely it primarily reflects the known multidimensionality of the free-energy landscape.^44^ The first GQ unit unfolded by strand slippage and rotation leading to the formation of cross-like structure in Na^+^ and K^+^ ions in SPC/E water model and K^+^ ions in OPC water model (Table 1 and Figure 3). Strand slippage is a movement of a strand by one step up or down, reducing the number of full quartets.^80, 81^ By cross-like structure we mean a perpendicular or at least significantly inclined arrangement of the G-strands.^80-82^ In the Na^+^ and OPC simulation, unfolding was initiated by the loss of second quartet after excessive buckling. The first quartet then disintegrated within 2-3 ns leading to the complete unfolding.

**Figure 3.**
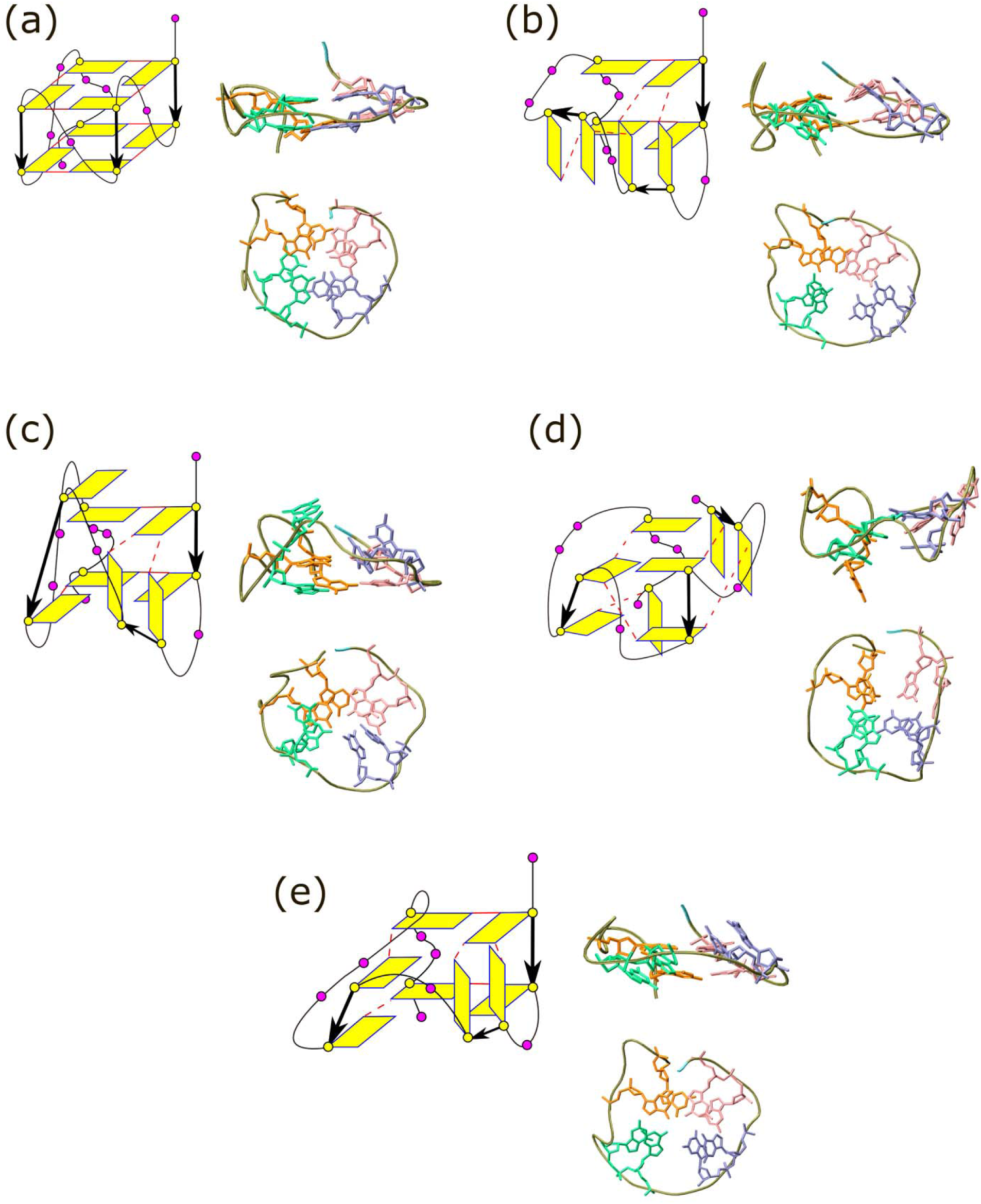
Simulations of the first GQ unit of 2N3M. The two-quartet structure unfolded in all the Simulations 4a-4d irrespective of the water model and stabilizing ion. Schematic and atomistic (both side and top) representations of (a) starting structure and main transient structure occurring during the unfolding in (b) Na^+^/SPC/E, (c) K^+^/SPC/E, (d) Na^+^/OPC and (e) K^+^/OPC are shown. See legend to Figure 1 for further details of the schematic representations. The solid red lines represent native Hoogsteen base pairing while dash red lines indicate any other hydrogen bonds. In the atomistic representations, the backbone of GQ is shown as tan tube with the 5′-end shown in cyan. The first, second, third and fourth strands of the GQ are shown in pink, blue, green and orange sticks, respectively. Cations are not shown in the figure. PDB files of the structures can be found in the Supporting Information.

In simulations of the second GQ unit as well, unfolding was marked by strand slippage and strand rotation in all the simulations (Table 1, Figures S5 and S6). Further details of the simulations are presented in the Supporting Information.

The simulations suggest that *parallel two-quartet DNA GQs are unstable per se as they unfolded in all conditions*. Obviously, the result can be affected by the force-field approximations (see the Discussion) but the unfolding is so decisive that we conclude that all-*anti* parallel-stranded two-quartet DNA GQs are intrinsically unstable. Hallmarks of the initial stages of unfolding events have been strand slippage and strand rotations.

### Tetramolecular two-quartet DNA GQs with *5′-syn* guanines are more stable than all-*anti* two-quartet DNA GQ (Simulations 5a - 5d, 6a - 6d and 7a - 7k)

We carried out two independent SPC/E simulations of two-quartet parallel-stranded d[GG]_4_ G-stems with all guanines in *anti* orientation in both Na^+^ and K^+^ ion conditions. The G-stems were highly unstable as strand slippage events followed by structure loss were observed at 5 and 6 ns in the Na^+^ ion simulations and at 76 and 228 ns in the two K^+^ ion simulations (Figure S7). The strand slippage events were not linked to any ion exchange events.

Simulations were then carried out for parallel-stranded d[GG]_4_ G-stems with all four 5′-guanines in *syn* orientation (Simulations 6a - 6d). The GQ was maintained in all the simulations (Figure 4a). As many as twenty ion-exchange events were observed in the simulations in Na^+^ ions while there was no cation exchange in the K^+^ ions simulations (Figures 4b-4e). The terminal *syn*-specific 5′-OH – G(N3) H-bond was dominantly sampled (Figure S8), as expected.^12, 83, 84^

**Figure 4.**
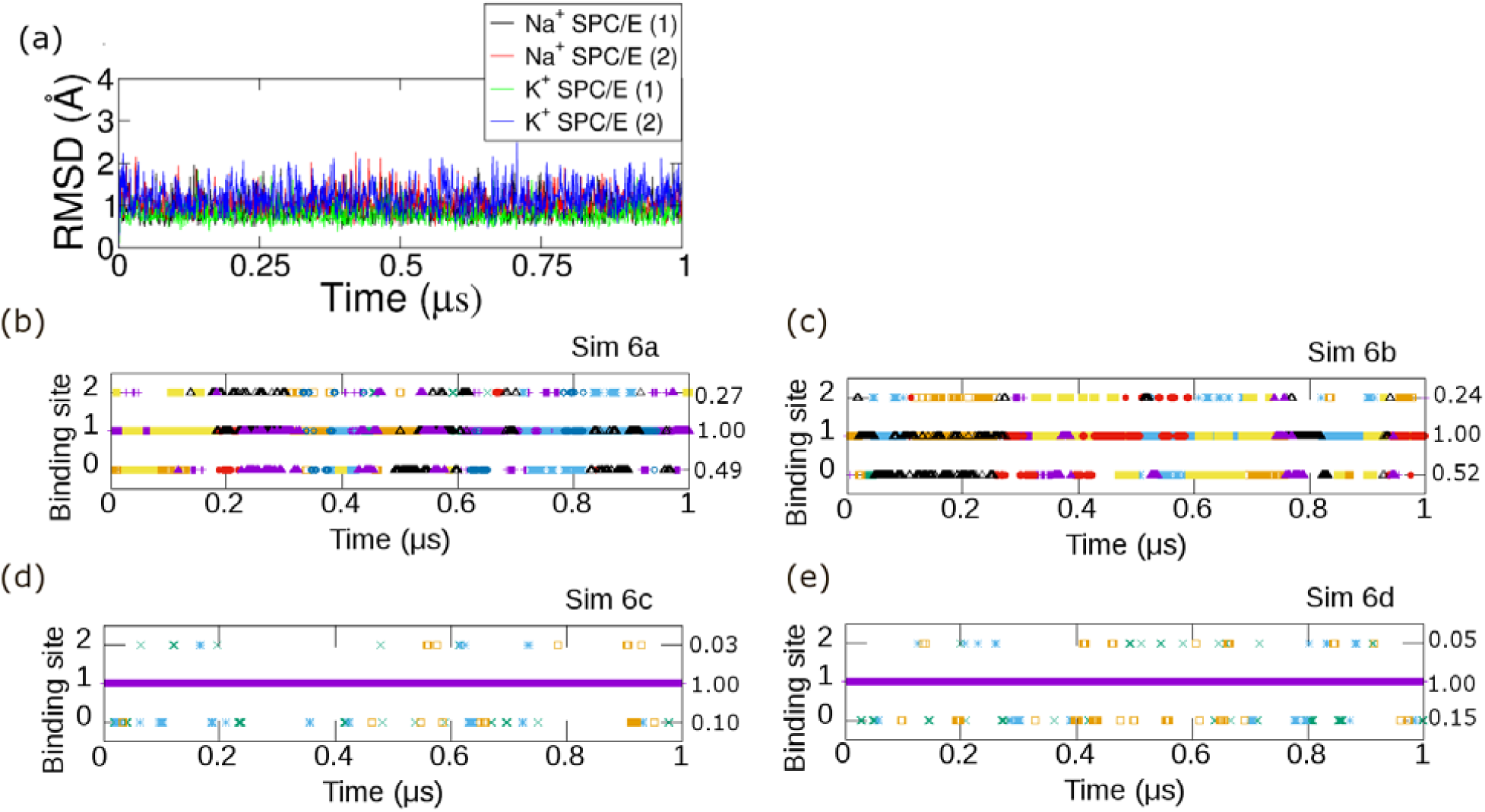
Backbone RMSD (a) and plots monitoring the cation binding (b-e) in parallel-stranded d[GG]_4_ with all 5′-bases in *syn* configuration in the Simulations 6a - 6d. Simulations 6a and 6b were carried out with Na^+^ ions and SPC/E water model while Simulations 6c and 6d were carried out with K^+^ ions and SPC/E water model. The cation binding site 1 corresponds to the typical position of cations in the channel of GQ between the two quartets. The sites 0 and 2 stand for the positions over the 5′-quartet and below the 3′-quartet, respectively. See panel (d) of Figure 1 for the representation of these sites. Distinct ions are represented by di□erent colors and symbols, and any change in the color/symbol indicates an exchange of an ion in the site. In the panels b - e, the numbers on the right show occupancies of the respective sites. Frequent cation exchange events were observed in the simulations carried out with Na^+^ ions while there was no cation exchange in the simulations carried out with K^+^ ions.

We also carried out simulations of two-quartet antiparallel d[GG]_4_ G-stem which maintained its structure in all five 5 μs long Na^+^ SPC/E simulations (Simulations 7a - 7e), although there were several (>10 per microsecond) ion-exchange events (Figure 5). The bases in the quartets were staggered for most of the simulation time but the two-quartet arrangement was maintained. In the two 5 μs long K^+^ SPC/E Simulations 7f-7g, no cation exchange was observed and the GQ was maintained although the quartets were again not entirely planar (Figure S9).

**Figure 5.**
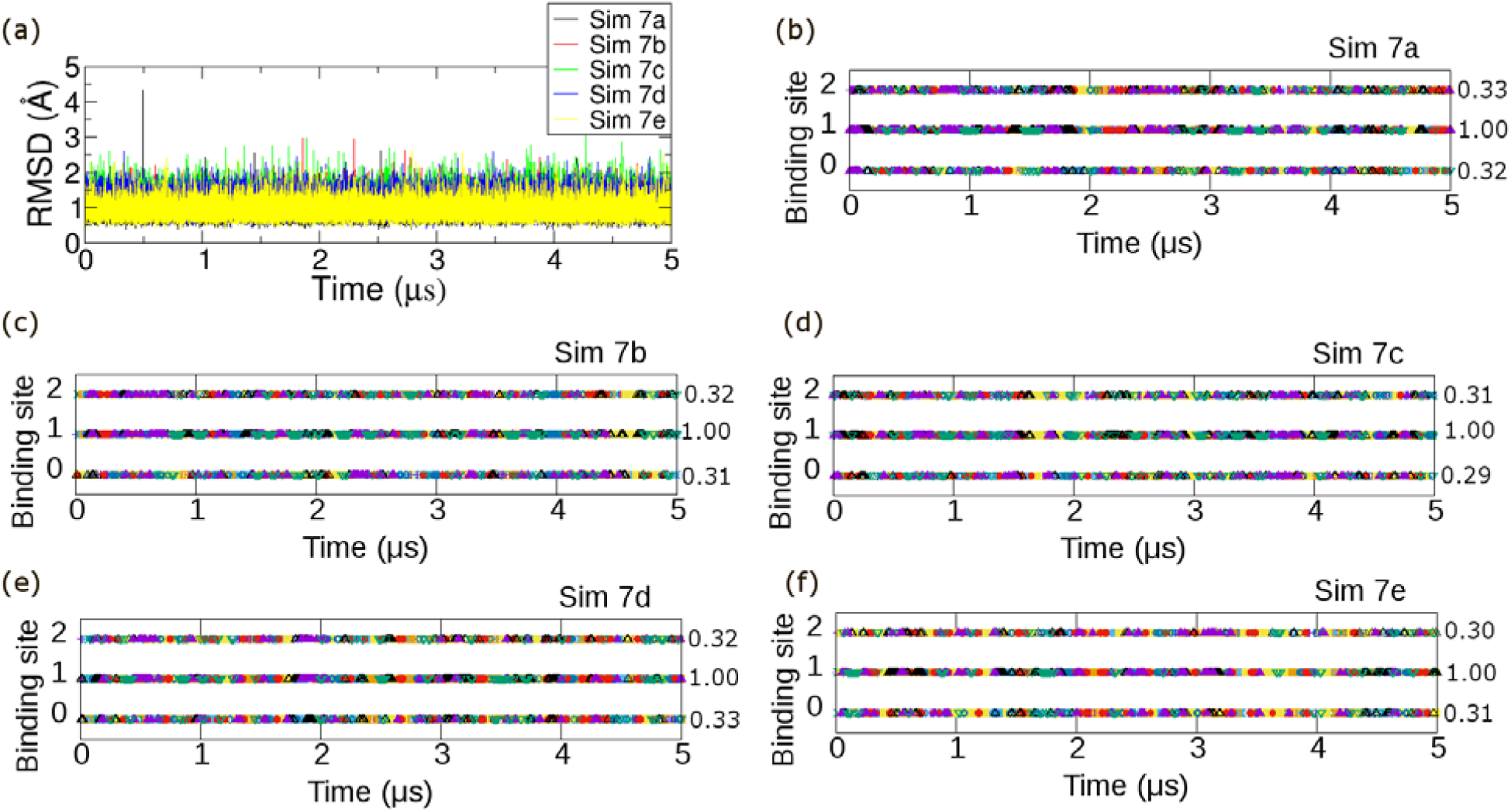
Backbone RMSD (a) and plots monitoring the cation binding (b-f) in antiparallel d[GG]_4_ G-stem in the Simulations 7a – 7e in the presence of Na^+^ ions and the SPC/E water model. The plots (b-f) reveal very fast cation exchanges in all the simulations. The cation binding site 1 corresponds to the typical position of cations in the channel of GQ between the two quartets. The sites 0 and 2 stand for the positions over the 5′-quartet and below the 3′-quartet, respectively. In the panels b - f, the numbers on the right show occupancies of the respective sites. For more details see the legends of Figure 4.

The two-quartet antiparallel G-stem was also stable in simulations in OPC water model (Simulations 7h-7k). In the 5 μs long simulation in Na^+^ ions there were continuous ion exchange events (Figure S10) but in the 2 μs long simulation in K^+^ ions no ion exchange was observed (Figure S11). This behaviour of Na^+^ and K^+^ ions was thus consistent across SPC/E and OPC water models (Figures 5, S9-S11).

In summary, all-*anti* parallel-stranded GQs unfolded in all the simulations. In contrast, greater stability of both parallel and antiparallel d[GG]_4_ G-stem with 5□-*syn* bases suggests that the opposite directionality of quartets hindering the strand slippage possibly aided also by the intramolecular 5□-OH – G(N3) H-bonds stabilizes the G-stem.

### Dimer of RNA two-quartet GQs is stable (Simulations 8a - 8f, 9a - 9b, 10)

We have carried out six 6-10 µs SPC/E K^+^ simulations, two SPC/E Na^+^ simulations and one OPC Na^+^ simulation (the latter is the potentially least stable combination of parameters) of the dimer of two-quartet RNA GQ 2RQJ. Various initial ion positions were considered (Table S1). Universally, all simulations were stable, the ions quickly occupied the Sites 1 and 3, Site 2 was vacant, and no ion exchanges occurred in the whole dataset once the final distribution of the ions was obtained (Figures S12 - S16). The results are described in details in Supporting Information.

### Two-quartet all-*anti* RNA GQ is more stable than the equivalent DNA GQ (Simulations 11a −11d, 12a-12d, 13, 14a-14c, 15a-15c, 16a-16d, 17 and 18a-18d)

We carried out four 10 μs K^+^ SPC/E simulations of a monomer of 2RQJ (Simulations 11a-11d). The two-quartet RNA GQ was stable in all the simulations and no ion exchange was observed (Figure 6). The terminal adenine (A12) was stacked below the strand 4 of the GQ as in the starting structure. The loop bases showed some dynamics but could not form any stable alignment with the G-stem. They either aligned in the groove of the GQ by forming transient hydrogen bonds with the G-stem bases or were exposed to the solvent.

**Figure 6.**
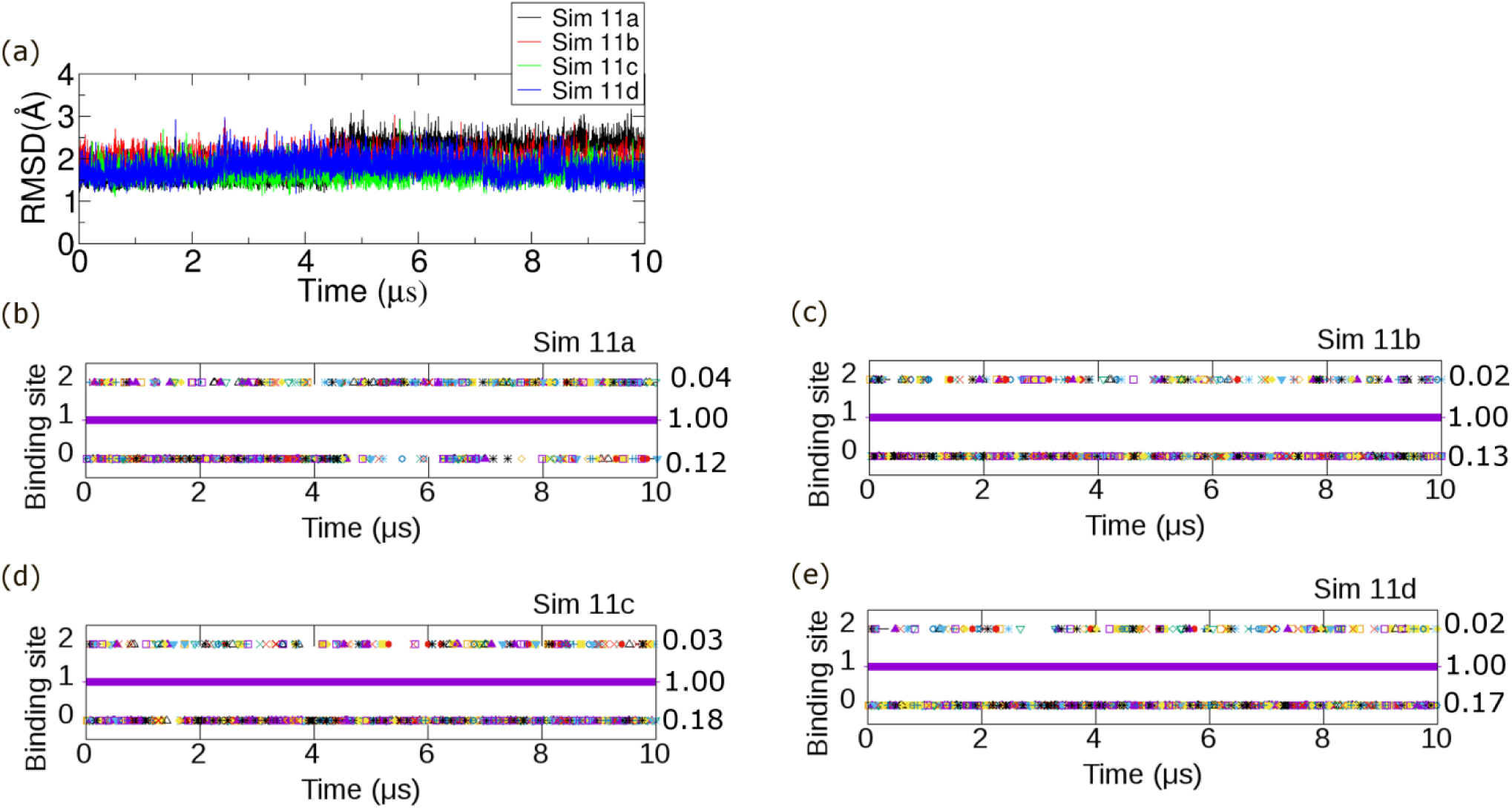
Backbone RMSD (a) and plots monitoring the cation binding (b-e) in the K^+^/SPC/E Simulations 11a - 11d of two-quartet GQ unit of 2RQJ. (a) The backbone RMSD of the GQ is shown in different colors for each simulation; the minor RMSD fluctuations are due to the loop dynamics. The plots (b-e) reveal that no cation exchange was observed at Site 1. The cation binding site 1 corresponds to the typical position of cations in the channel of GQ between the two quartets. The sites 0 and 2 stand for the positions over the 5′-quartet and below the 3′-quartet, respectively. In the panels b - e, the numbers on the right show occupancies of the respective sites. For more details see the legends of Figure 4.

We also carried out simulations of a monomer of 2RQJ without the 3′ flanking base in Na^+^ and K^+^ ions and in both SPC/E and OPC water models (simulation series 12 - 15). The GQ was maintained in three out of four 5 μs long Na^+^ SPC/E Simulations 12a-12d (Figure S17). The GQ unfolded in one simulation at 4.27 μs (Figure S17b). The χ angle of G1 attained *syn* orientation at 4.27 μs and G2 moved towards the solvent. The first propeller loop was lost as G2 moved to stack over G1 while G1 could not re-attain *anti* orientation till the end of the simulation. G1 also sampled the terminal *syn*-specific 5′-OH – G(N3) H-bond when it was in *syn* orientation. More than 20 ion exchange events were observed in all four of these simulations (Figure S18). The GQ also was stable in the 1 μs long K^+^ SPC/E Simulation 13. No ion exchange was observed in this simulation and the loop bases showed similar dynamics as in the Simulations 11a-11d (Figure S19).

We carried out three 1 μs long simulations (14a-14c) of the same RNA GQ in Na^+^ ions and OPC water model. In Simulation 14a, GQ was stable and loop bases behaved similarly to the equivalent stable SPC/E simulations (Figure S20). An ion exchange event was observed at 750 ns. However, the GQ unfolded in Simulations 14b and 14c (Figure S20). In Simulation 14b, a cross-like structure was formed due to the buckling and rotation of strands 1 and 4 at 400 ns. In Simulation 14c, strand 4 formed a cross-like structure at 130 ns leading to the unfolding of GQ. The unfolding was not linked to any cation exchange movement. In contrast, the RNA GQ was stable in three 1 μs K^+^ OPC simulations (15a-15c) and no cation exchange event was observed (Figure S21).

In brief, simulation series 11-15 suggests *universally enhanced stability of parallel-stranded all anti two-quartet RNA GQs over DNA GQs*. Previous works have shown that 2□-hydroxyl group in ribose sugar allows for more interactions with the bases, cations and water molecules thereby increasing the stability of the RNA GQ.^14, 18^ The GQ was stable in all K^+^ simulations. The two-quartet RNA GQ was able to maintain its structure in Na^+^ SPC/E simulations despite the multiple ion exchange events. The simulations nevertheless reveal that the combination of OPC water model with JC TIP4Pew Na^+^ ions destabilizes the RNA GQ compared to the other water-ion combinations. The cation exchange events may alter the energy of GQ due to change in solvation, ion-coordination and structural rearrangements within the GQ.^85^ However, the unfolding of GQs in the present OPC Na^+^ simulations could not be linked to any cation movements.

Supporting Information including Figures S22-S25 presents data of some less essential simulations, namely Simulations 16a-16d of 2RQJ RNA monomer using TIP4P-D water model and CHARMM22 ions, Simulation 17 substituting thymine by adenine in the second GQ monomer of 2N3M and Simulations 18a-18d of antiparallel RNA G-stem.

### Tetramolecular three-quartet antiparallel DNA G-stem is more stable than parallel-stranded G-stem (Simulations 19a - 19b, 20a - 20b and 21a - 21d)

In order to see the effect in adding one quartet (and ion), we have carried out a set of parallel-stranded d[GGG]_4_ simulations. In the two Na^+^ SPC/E simulations, strand slippage was observed at 1.6 μs and 530 ns, respectively (Figures S26a and S26b). However, the G-stem was entirely stable in the two 6 μs and 5 μs long K^+^ SPC/E simulations (Figures S26c and S26d). In Simulation 20a, a channel K^+^ was exchanged at the 5′-end at 4.8 μs while in the Simulation 20b it was exchanged at the 3′-end at 273 ns (Figures S27b). In both cases, the cation-leaving site was vacant for ∼25-30 ns followed by the entry of new cation from bulk at the same site. Thus, using the SPC/E water model the G-stem was stable in K^+^ ions but have shown a strand slippage in Na^+^ ion simulations.

The d[GGG]_4_ antiparallel G-stem was fully stable in SPC/E simulations with both ion types (Figure S28). In the Na^+^ simulation <10 ion exchange events were observed while no ion exchange was observed in the K^+^ simulation (Figure S29). In the Na^+^ OPC simulation (Simulation 21c), ion between the middle and top quartet escaped into the solvent at 65 ns. This resulted in instability of the quartets and G12 from the first quartet moved towards the solvent at 185 ns (Figure S28a). However, the G-stem was maintained in the Simulation 21d with K^+^ ions in OPC water model (Figure S28b).

To summarize, the all-*anti* parallel-stranded three-quartet G-stem was more stable than equivalent two-quartet G-stem, as we have seen rapid complete unfolding in all simulations of parallel d[GG]_4_ (see above). There is, however, a tendency for strand slippage in the presence of Na^+^ and for occasional K^+^ ion exchange. The antiparallel d[GGG]_4_ is evidently more stable than the all-*anti* parallel-stranded d[GGG]_4_. Additional simulations of the all parallel all-*anti* [GGG]_4_ in diverse conditions can be found in ref.;^29^ the SPC/E results are consistent with the present simulations while OPC simulations show lower stability compared to the SPC/E water model.

### Simulations of tetramolecular DNA four-quartet parallel-stranded all-*anti* G-stem show excessive ion dynamics (Simulations 22a - 22e, 23a - 23e, 24a - 24e, 25a - 25e and 26a - 26g)

In order to obtain more insights into the balance of forces in GQs and also to better understand the force-field limitations, we have carried out a series of simulations of parallel-stranded all-*anti* d[GGGG]_4_ tetrameric G-stem with three ions in the channel. The d[GGGG]_4_ simulations can be compared with the 2N3M simulations (simulation series 1-3) and with d[GGG]_4_ simulations (simulation series 19-21). The d[GGGG]_4_ all parallel all-*anti* G-stem has not been studied extensively before by long time-scale MD and, intuitively, one would expect it to be very stable. However, the results show a surprisingly complex picture.

The G-stem was maintained in all ten 1 μs long simulations in the SPC/E water model with either Na^+^ (Simulations 22a-22e) or K^+^ (Simulations 23a-23e). However, in four out of five Na^+^ simulations, <10 ion exchange events were observed (Figure S30a and S30c-30d) while no ion exchange was seen in one simulation (Figure S30b). All the ion movements were concerted. The Na^+^ ions in the channel were mobile, confirming that the stems sample diverse subtly different heterogeneous substates, interconnecting the detailed ion dynamics/distribution with modest differences in the internal G-stem conformation.^83^ The Na^+^ ions were mostly residing *close to* rather than *in between* the quartet planes (Figures S30 and S31).

In all the SPC/E simulations with K^+^ there has been a visible tendency to reduce the number of the ions in the channel to two (Figure S32), indicating structural stress associated with the three closely-spaced cations (see Discussion). This has not been observed in the above (simulation series 19-21) and earlier^29^ simulations of the d[GGG]_4_ with the same force-field parameters, *i*.*e*., two K^+^ are stable in d[GGG]_4_. In early stages of the d[GGGG]_4_ K^+^ simulations, the ion between quartet 1 and 2 (Site 1) was pushed above the stem (above its 5′-end). After the first shift of cation from Site 1, Site 2 cation moved to occupy Site 1 in one simulation and in another one the K^+^ ion returned back to Site 1 (Figures S32d & S32e). Otherwise, the cations stayed above the first quartet throughout the simulations (Figures 7 and S32). As this newly formed “outer” ion-binding site was exposed to the solvent, four to >20 cation exchanges with bulk ions were observed in the 1 µs simulations (Figure S32). Additionally, in one simulation, the bottom ion was lost to the solvent. Then the ion from the central cavity Site 2 moved to Site 3 and the middle site remained vacant for most of Simulation 23e (Figure S32e), thus resulting in only two ions in the stem. The K^+^ ions that remained within the G-stems mostly stayed in the inter-quartet cavities, as expected for the K^+^ ions.

**Figure 7.**
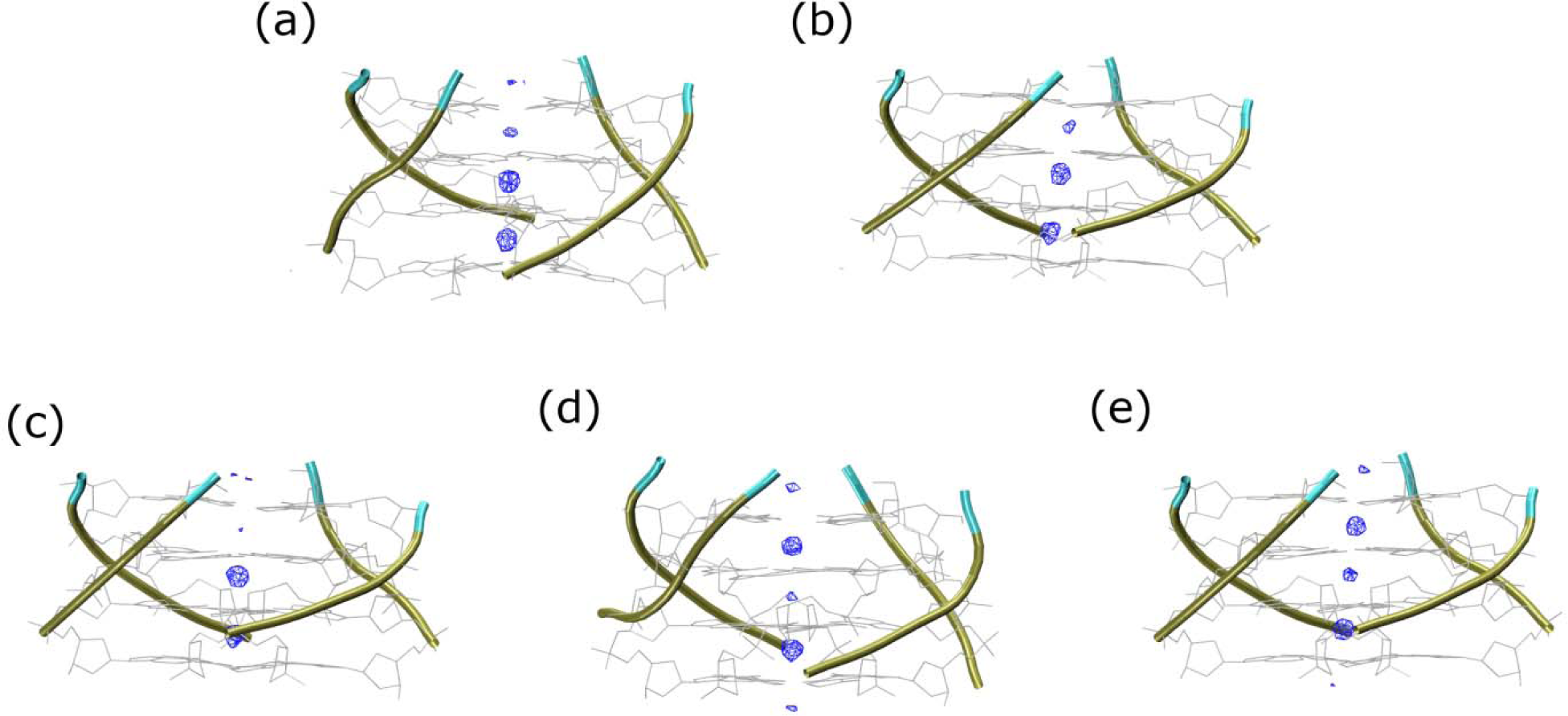
Cation binding sites (densities) observed in the K^+^/SPC/E Simulations (a) 23a, (b) 23b, (c) 23c, (d) 23d and (e) 23e of d[GGGG]_4_. The backbone of the G-stem is shown in tan tube and the 5′-end of the GQ backbone is shown in cyan. The guanine nucleotides are shown in silver lines and the K^+^ binding sites within the G-stem are shown in blue. Cation densities were developed above the G-stem due to the movement of cation from Site 1 to above the 5′-quartet. Cation densities were also developed below the G-stem as solvent cations were transiently trapped below the 3′-quartet. The figure shows that mostly two cations were present in the channel due to repulsion between the closely spaced cations. A 300 Å^-3^ threshold was used for density representation.

We also carried out five 1 μs long simulations of d[GGGG]_4_ in OPC with Na^+^ (Figure S33). The systems were visibly distorted in three out of five simulations as strand slippage events were observed. In one simulation the GQ was stable and no ion exchange event was observed while in the last simulation, four cation exchange events were observed but the stem was stable. The strand slippage events in this series of simulations were initiated by melting of the terminal quartet. One guanine left the quartet, stacked on the remaining G-triad, and it eventually enforced strand slippage, as the lone stacked guanine pulled the whole G-strand by one level (Figure S34). Substantially non-planar G-quartets were sampled from the onset of the simulations (because of the apparently imbalanced G(O6)-Na^+^ attraction resulting in buckling of Gs), but that was likely a prelude to the described strand slippage. Considering the short simulation time scale, the simulations suggest substantial volatility of the d[GGGG]_4_ (see Discussion).

In the 1 μs long simulations of d[GGGG]_4_ in K^+^ and OPC water model, the G-stem was stable in only two out of five simulations (Figure S35). In the Simulations 25a - 25c, Site 1 became vacant and a transient cation binding site over the first quartet was observed. The first (5′-end) quartet in these simulations was disrupted at 180 ns, 280 ns and 350 ns, respectively, by the movement of one or two guanines over the remaining bases of the quartet. The other three quartets of the G-stem did not fully disintegrate but exhibited substantial fluctuations. The cations at Sites 2 and 3 were retained in these simulations (Figure S36a - S36c). In Simulation 25d, the number of ions in the stem was reduced to two, Site 2 became vacant and additional transient cation binding sites were formed above and below the G-stem (Figure S36d). In Simulation 25e, the ion from Site 1 was pushed above the first quartet. Re-entry of this cation back into Site 1 pushed the cation from Site 3 below the G-stem and when the latter ion returned back to Site 3 the ion from Site 1 moved again above the stem (Figure S36e). Ion exchange was also observed at Site 3. These results indicate the difficulty in stable incorporation of three ions in the channel.

We also tested LM Na^+^ and K^+^ ions in the SPC/E water model in the Simulations 26a-26c for the d[GGGG]_4_ G-stem. The G-stem was stable in the Na^+^ Simulation 26a but disruption of a quartet was observed in one of the two simulations in K^+^ ions, (Simulation 26b, Figure S37a). The SPC/E adapted LM Na^+^ ions showed similar behaviour as the JC Na^+^ ions as three ions stabilized the G-stem and ∼20 ion exchange events were observed (Figure S38a). In the Simulation 26b with the SPC/E adapted LM K^+^ ions, ion from Site 3 was lost to the solvent at 90 ns and guanine from the last quartet moved towards the solvent, disrupting the quartet. The other three quartets were maintained albeit with substantial fluctuations. In the second simulation in SPC/E adapted LM K^+^ ions (Simulation 26c), ion from Site 1 was pushed over the first quartet until the movement of ion from Site 2 to Site 1 at 0.7 μs (Figure S38b). Cations at Sites 1 and 3 stabilized the G-stem and no cation was present in Site 2 for the remaining of the simulation time. Thus, the LM Na^+^ and K^+^ ions in Simulations 26a and 26c behaved similarly to JC ions.

In Simulation 26d with OPC water model and HFE Na^+^ ions, GQ was stable but multiple ion exchange events were observed (Figures S37b and S38c) and the fourth quartet was buckled. In the OPC water model and HFE K^+^ ions (Simulation 26e), 5′-quartet was disrupted at 1.7 μs (Figures S37b and S38d). Disruption of 5′-quartet was also observed in the Simulations 26f and 26g with OPC water model and IOD Na^+^ and K^+^ ions at 1.0 μs and 400 ns, respectively (Figure S37c).

### Additional simulations

We have carried out simulations of d[GGGG]_4_ using TIP4P-D water with JC and CHARMM22 Na^+^ and K^+^ ions (simulation series 27). The results are presented in Supporting Information including Figures S39 and S40.

We also ran simulations of two quartet parallel and antiparallel G-stems (Simulations 28a −28f) and RNA two-quartet dimer and monomer (Simulations 29a - 29f) using the Langevin thermostat. The GQ system behaved similarly as with the Berendsen thermostat, within the limits of sampling (Figures S41 - S44). Further details are presented in Supporting Information.

### Complex dynamics of water molecules inside the channel of four-quartet GQs (Simulation series 1-3, 8-10, 22-27)

The trajectories of both RNA and DNA four-quartet GQs showed that water molecules may enter the channel via its 5′ or 3′ end. These waters may then be pushed further into the channel by an incoming cation. Then they may leave the channel concurrently with another ion exchange through the channel ends. In the case of four-quartet GQs with discontinuous strands (2RQJ, 2N3M), we rarely observed that waters could escape from Site 2 (the middle one) into the groove. This process is facilitated by the incorporation of leaving water molecule into a G-quartet, which then hosts one water-mediated G:G base pair (Figure S45). Such a G-quartet perturbation may last for microseconds, but it apparently does not destabilize the GQ. In one case we saw the process in reverse, since a water molecule penetrated into the channel from the groove. In tetramolecular GQs with continuous strands (d[GGGG]_4_), water molecules could escape the channel via incorporation into a quartet, but only the terminal ones; we did not see escape through the middle quartets into the groove. The analysis again confirms that the ensembles of the simulated G-stem structures are not homogeneous^83^ and contain diverse long-living substates. Interaction of long-residency water molecules in G-triads participating in a G-stem core has been characterized by NMR and MD simulations.^86^ However, incorporation of water molecules into a G-quartet appears to be a new observation. It may lead to rare conformational substates of the G-stems which might affect their interactions with other molecules.

## DISCUSSION

### Potential roles of two-quartet GQs

GQs are often assumed to need three consecutive G-quartets to be stable. On the other hand, the formation of stable two-quartet DNA or RNA GQs would imply that GQs could be far more widespread in the genome.^87^ In addition, RNA is single-stranded and in principle could form GQ whenever the right conditions are met.^88-94^ It has been shown that RNA GQs formed by 5′-untranslated regions of mRNAs could modulate translation *in vitro*.^95-99^ The stability of two-quartet RNA GQs would imply that shorter RNA molecules such as microRNAs may form GQs under physiological conditions and modulate cellular functions.^100^ In addition, these less stable GQs could have interesting dynamic properties, as they can quickly fold and unfold, and sensitively respond to the changes in cellular conditions. Various two-quartet GQs could be transiently formed during folding of larger GQs. The current experimental data are not sufficient to unambiguously assess the intrinsic stability of two-quartet GQs, albeit they show that very stable two-quartet GQs can form if stabilized by additional structural features such as favourable loop alignments or adjacent G-triads. Two most known cases are the DNA Thrombin binding aptamer (TBA) stabilized by the loops and the two-quartet form of the DNA human telomeric GQ stabilized by a G-triad,^21, 27^ both being anti-parallel GQs. More recent two-quartet DNA GQs have also been discovered in anti-parallel conformation.^36, 101^

### Stability of two-quartet GQs as captured by MD simulations

We investigated the potential to form two-quartet GQs with minimal or no support by additional stabilizing elements. We have performed a series of standard 1-10 μs long atomistic explicit solvent MD simulations assessing the intrinsic structural stability of two-quartet DNA and RNA GQs starting from the folded state. Such computations aim to capture the unfolding process, *i*.*e*., the lifetime (1/k_off_) of a given GQ when folded, as described by the simulation force field. Although the computations do not address directly the thermodynamic stability, it is plausible to assume that species with too short lifetimes cannot significantly contribute to the folding landscape. Due to many force-field approximations discussed below, we assume that the simulations underestimate the lifetimes of the studied species, but should provide a fairly good estimate of their relative structural stabilities. Most of our efforts were devoted to the more elusive parallel-stranded all-*anti* arrangements.

Our simulations predict that *two-quartet parallel-stranded all-anti DNA G-stems are intrinsically very unstable*, and cannot be formed (except very transiently) without extensive support by additional structural elements (Simulations 4a-4h and 5a-5d, Figures 3, S5 and S7). Parallel-stranded G-stems with 5′-*syn* bases and antiparallel two-quartet DNA G-stem are evidently more stable (Simulations 6a-6d and 7a-7k, Figures 4, 5 and S9-S11). This is consistent with the fact that the solution structures of antiparallel two-quartet DNA GQs with alignments on both sides of the G-stems are known.^21^ Similarly, the two-quartet RNA GQ is predicted to be more stable than its parallel DNA two-quartet counterpart, which is consistent with experiments suggesting larger stability of RNA over DNA GQs (simulation series 11 −15, Figures 6 and S17-S21).^38, 88, 91, 102^ Mergny and colleagues studied the kinetics and thermodynamics of tetramolecular DNA and RNA quadruplexes and concluded that RNA quadruplexes are more stable than their DNA counterparts as a result of both faster association and slower dissociation.^34^ Note, however, that all these comparisons were based on tetramolecular structures involving at least three quartets. The simulations reveal the origin of short lifetimes of unsupported two-quartet all-*anti* DNA G-stems (Table 1). They are volatile due to the ability of unobstructed vertical movements (slippage) of the strands caused by common thermal fluctuations.^80, 81^ In contrast to longer stems, any strand slippage event within a two-quartet stem leads to a weak structure with only one quartet that is prone to immediate disintegration. In the case of antiparallel G-stems, mutual vertical movements of the strands are hindered by the mixture of *syn* and *anti* residues in the stem. Additionally, where relevant, the terminal *syn*-specific 5′-OH – G(N3) H-bond may also provide stability to the antiparallel G-stem. In the case of RNA GQs, we suggest that their larger rigidity due to the presence of the 2 □-OH hydroxyl group reduces their propensity to vertical strand slippage.

### Force-field approximations and their effect on G-stem simulations

The simulations are influenced by diverse force-field imbalances.^66^ Generally, outcomes of simulation studies of fully folded cation-stabilized GQ structures are often unaffected by these imbalances, due to the extraordinary stability and stiffness of GQs.^44^ Simulations with appropriate force fields starting from fully folded cation-stabilized GQs are usually entirely stable, as their lifetimes, even as described by the force field, extend far beyond the simulation time scale. In contrast, any larger perturbation of the quartets in simulations of GQ structures derived from experiments indicates that an inadequate force field has been used.^44, 103^ Still, even with appropriate force fields, subtle details of the structures (details of quartet′s H-bonds,^29, 78^ cation distribution,^29^ backbone substates^63^ and loop conformations^74, 75, 77^) may be sampled imperfectly.

Force-field imbalances may impact considerably more simulations of GQ folding intermediates^44, 82^ and other weak structures such as the two-quartet stems studied here. Such simulations may be already visibly affected by force-field approximations. Our above-presented analysis of different stabilities of two-quartet GQ stems has been made based on primary MD data with consideration of the known systematic errors in the force-field description. Thus, we suggest that our interpretation of the simulation data is justified. Let us now explicitly comment on the imbalances, explain how they influence the presented primary simulation results and what new insights were learnt from the simulations about force-field limitations.

i. Stability of the GG H-bonding is generally assumed to be underestimated^44^ with the presently- used non-bonded terms.^66^ This problem can be partly corrected, for example, by adding a specific term to stabilize the H-bonds.^104^ Note that it does not mean that the H-bonding is uniformly underestimated in MD simulations of nucleic acids; there are other H-bonds which are over-stabilized by the force field, especially in RNA.^66, 104^ No means to enhance the stability of GG base pairing was used in this study.
ii. There are many indications that the base stacking is often over-stabilized in simulations of nucleic acids with the presently-used non-bonded terms, though this effect is likely context-dependent and non-uniform (reviewed in ref.^66^). For GQs, some over-stabilization of G-quartet stacking may be helpful since it compensates for some other force-field deficiencies. Stability of H-bonding and stacking in nucleic acids simulations is determined not only by the solute-solute interactions but mainly by the balance with the water model.^105-107^
iii. The force fields are suspected to provide an inaccurate description of the propeller loops.^74, 76, 77^ They are unable to sample the experimentally-determined (if known unambiguously)^74^ propeller loop geometries in otherwise stable simulations of fully folded GQs and are assumed to excessively destabilize the parallel G-hairpins and G-triplexes which affects simulations of folding intermediates.^41, 82^ The exact origin and magnitude of this imbalance remains unknown.^74^ The description of lateral and diagonal loops appears to be more realistic.^75^
iv. With commonly used force-field parameters, the pair additive force field underestimates the strength of the O6-cation interaction and thus also direct cation binding to the G-quartets.^77, 108^ In addition, when using monovalent cations that have parameters optimized to reproduce solvation energies with current water models, the cations inside the G-stem channel appear as too large (having a too large radius or too steep onset of the van der Waals repulsion).^77, 108^ These inaccuracies, together with approximate description of the base-base H-bonds, probably contribute to common sampling of bifurcated GG H-bonds in the simulated G-stems.^29, 78^
v. When two or more cations are lining up in the G-stem, yet another and likely severe imbalance arises, which has been noticed only quite recently.^109^ It is manifested as an excessive inter-cation repulsion because the ions are modelled as having fixed +1 charges. In real GQs, the inter-cation repulsion is weakened due to a complex interplay of polarization effects between the ions and the G-quartets, which is strongly geometry-dependent.^109^ This problem thus cannot be fixed when using the formalism of pair-additive force fields.

Due to the latter two imbalances, we remain rather sceptical with respect to MD simulation studies aiming at quantitative (free-energy) description of ion binding inside GQs, including studies attempting to explain the Na^+^ *vs*. K^+^ stabilization difference. In addition, as shown here and elsewhere,^83^ binding of ions in GQs includes diverse local substates which have a dramatic impact on results of free-energy calculations using the approximate MM-PBSA (or similar) methods. These methods magnify the free-energy differences among the substates while the simulation time scale is not sufficient to provide their converged populations.^83^ Recent literature data suggest that polarizable force fields could substantially improve the description of ions in GQs,^103^ though they may still require careful calibration and it is not yet clear if they do not introduce other hitherto not identified problems. However, G-stems stabilized by cations are genuine systems for application of the polarizable force fields.^103, 110-113^ Well-calibrated polarizable force fields should be capable to simultaneously balance parameters for the bulk and bound ions as well as to correctly describe the inter-cation repulsion inside the GQs.

The above-listed imbalances explain why we suggest that the structural stability of the two-quartet GQs is underestimated by the present simulations. It is affected by under-stabilization of the ion-quartet interactions and the GG base pairs, which is only partly compensated for by the likely overestimated stacking. Further, the intra- and bimolecular parallel GQs may be destabilized by the description of the propeller loops.

Our simulations of the larger systems are further affected by the exaggerated description of the inter-cation repulsion. This can contribute to the observation that in the simulations of the 2N3M structure, Site 2 of ion is often not occupied (Table 1). This specific prediction should thus be taken with care.

### The force field does not capture the increase of stability upon increasing the number of G-quartets correctly

Consideration of the excessive inter-cation repulsion also explains the quite frequent and likely excessive exchanges and ion shifts seen in simulations of the d[GGGG]_4_ all parallel tetrameric GQ (Figure 7). It also explains why the ion exchanges are not attenuated in the present d[GGGG]_4_ simulations compared to the earlier simulations of d[GGG]_4_.^29^ While at first sight, one might expect a major stabilization/rigidification of the system when moving from three quartets to four quartets, the force-field imbalance of the inter-cation interaction is getting more severe for the latter system. *In other words, we suggest that the fact that we see more ion dynamics, exchanges and expulsions for the d[GGGG]*_*4*_ *than for d[GGG]*_*4*_ *is a simulation artefact*. Note that in some of our simulations we have even seen modest perturbations of the stems. It indicates that simulations with pair-additive force-fields do not properly capture the increase of stability of GQs upon the increase of the number of G-quartets, which could be several orders of magnitude per one added G-quartet.^33, 34, 114^ Although direct calculation of free-energy differences between GQs with different number of quartets are not realistic due to sampling limitations, even structural-dynamics simulations can be affected when smooth lining up of the multiple ions in the channel is disturbed, as evidenced above. Fortunately, there are indications that the ion dynamics is attenuated in simulations of complete GQs compared to simulations of just the G-stems.^29, 83^

We did not see any preference for cation exchanges through the 5′-end and 3′-end quartets. Previous MD studies also indicated that in all-*anti* parallel-stranded GQ, the energy barriers for cations escaping the G-stem is approximately equal for both the 5′- and 3′-end.^115^

### Simulations with more sophisticated water models may under-stabilize G-stems

The above-noted imbalances also provide clues about the lesser structural stability of the simulated systems when using more sophisticated four-site OPC (the Main text) and TIP4P-D (Supporting Information) water models. The OPC water model is becoming very popular in simulations of RNA molecules since it appears to weaken the base stacking and spurious H-bonds;^66, 104-106^ overstabilization of these interactions with simple water models is a result of the balance of solute-solvent and solvent-solvent interactions. TIP4P-D enhances dispersion interactions and it was developed to balance the populations of compacted and loosely ordered states of disordered proteins because the widely used three-site TIP3P model overstabilizes the compacted state.^54^ TIP4P-D has also been suggested to improve the conformational behaviour of RNA tetraloops.^57^ However, as shown in the present work, the OPC and TIP4P-D water models (at least with the used ion parameters) are suboptimal for GQ simulations. They can lead to destabilization of at least some GQ systems, probably due to the loss of fortunate compensation of errors present with some more simple water models such as SPC/E. It appears that finding a universal water model for nucleic acids simulations is a complex problem and that the SPC/E water model with JC ions is a reasonable albeit not ideal compromise for GQ simulations. It should be noted that we do not want to *a priori* disregard the more sophisticated water models from GQ simulations, as they can bring potential benefits in the description of loop stacking, ligand binding or some other aspects of the simulations. The instabilities reported in the present work for several systems may be attenuated for complete intramolecular GQs with three loops.

However, one should be aware that these water models may under-stabilize the G-stems and lead to occasional spurious perturbations of simulated GQ structures, especially in longer simulations. The success of these water models has been related to a reduction of excessive compaction of biomolecular chains mostly due to weakening of some H-bonding and stacking interactions.^54, 66, 104-106^ This property, however, can for some other systems lead to undesirable under-stabilization of native folds.

## CONCLUSIONS

Nearly 250 GQ structures are currently accessible in the PDB database. The majority of the crystallised GQs are parallel-stranded and have three quartets. Solution structures of antiparallel two-quartet DNA GQs have also been observed.^21, 36^ The antiparallel two-quartet DNA GQs have extensive loop alignments above and below the guanine quartets or other stabilizing factors which could provide stability to the GQ. However, it remains to be explored if independent parallel-stranded two-quartet DNA GQs are unstable *per se* or are yet to be discovered.

Diverse experimental evidence suggests larger stability of RNA over DNA GQs. The 2□ -hydroxyl group in ribose sugar allows for more interactions with the bases, cations and water molecules thereby increasing the stability of the RNA GQs.^14^ It also restricts the orientation of the base favouring the *anti*-orientation. Therefore, the majority of the RNA GQs have been observed in the parallel-stranded topology with all bases in the *anti*-orientation.

Based on the simulations we suggest that parallel-stranded all-*anti* two-quartet DNA G-stems are intrinsically very unstable due to the propensity of strand slippage, leading to very short life-times. Such stems can be stably formed only with the help of additional stabilizing interactions, such as dimerization or linking of two-quartet parallel-stranded DNA GQs.^45^ Nevertheless, it does not preclude the occurrence of parallel-stranded two-quartet stems in transitory ensembles^41^ during folding processes. Antiparallel DNA two-quartet stems, as well as RNA two-quartet stems, are structurally considerably more stable than the parallel DNA two-quartet stems, although even these species likely require additional stabilizing elements to achieve detectable thermodynamic stability.

On the methodological side, we identify a visible consequence of exaggerated inter-cation repulsion in the G-stems caused by the force-field approximation.^109^ This leads to an enhanced rate of cation exchanges and even expulsions of ions in the d[GGGG]_4_ parallel all-*anti* G-stem compared to the d[GGG]_4_. This has been an entirely unexpected and counterintuitive result, considering the experimentally known sharp (several orders of magnitude) increase of GQ stability and life-times with the number of G-quartets.^33, 34, 114^ Although there is no direct experimental evidence of ion dynamics in these constructs, we assume that on µs time scale the ion-exchange dynamics in d[GGGG]_4_ should be minimal, the stem should be fully occupied by ions and entirely stable. This suggests that the force fields do not capture the increase of G-quadruplex stability with the number of G-quartets. The simulations also show differences among the tested water and ion parameters, confirming an earlier suggestion^29^ that improved water models such as OPC may be non-optimal for simulations of GQs.

## ASSOCIATED CONTENT

### Supporting Information

Methods for visualization of cation binding sites, detailed results of Simulations 4a - 4h, simulation series 8 - 10, simulation series 16 - 18 and simulation series 27 - 29 are presented in the Supporting Information. Tables S1 - S2 and Figures S1 - S45 supporting the results are also presented in the Supporting Information. PDB files associated with simulation series 4 are provided in the Supporting Information. The following files are available free of charge Supporting_Information.pdf.

## AUTHOR INFORMATION

### Author Contributions

The manuscript was written through the contributions of all authors. All authors have given approval to the final version of the manuscript.

## ACKNOWLEDGEMENT

B.I. and P.S. greatly appreciate access to storage facilities owned by parties and projects contributing to the National Grid Infrastructure MetaCentrum, provided under the programme “Projects of Large Research, Development, and Innovations Infrastructures” (CESNET LM2015042).

For Table of Contents Use Only

## Synopsis

We have carried out extended unbiased atomistic simulations using different ion and water parameters to study the stability of two and four quartet G-quadruplexes (GQs). The all- *anti* parallel-stranded two-quartet DNA GQs are highly unstable compared to their RNA counterpart in all the ion and water conditions. GQs composed of two stacked units of two-quartet GQs were, however, stable for both DNA and RNA.

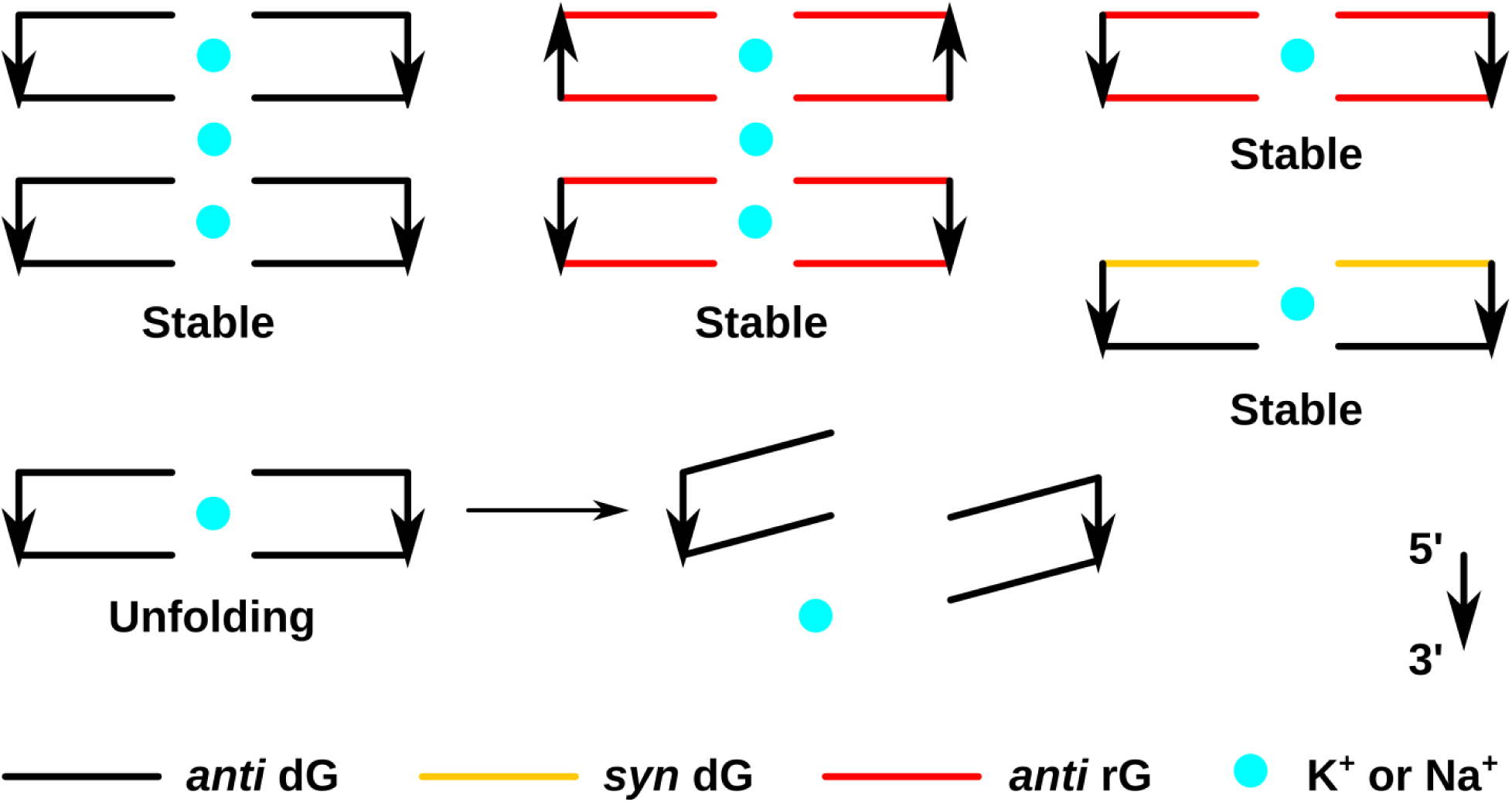

## REFERENCES

1. Bochman, M. L.; Paeschke, K.; Zakian, V. A., DNA secondary structures: stability and function of G-quadruplex structures. Nat. Rev. Genet. 2012, 13, 770–780.

2. Gilbert, D. E.; Feigon, J., Multistranded DNA structures. Curr. Opin. Struct. Biol. 1999, 9, 305–314.

3. Burge, S.; Parkinson, G. N.; Hazel, P.; Todd, A. K.; Neidle, S., Quadruplex DNA: sequence, topology and structure. Nucleic Acids Res. 2006, 34, 5402–5415.

4. Sen, D.; Gilbert, W., Formation of parallel four-stranded complexes by guanine-rich motifs in DNA and its implications for meiosis. Nature 1988, 334, 364–366.

5. Mao, S. Q.; Ghanbarian, A. T.; Spiegel, J.; Martinez Cuesta, S.; Beraldi, D.; Di Antonio, M.; Marsico, G.; Hansel-Hertsch, R.; Tannahill, D.; Balasubramanian, S., DNA G-quadruplex structures mold the DNA methylome. Nat. Struct. Mol. Biol. 2018, 25, 951–957.

6. Guilbaud, G.; Murat, P.; Recolin, B.; Campbell, B. C.; Maiter, A.; Sale, J. E.; Balasubramanian, S., Local epigenetic reprogramming induced by G-quadruplex ligands. Nat. Chem. 2017, 9, 1110–1117.

7. Gellert, M.; Lipsett, M. N.; Davies, D. R., Helix formation by guanylic acid. Proc. Natl. Acad. Sci. U. S. A. 1962, 48, 2013–2018.

8. Parkinson, G. N.; Lee, M. P.; Neidle, S., Crystal structure of parallel quadruplexes from human telomeric DNA. Nature 2002, 417, 876–880.

9. Sket, P.; Plavec, J., Tetramolecular DNA quadruplexes in solution: insights into structural diversity and cation movement. J. Am. Chem. Soc. 2010, 132, 12724–12732.

10. Webba da Silva, M., Geometric formalism for DNA quadruplex folding. Chemistry 2007, 13, 9738–9745.

11. Phan, A. T., Human telomeric G-quadruplex: structures of DNA and RNA sequences. FEBS J. 2010, 277, 1107–1117.

12. Cang, X.; Sponer, J.; Cheatham, T. E., 3rd, Explaining the varied glycosidic conformational, G-tract length and sequence preferences for anti-parallel G-quadruplexes. Nucleic Acids Res. 2011, 39, 4499–4512.

13. Dvorkin, S. A.; Karsisiotis, A. I.; Webba da Silva, M., Encoding canonical DNA quadruplex structure. Sci. Adv. 2018, 4, eaat3007.

14. Sacca, B.; Lacroix, L.; Mergny, J. L., The effect of chemical modifications on the thermal stability of different G-quadruplex-forming oligonucleotides. Nucleic Acids Res. 2005, 33, 1182–1192.

15. Xu, Y.; Kaminaga, K.; Komiyama, M., G-quadruplex formation by human telomeric repeats-containing RNA in Na+ solution. J. Am. Chem. Soc. 2008, 130, 11179–11184.

16. Martadinata, H.; Phan, A. T., Structure of propeller-type parallel-stranded RNA G-quadruplexes, formed by human telomeric RNA sequences in K+ solution. Journal of the American Chemical Society 2009, 131, 2570–2578.

17. Martadinata, H.; Phan, A. T., Structure of human telomeric RNA (TERRA): stacking of two G-quadruplex blocks in K(+) solution. Biochemistry 2013, 52, 2176–2183.

18. Collie, G. W.; Haider, S. M.; Neidle, S.; Parkinson, G. N., A crystallographic and modelling study of a human telomeric RNA (TERRA) quadruplex. Nucleic Acids Res. 2010, 38, 5569–5580.

19. Sweeney, B. A.; Roy, P.; Leontis, N. B., An introduction to recurrent nucleotide interactions in RNA. Wiley Interdiscip. Rev. RNA 2015, 6, 17–45.

20. Dai, J.; Punchihewa, C.; Ambrus, A.; Chen, D.; Jones, R. A.; Yang, D., Structure of the intramolecular human telomeric G-quadruplex in potassium solution: a novel adenine triple formation. Nucleic Acids Res. 2007, 35, 2440–2450.

21. Schultze, P.; Macaya, R. F.; Feigon, J., Three-dimensional solution structure of the thrombin-binding DNA aptamer d(GGTTGGTGTGGTTGG). J. Mol. Biol. 1994, 235, 1532–47.

22. Wang, Y.; Patel, D. J., Solution structure of the human telomeric repeat d[AG3(T2AG3)3] G-tetraplex. Structure 1993, 1, 263–282.

23. Lim, K. W.; Ng, V. C.; Martin-Pintado, N.; Heddi, B.; Phan, A. T., Structure of the human telomere in Na+ solution: an antiparallel (2+2) G-quadruplex scaffold reveals additional diversity. Nucleic Acids Res. 2013, 41, 10556–10562.

24. Dai, J.; Carver, M.; Punchihewa, C.; Jones, R. A.; Yang, D., Structure of the Hybrid-2 type intramolecular human telomeric G-quadruplex in K+ solution: insights into structure polymorphism of the human telomeric sequence. Nucleic Acids Res. 2007, 35, 4927–4940.

25. Luu, K. N.; Phan, A. T.; Kuryavyi, V.; Lacroix, L.; Patel, D. J., Structure of the human telomere in K+ solution: an intramolecular (3 + 1) G-quadruplex scaffold. J. Am. Chem. Soc. 2006, 128, 9963–9970.

26. Phan, A. T.; Kuryavyi, V.; Luu, K. N.; Patel, D. J., Structure of two intramolecular G-quadruplexes formed by natural human telomere sequences in K+ solution. Nucleic Acids Res. 2007, 35, 6517–25.

27. Lim, K. W.; Amrane, S.; Bouaziz, S.; Xu, W.; Mu, Y.; Patel, D. J.; Luu, K. N.; Phan, A. T., Structure of the human telomere in K+ solution: a stable basket-type G-quadruplex with only two G-tetrad layers. J. Am. Chem. Soc. 2009, 131, 4301–4309.

28. Schultze, P.; Hud, N. V.; Smith, F. W.; Feigon, J., The effect of sodium, potassium and ammonium ions on the conformation of the dimeric quadruplex formed by the Oxytricha nova telomere repeat oligonucleotide d(G(4)T(4)G(4)). Nucleic Acids Res. 1999, 27, 3018–3028.

29. Havrila, M.; Stadlbauer, P.; Islam, B.; Otyepka, M.; Šponer, J., Effect of Monovalent Ion Parameters on Molecular Dynamics Simulations of G-Quadruplexes. J. Chem. Theory Comput. 2017, 13, 3911–3926.

30. Marchand, A.; Gabelica, V., Folding and misfolding pathways of G-quadruplex DNA. Nucleic Acids Res. 2016, 44, 10999–11012.

31. Hardin, C. C.; Henderson, E.; Watson, T.; Prosser, J. K., Monovalent cation induced structural transitions in telomeric DNAs: G-DNA folding intermediates. Biochemistry 1991, 30, 4460–4472.

32. Payet, L.; Huppert, J. L., Stability and structure of long intramolecular G-quadruplexes. Biochemistry 2012, 51, 3154–3161.

33. Bardin, C.; Leroy, J. L., The formation pathway of tetramolecular G-quadruplexes. Nucleic Acids Res. 2008, 36, 477–488.

34. Mergny, J.-L.; De Cian, A.; Ghelab, A.; Saccà, B.; Lacroix, L., Kinetics of tetramolecular quadruplexes. Nucleic Acids Res. 2005, 33, 81–94.

35. Lee, J. Y.; Yoon, J.; Kihm, H. W.; Kim, D. S., Structural diversity and extreme stability of unimolecular Oxytricha nova telomeric G-quadruplex. Biochemistry 2008, 47, 3389–3396.

36. Lenarčič, Ž. M.; Rozman, J.; Plavec, J., Adenine-Driven Structural Switch from a Two-to Three-Quartet DNA G-Quadruplex. Angew. Chem., Int. Ed. 2018, 57, 15395–15399.

37. Mashima, T.; Matsugami, A.; Nishikawa, F.; Nishikawa, S.; Katahira, M., Unique quadruplex structure and interaction of an RNA aptamer against bovine prion protein. Nucleic Acids Res. 2009, 37, 6249–6258.

38. Arora, A.; Maiti, S., Differential biophysical behavior of human telomeric RNA and DNA quadruplex. J. Phys. Chem. B 2009, 113, 10515–10520.

39. Kejnovska, I.; Bednarova, K.; Renciuk, D.; Dvorakova, Z.; Skolakova, P.; Trantirek, L.; Fiala, R.; Vorlickova, M.; Sagi, J., Clustered abasic lesions profoundly change the structure and stability of human telomeric G-quadruplexes. Nucleic Acids Res. 2017, 45, 4294–4305.

40. Bednářová, K.; Kejnovská, I.; Vorlíčková, M.; Renčiuk, D., Guanine Substitutions Prevent Conformational Switch from Antiparallel to Parallel G-Quadruplex. Chem.: Eur. J. 2019, 25, 13422–13428.

41. Stadlbauer, P.; Kührová, P.; Vicherek, L.; Banáš, P.; Otyepka, M.; Trantírek, L.; Šponer, J., Parallel G-triplexes and G-hairpins as potential transitory ensembles in the folding of parallel-stranded DNA G-Quadruplexes. Nucleic Acids Res. 2019, 47, 7276–7293.

42. Qin, M.; Chen, Z.; Luo, Q.; Wen, Y.; Zhang, N.; Jiang, H.; .uaiyu, Two-Quartet G-Quadruplexes Formed by DNA Sequences Containing Four Contiguous GG Runs. J. Phys. Chem. B 2015, 119, 3706–3713.

43. Zahin, M.; Dean, W. L.; Ghim, S.-J.; Joh, J.; Gray, R. D.; Khanal, S.; Bossart, G. D.; Mignucci-Giannoni, A. A.; Rouchka, E. C.; Jenson, A. B.; Trent, J. O.; Chaires, J. B.; Chariker, J. H., Identification of G-quadruplex forming sequences in three manatee papillomaviruses. PLOS ONE 2018, 13, e0195625.

44. Šponer, J.; Bussi, G.; Stadlbauer, P.; Kührová, P.; Banáš, P.; Islam, B.; Haider, S.; Neidle, S.; Otyepka, M., Folding of guanine quadruplex molecules–funnel-like mechanism or kinetic partitioning? An overview from MD simulation studies. Biochim. Biophys. Acta 2017, 1861, 1246–1263.

45. Do, N. Q.; Chung, W. J.; Truong, T. H. A.; Heddi, B.; Phan, A. T., G-quadruplex structure of an anti-proliferative DNA sequence. Nucleic Acids Res. 2017, 45, 7487–7493.

46. Clark, G. R.; Pytel, P. D.; Squire, C. J., The high-resolution crystal structure of a parallel intermolecular DNA G-4 quadruplex/drug complex employing syn glycosyl linkages. Nucleic Acids Res. 2012, 40, 5731–5738.

47. Mandal, P. K.; Collie, G. W.; Kauffmann, B.; Huc, I., Racemic DNA crystallography. Angew. Chem. Int. Ed. Engl. 2014, 53, 14424–14427.

48. Laughlan, G.; Murchie, A. I.; Norman, D. G.; Moore, M. H.; Moody, P. C.; Lilley, D. M.; Luisi, B., The high-resolution crystal structure of a parallel-stranded guanine tetraplex. Science 1994, 265, 520–524.

49. Zgarbová, M.; Otyepka, M.; Šponer, J.; Lankaš, F.; Jurečka, P., Base Pair Fraying in Molecular Dynamics Simulations of DNA and RNA. J. Chem. Theory Comput. 2014, 10, 3177–3189.

50. Galindo-Murillo, R.; Roe, D. R.; Cheatham, T. E., Convergence and reproducibility in molecular dynamics simulations of the DNA duplex d(GCACGAACGAACGAACGC). BBA Gen. Subjects 2015, 1850, 1041–1058.

51. Romanucci, V.; Marchand, A.; Mendoza, O.; D’Alonzo, D.; Zarrelli, A.; Gabelica, V.; Di Fabio, G., Kinetic ESI-MS Studies of Potent Anti-HIV Aptamers Based on the G-Quadruplex Forming Sequence d(TGGGAG). ACS Med. Chem. Letts. 2016, 7, 256–260.

52. Berendsen, H. J. C.; Grigera, J. R.; Straatsma, T. P., The missing term in effective pair potentials. J. Phys. Chem. 1987, 91, 6269–6271.

53. Izadi, S.; Anandakrishnan, R.; Onufriev, A. V., Building Water Models: A Different Approach. J. Phys. Chem. Lett. 2014, 5, 3863–3871.

54. Piana, S.; Donchev, A. G.; Robustelli, P.; Shaw, D. E., Water Dispersion Interactions Strongly Influence Simulated Structural Properties of Disordered Protein States. J. Phys. Chem. A 2015, 119, 5113–5123.

55. Joung, I. S.; Cheatham, T. E., 3rd, Determination of alkali and halide monovalent ion parameters for use in explicitly solvated biomolecular simulations. J. Phys. Chem. B 2008, 112, 9020–9041.

56. MacKerell, A. D.; Bashford, D.; Bellott, M.; Dunbrack, R. L.; Evanseck, J. D.; Field, M. J.; Fischer, S.; Gao, J.; Guo, H.; Ha, S.; Joseph-McCarthy, D.; Kuchnir, L.; Kuczera, K.; Lau, F. T. K.; Mattos, C.; Michnick, S.; Ngo, T.; Nguyen, D. T.; Prodhom, B.; Reiher, W. E.; Roux, B.; Schlenkrich, M.; Smith, J. C.; Stote, R.; Straub, J.; Watanabe, M.; Wiórkiewicz-Kuczera, J.; Yin, D.; Karplus, M., All-Atom Empirical Potential for Molecular Modeling and Dynamics Studies of Proteins. J. Phys. Chem. B 1998, 102, 3586–3616.

57. Tan, D.; Piana, S.; Dirks, R. M.; Shaw, D. E., RNA force field with accuracy comparable to state-of-the-art protein force fields. Proc. Natl. Acad. Sci. U. S. A. 2018, 115, E1346–E1355.

58. Li, P.; Song, L. F.; Merz, K. M., Systematic Parameterization of Monovalent Ions Employing the Nonbonded Model. J. Chem. Theory Comput. 2015, 11, 1645–1657.

59. Case, D.; Betz, R.; Cerutti, D. S.; Cheatham, T. E., 3rd; Darden, T.; Duke, R.; Giese, T. J.; Gohlke, H.; Götz, A.; Homeyer, N.; Izadi, S.; Janowski, P.; Kaus, J.; Kovalenko, A.; Lee, T.-S.; LeGrand, S.; Li, P.; Lin, C.; Luchko, T.; A. Kollman, P., Amber 16, University of California, San Francisco. 2016.

60. Zgarbova, M.; Sponer, J.; Otyepka, M.; Cheatham, T. E., 3rd; Galindo-Murillo, R.; Jurecka, P., Refinement of the Sugar-Phosphate Backbone Torsion Beta for AMBER Force Fields Improves the Description of Z- and B-DNA. J. Chem. Theory Comput. 2015, 11, 5723–5736.

61. Zgarbová, M.; Otyepka, M.; Sponer, J.; Mládek, A.; Banáš, P.; Cheatham, T. E., 3rd; Jurečka, P., Refinement of the Cornell et al. Nucleic Acids Force Field Based on Reference Quantum Chemical Calculations of Glycosidic Torsion Profiles. J. Chem. Theory Comput. 2011, 7, 2886–2902.

62. Perez, A.; Marchan, I.; Svozil, D.; Sponer, J.; Cheatham, T. E., 3rd; Laughton, C. A.; Orozco, M., Refinement of the AMBER force field for nucleic acids: improving the description of alpha/gamma conformers. Biophysical journal 2007, 92, 3817–3829.

63. Krepl, M.; Zgarbova, M.; Stadlbauer, P.; Otyepka, M.; Banas, P.; Koca, J.; Cheatham, T. E., 3rd; Jurecka, P.; Sponer, J., Reference simulations of noncanonical nucleic acids with different chi variants of the AMBER force field: quadruplex DNA, quadruplex RNA and Z-DNA. J. Chem. Theory Comput. 2012, 8, 2506–2520.

64. Zgarbova, M.; Luque, F. J.; Sponer, J.; Cheatham, T. E., 3rd; Otyepka, M.; Jurecka, P., Toward Improved Description of DNA Backbone: Revisiting Epsilon and Zeta Torsion Force Field Parameters. J. Chem. Theory Comput. 2013, 9, 2339–2354.

65. Cornell, W. D.; Cieplak, P.; Bayly, C. I.; Gould, I. R.; Merz, K. M.; Ferguson, D. M.; Spellmeyer, D. C.; Fox, T.; Caldwell, J. W.; Kollman, P. A., A Second Generation Force Field for the Simulation of Proteins, Nucleic Acids, and Organic Molecules. J. Am. Chem. Soc. 1995, 117, 5179–5197.

66. Sponer, J.; Bussi, G.; Krepl, M.; Banas, P.; Bottaro, S.; Cunha, R. A.; Gil-Ley, A.; Pinamonti, G.; Poblete, S.; Jurecka, P.; Walter, N. G.; Otyepka, M., RNA Structural Dynamics As Captured by Molecular Simulations: A Comprehensive Overview. Chem. Rev. 2018, 118, 4177–4338.

67. Darden, T.; York, D.; Pedersen, L., Particle mesh Ewald: An N·log(N) method for Ewald sums in large systems. J. Chem. Phys. 1993, 98, 10089–10092.

68. Ryckaert, J.-P.; Ciccotti, G.; Berendsen, H. J. C., Numerical integration of the cartesian equations of motion of a system with constraints: molecular dynamics of n-alkanes. J. Comput. Phys. 1977, 23, 327–341.

69. Hopkins, C. W.; Le Grand, S.; Walker, R. C.; Roitberg, A. E., Long-Time-Step Molecular Dynamics through Hydrogen Mass Repartitioning. J. Chem. Theory Comput. 2015, 11, 1864–1874.

70. Berendsen, H. J. C.; Postma, J. P. M.; van Gunsteren, W. F.; DiNola, A.; Haak, J. R., Molecular dynamics with coupling to an external bath. J. Chem. Phys. 1984, 81, 3684–3690.

71. Roe, D. R.; Cheatham, T. E., 3rd, PTRAJ and CPPTRAJ: Software for Processing and Analysis of Molecular Dynamics Trajectory Data. J. Chem. Theory Comput. 2013, 9, 3084–3095.

72. Humphrey, W.; Dalke, A.; Schulten, K., VMD: Visual molecular dynamics. J. Mol. Graph. 1996, 14, 33–38.

73. Akhshi, P.; Wu, G., Umbrella sampling molecular dynamics simulations reveal concerted ion movement through G-quadruplex DNA channels. Phys. Chem. Chem. Phys. 2017, 19, 11017–11025.

74. Islam, B.; Stadlbauer, P.; Gil-Ley, A.; Pérez-Hernández, G.; Haider, S.; Neidle, S.; Bussi, G.; Banas, P.; Otyepka, M.; Sponer, J., Exploring the Dynamics of Propeller Loops in Human Telomeric DNA Quadruplexes Using Atomistic Simulations. J. Chem. Theory Comput. 2017, 13, 2458–2480.

75. Islam, B.; Stadlbauer, P.; Krepl, M.; Havrila, M.; Haider, S.; Sponer, J., Structural Dynamics of Lateral and Diagonal Loops of Human Telomeric G-Quadruplexes in Extended MD Simulations. J. Chem. Theory Comput. 2018, 14, 5011–5026.

76. Islam, B.; Stadlbauer, P.; Krepl, M.; Koca, J.; Neidle, S.; Haider, S.; Sponer, J., Extended molecular dynamics of a c-kit promoter quadruplex. Nucleic Acids Res. 2015, 43, 8673–8693.

77. Fadrna, E.; Spackova, N.; Sarzynska, J.; Koca, J.; Orozco, M.; Cheatham, T. E., 3rd; Kulinski, T.; Sponer, J., Single Stranded Loops of Quadruplex DNA As Key Benchmark for Testing Nucleic Acids Force Fields. J. Chem. Theory Comput. 2009, 5, 2514–2530.

78. Špačková, N.; Berger, I.; Šponer, J., Nanosecond Molecular Dynamics Simulations of Parallel and Antiparallel Guanine Quadruplex DNA Molecules. J. Am. Chem. Soc. 1999, 121, 5519–5534.

79. Akhshi, P.; Mosey, N. J.; Wu, G., Free-Energy Landscapes of Ion Movement through a G-Quadruplex DNA Channel. Angew. Chem., Int. Ed. 2012, 51, 2850–2854.

80. Stadlbauer, P.; Krepl, M.; Cheatham, T. E., 3rd; Koča, J.; Šponer, J., Structural dynamics of possible late-stage intermediates in folding of quadruplex DNA studied by molecular simulations. Nucleic Acids Res. 2013, 41, 7128–7143.

81. Stefl, R.; Cheatham, T. E., 3rd; Spacková, N.; Fadrná, E.; Berger, I.; Koca, J.; Sponer, J., Formation pathways of a guanine-quadruplex DNA revealed by molecular dynamics and thermodynamic analysis of the substates. Biophysical journal 2003, 85, 1787–1804.

82. Havrila, M.; Stadlbauer, P.; Kührová, P.; Banáš, P.; Mergny, J.-L.; Otyepka, M.; Šponer, J., Structural dynamics of propeller loop: towards folding of RNA G-quadruplex. Nucleic Acids Res. 2018, 46, 8754–8771.

83. Islam, B.; Stadlbauer, P.; Neidle, S.; Haider, S.; Sponer, J., Can We Execute Reliable MM-PBSA Free Energy Computations of Relative Stabilities of Different Guanine Quadruplex Folds? J. Phys. Chem. B 2016, 120, 2899–2912.

84. Sponer, J.; Mladek, A.; Spackova, N.; Cang, X.; Cheatham, T. E., 3rd; Grimme, S., Relative stability of different DNA guanine quadruplex stem topologies derived using large-scale quantum-chemical computations. J. Am. Chem. Soc. 2013, 135, 9785–9796.

85. Largy, E.; Mergny, J. L.; Gabelica, V., Role of Alkali Metal Ions in G-Quadruplex Nucleic Acid Structure and Stability. Met. Ions Life Sci. 2016, 16, 203–258.

86. Heddi, B.; Martín-Pintado, N.; Serimbetov, Z.; Kari, T. M. A.; Phan, A. T., G-quadruplexes with (4n - 1) guanines in the G-tetrad core: formation of a G-triad·water complex and implication for small-molecule binding. Nucleic Acids Res. 2015, 44, 910–916.

87. Kwok, C. K.; Marsico, G.; Sahakyan, A. B.; Chambers, V. S.; Balasubramanian, S., rG4-seq reveals widespread formation of G-quadruplex structures in the human transcriptome. Nature Methods 2016, 13, 841.

88. Fay, M. M.; Lyons, S. M.; Ivanov, P., RNA G-Quadruplexes in Biology: Principles and Molecular Mechanisms. J. Mol. Biol. 2017, 429, 2127–2147.

89. Malgowska, M.; Czajczynska, K.; Gudanis, D.; Tworak, A.; Gdaniec, Z., Overview of the RNA G-quadruplex structures. Acta Biochim. Pol. 2016, 63, 609–621.

90. Ji, X.; Sun, H.; Zhou, H.; Xiang, J.; Tang, Y.; Zhao, C., Research Progress of RNA Quadruplex. Nucleic Acid Ther. 2011, 21, 185–200.

91. Zhang, D. H.; Fujimoto, T.; Saxena, S.; Yu, H. Q.; Miyoshi, D.; Sugimoto, N., Monomorphic RNA G-quadruplex and polymorphic DNA G-quadruplex structures responding to cellular environmental factors. Biochemistry 2010, 49, 4554–4563.

92. Pandey, S.; Agarwala, P.; Maiti, S., Effect of Loops and G-Quartets on the Stability of RNA G-Quadruplexes. J. Phys. Chem. B 2013, 117, 6896–6905.

93. Biffi, G.; Di Antonio, M.; Tannahill, D.; Balasubramanian, S., Visualization and selective chemical targeting of RNA G-quadruplex structures in the cytoplasm of human cells. Nature chemistry 2014, 6, 75–80.

94. Malgowska, M.; Gudanis, D.; Kierzek, R.; Wyszko, E.; Gabelica, V.; Gdaniec, Z., Distinctive structural motifs of RNA G-quadruplexes composed of AGG, CGG and UGG trinucleotide repeats. Nucleic Acids Res. 2014, 42, 10196–10207.

95. Kumari, S.; Bugaut, A.; Huppert, J. L.; Balasubramanian, S., An RNA G-quadruplex in the 5’ UTR of the NRAS proto-oncogene modulates translation. Nature Chem. Biol. 2007, 3, 218–221.

96. Murat, P.; Marsico, G.; Herdy, B.; Ghanbarian, A. T.; Portella, G.; Balasubramanian, S., RNA G-quadruplexes at upstream open reading frames cause DHX36- and DHX9-dependent translation of human mRNAs. Genome Biol. 2018, 19, 229.

97. Wolfe, A. L.; Singh, K.; Zhong, Y.; Drewe, P.; Rajasekhar, V. K.; Sanghvi, V. R.; Mavrakis, K. J.; Jiang, M.; Roderick, J. E.; Van der Meulen, J.; Schatz, J. H.; Rodrigo, C. M.; Zhao, C.; Rondou, P.; de Stanchina, E.; Teruya-Feldstein, J.; Kelliher, M. A.; Speleman, F.; Porco, J. A., Jr.; Pelletier, J.; Rätsch, G.; Wendel, H.-G., RNA G-quadruplexes cause eIF4A-dependent oncogene translation in cancer. Nature 2014, 513, 65–70.

98. Patel, D. J.; Phan, A. T.; Kuryavyi, V., Human telomere, oncogenic promoter and 5’-UTR G-quadruplexes: diverse higher order DNA and RNA targets for cancer therapeutics. Nucleic Acids Res. 2007, 35, 7429–7455.

99. Beaudoin, J.-D.; Perreault, J.-P., 5’-UTR G-quadruplex structures acting as translational repressors. Nucleic Acids Res. 2010, 38, 7022–7036.

100. Chan, K. L.; Peng, B.; Umar, M. I.; Chan, C.-Y.; Sahakyan, A. B.; Le, M. T. N.; Kwok, C. K., Structural analysis reveals the formation and role of RNA G-quadruplex structures in human mature microRNAs. ChemComm 2018, 54, 10878–10881.

101. Kotar, A.; Rigo, R.; Sissi, C.; Plavec, J., Two-quartet kit* G-quadruplex is formed via double-stranded pre-folded structure. Nucleic acids research 2018, 47, 2641–2653.

102. Kankia, B., Stability Factors of the Parallel Quadruplexes: DNA Versus RNA. J. Phys. Chem. B 2019, 123, 1060–1067.

103. Salsbury, A. M.; Lemkul, J. A., Molecular Dynamics Simulations of the c-kit1 Promoter G-Quadruplex: Importance of Electronic Polarization on Stability and Cooperative Ion Binding. J. Phys. Chem. B 2019, 123, 148–159.

104. Kuhrova, P.; Mlynsky, V.; Zgarbova, M.; Krepl, M.; Bussi, G.; Best, R. B.; Otyepka, M.; Sponer, J.; Banas, P., Improving the Performance of the Amber RNA Force Field by Tuning the Hydrogen-Bonding Interactions. J. Chem. Theory Comput. 2019, 15, 3288–3305.

105. Bergonzo, C.; Cheatham, T. E., 3rd, Improved Force Field Parameters Lead to a Better Description of RNA Structure. J. Chem. Theory Comput. 2015, 11, 3969–3972.

106. Bergonzo, C.; Henriksen, N. M.; Roe, D. R.; Cheatham, T. E., 3rd, Highly sampled tetranucleotide and tetraloop motifs enable evaluation of common RNA force fields. Rna 2015, 21, 1578–1590.

107. Hase, F.; Zacharias, M., Free energy analysis and mechanism of base pair stacking in nicked DNA. Nucleic Acids Res. 2016, 44, 7100–7108.

108. Sponer, J.; Spackova, N., Molecular dynamics simulations and their application to four-stranded DNA. Methods 2007, 43, 278–90.

109. Gkionis, K.; Kruse, H.; Platts, J. A.; Mládek, A.; Koča, J.; Šponer, J., Ion Binding to Quadruplex DNA Stems. Comparison of MM and QM Descriptions Reveals Sizable Polarization Effects Not Included in Contemporary Simulations. J. Chem. Theory Comput. 2014, 10, 1326–1340.

110. Lemkul, J. A., Same fold, different properties: polarizable molecular dynamics simulations of telomeric and TERRA G-quadruplexes. Nucleic Acids Res. 2020, 48, 561–575.

111. Lemkul, J. A.; MacKerell, A. D., Polarizable Force Field for DNA Based on the Classical Drude Oscillator: I. Refinement Using Quantum Mechanical Base Stacking and Conformational Energetics. J. Chem. Theory Comput. 2017, 13, 2053–2071.

112. Jing, Z.; Qi, R.; Thibonnier, M.; Ren, P., Molecular Dynamics Study of the Hybridization between RNA and Modified Oligonucleotides. J. Chem. Theory Comput. 2019, 15, 6422–6432.

113. Zhang, C.; Lu, C.; Jing, Z.; Wu, C.; Piquemal, J.-P.; Ponder, J. W.; Ren, P., AMOEBA Polarizable Atomic Multipole Force Field for Nucleic Acids. J. Chem. Theory Comput. 2018, 14, 2084–2108.

114. Gros, J.; Rosu, F.; Amrane, S.; De Cian, A.; Gabelica, V.; Lacroix, L.; Mergny, J.-L., Guanines are a quartet’s best friend: impact of base substitutions on the kinetics and stability of tetramolecular quadruplexes. Nucleic Acids Res. 2007, 35, 3064–3075.

115. Zhu, H.; Xiao, S.; Wang, L.; Liang, H., Communication: Asymmetrical cation movements through G-quadruplex DNA. J. Chem. Phys. 2014, 141, 041103.

