## Supporting_Information for "Stability of Two-quartet G-quadruplexes and Their Dimers in Atomistic Simulations"

### *Supporting Methods*

#### Visualization of cation binding sites

The distance of all the cations to the guanine O6 atoms was calculated using ptraj module of Ambertools. For each quartet in a given GQ, we identified which of the four O6 atoms within a single quartet was closest to the particular cation and which O6 atom was farthest. In the following description, whenever we mention the distance of quartet X to a cation, we mean the shortest cation-O6 distance of the four guanines within the quartet X, unless explicitly stated otherwise. By comparison of these distances among all quartets in the system, we determined the cation binding site as follows:

1) For a two-quartet system, if the distance of a given cation to both quartets was shorter than 4 Å, the cation resided in Site 1 (between the two quartets). If the cation distance from one quartet was longer than 4 Å, but the longest cation-O6 distance of the other quartet was shorter than 4 Å, the cation resided either in Site 0 or 2 (that is either above or below the GQ) depending on the proximity to the 5′ or the 3′-quartet, respectively. Anything else was considered unbound.
2) For a three-quartet system, occupancies of Sites 0 and 3 (above or below the GQ) were determined as for the two-quartet system. Else if the cation was not farther than 4.5 Å from any of the O6 atoms of the middle quartet and the cation distance was shorter to the 5′-quartet than to the 3′-quartet, the cation resided in Site 1; if it was closer to the 3′-quartet than to the 5′-quartet, it resided in Site 2. Otherwise, the cation was considered unbound.
3) For a four-quartet system, occupancies of Sites 0 and 4 (above or below the GQ) were determined as for the two-quartet system. Else if the cation distance to the 5′-quartet was shorter than to the 3′-adjacent quartet, and the cation was not farther than 4.5 Å from any of the O6 atoms in the 5′-adjacent quartet, the cation was considered to be in Site 1. Analogously, the occupation of Site 3 was determined. If none of the conditions was true and cation distance to the 5′-adjacent quartet was shorter than to the 3′-quartet and at the same time the distance to the 3′-adjacent quartet was shorter than to the 5′-quartet, while the cation was not farther than 4.5 Å from any of the O6 atoms in the 5′-adjacent quartet, the cation resided in Site 2. Otherwise, the cation was considered unbound.

Note that the used visualization algorithm does not specifically differentiate in-quartet-plane binding positions frequently sampled by $Na^+$ (but much less frequently by $K^+$) ions as it primarily aims to visualize binding in the cavities between the G-quartets. Such an in-plane binding site within a particular G-quartet can be manifested in the cation-binding graphs as very frequent exchanges of a single cation between two adjacent sites or even as a simultaneous occurrence of a single cation in two binding sites due to the limited visual resolution of the individual points (we use typically 1000 points per microsecond) in the graphs. However, we emphasize that we have monitored the ion positions also by other means, such as the distance of the ions from the GQ center of the mass and by visual inspection of the trajectories. Therefore, ion dynamics in the trajectories was unambiguously resolved. We have selected the "cavity-centered" ptraj-based analysis for presentation of the results, as it provides the most compact information about the ion dynamics. Nevertheless, the text description is always based on the overall analysis using all the tools, provides the most accurate qualitative information about the ion behavior and should be taken as the most relevant information. Note also that while the internal ions (bound in the cavities between two G-quartets) are always restricted in their movements to the central channel, the ions detected at the outer sites can be loosely bound. Complete quantitative description of the ion positions would be a multidimensional description that would complicate the reading. Note also that due to the density of points the Figures cannot be used to estimate populations of the ion binding sites. Therefore, we report occupancy of each site (as defined by the used criteria) in all the Figures by numbers that are placed right to the graphs.

#### *Supporting Results*

##### Description of the simulations of individual GQ units of 2N3M (Simulations 4a - 4h)

In the simulation of the first GQ unit in the SPC/E with $Na^+$ (Simulation 4a), the second strand rotated and aligned perpendicular to the other three strands at 400 ns. The third strand also rotated subsequently, followed by the misalignment of the first and fourth strands (Main text, Figure 3b). The GQ completely unfolded within 20 ns after the first perturbation event. Although six ion-exchange events took place before the unfolding, the strand rotation specifically was not correlated with any ion movement.

In the simulation of the first GQ unit in SPC/E with $K^+$ (Simulation 4b), G8 moved into the solvent at 1.83 μs. Following this, the second strand rotated forming a cross-like structure leading to the unfolding of GQ (Main text, Figure 3c). No ion movement was observed in the stable part of this trajectory.

In the simulation in OPC with $Na^+$ (Simulation 4c), an additional $Na^+$ ion aligned below the GQ. The GQ was maintained till 1.08 μs, although the quartets were not entirely planar. Excessive buckling of quartets then led to the disintegration of the bottom quartet and rotation of the first strand (Main text, Figure 3d). G15 stacked with the T16 and moved towards the solvent. Then the first quartet also disintegrated within 2-3 ns leading to the complete unfolding of GQ.

In the simulation in OPC with $K^+$ (Simulation 4d) the GQ was stable till 6.58 μs. T1 and T14 stacked on G2 and G12, respectively and sampled T1(CH3)-T14(O2) hydrogen bond interaction. T14 moved slightly towards the groove at 6.45 μs and T11 aligned over the quartets at 6.52 μs. The alignment of T1, T14 and T11 caused a strain on the G-stem. Strand 2 slipped and stacked with T7 and then rotated to form a cross-like structure at 6.58 μs (Main text, Figure 3e). Subsequently, the GQ completely unfolded within 40 ns of the first perturbation event. No exchange of channel ions with the solvent was observed in the stable part of the simulation.

In the simulation of the second GQ unit in SPC/E and $Na^+$ (Simulation 4e), unfolding was observed at ~880 ns as the third strand of GQ rotated followed by the rotation of the second strand (Figure S5b). These two strands arranged in perpendicular to the other two strands to form a cross-like structure (Figure S5b).[1] Frequent (roughly every 50 ns) ion exchanges were observed in the initial 850 ns of the simulation (Figure S6a).

In the Simulation 4f in SPC/E and $K^+$, no ion movement was observed until 4.65 μs (Figure S6b). T1 stacked over G2 of strand 1 and T13 stacked below G12 of strand 4, respectively. Unfolding initiated at 4.65 μs as T1 moved to stack over G11 of strand 4 and T10 of the loop aligned at an angle with the bases. Ion exchange was observed and strand 1 slipped and rotated followed by the rotation of strand 2 (Figure S5c). Strand 3 also aligned perpendicularly to strands 1 and 2 leading eventually to complete unfolding of GQ.

In the Simulation 4g in OPC and $Na^+$, the first strand showed strand slippage followed by rotation at 25 ns. Strand-slippage is a movement of a strand by one step reducing the number of full quartets.[1, 2] The third strand also rotated at ~85 ns leading to the unfolding within next 40 ns (Figure S5d).

In the Simulation 4h in OPC and $K^+$, strand 2 rotated followed by the rotation of strand 3 and formed a cross-like structure at 1.3 µs leading to the unfolding of the GQ (Figure S5e).

**Description of Simulations 8a - 8f, 9a - 9b, 10**

In Simulation 8a, $K^+$ ions were added to all the three ion sites. Linear ion movement of $K^+$ between the second and third site was observed after 10 ns of the simulation. $K^+$ from the Site 3 left the GQ and within 2 ns $K^+$ ion from the Site 2 moved to occupy the Site 3. No further ion movements were observed and Site 2 remained vacant for all the remaining simulation time (Figure S13a). The hexad alignment as in the starting structure was not sampled. Instead, the loop nucleotides of the two GQs (A3|A15 and A9|A21) stacked together and aligned slightly away from the GQ groove towards the solvent. There were two new intermittently sampled interactions of loops with the guanines as A6 and A18 aligned across the grooves in both the GQs. A6 was tilted with reference to the plane of G-stem but A18(H62)-G16(N3) and A18(N7)-G16(H22) interactions were sampled for 60% of the simulation time.

All the ion sites were vacant in the starting structure of Simulation 8b. Site 3 was occupied during the equilibration stage by the entry of $K^+$ ion through the bottom quartet. Site 1 was occupied within 5 ns of the start of the simulation by the entry of $K^+$ ion through the top of the GQ. As in the other simulations with $K^+$ ions, Site 2 remained vacant and there was no exchange of ions throughout the 6.8 µs long simulation (Figure S13b). A6 and A18 aligned in the groove along the G-stem throughout the simulation and did not sample any planar alignment. The loop nucleotides of the two GQs (A3|A15 and A9|A21) stacked together and aligned slightly away from the GQ groove towards the solvent.

In the 6.8 µs long Simulation 8c, Site 1 was occupied in the starting structure. Site 3 was occupied through the entry of $K^+$ within the first ns of the start of the simulation. There was no ion exchange event in this simulation as well (Figure S13c). As in Simulations 8a and 8b, A3|A15 and A9|A21 stacked together but could not sample any alignment with the G-stem.

In the 10 μs long Simulation 8d, only Site 2 was occupied in the starting structure. $K^+$ from the bulk occupied Site 3 within 100 ns of the simulation, pushing the ion from Site 2 to Site 1. Thus both Site 1 and Site 3 were occupied while Site 2 remained vacant for the remaining part of the simulation (Figure S13d). The loop nucleotides of the two GQs (A3|A15 and A9|A21) stacked together and aligned slightly away from the GQ groove towards the solvent. The (GGGG)A pentad was sampled intermittently during the simulation as A15 formed hydrogen bonds with G13 while A3 formed hydrogen bonds with G1 (Figure S14).

In the 10 μs long Simulation 8e, $K^+$ ions were placed at Site 1 and 2 in the GQ. The $K^+$ ion from Site 2 shifted to Site 3 at 190 ns of the start of the simulation. In the Simulation 8f, two $K^+$ ions at Site 1 and 3 were placed in the GQ. No cation exchange was observed in the simulations and Site 2 was vacant throughout the simulations (Figures S13e and S13f).

In summary, the RNA GQ dimer was stable in all the simulations. In addition, the channel eventually bound two $K^+$ ions at Sites 1 and 3, regardless of the initial ion placement.

RNA GQ dimer was also stable in the 5 μs long simulations carried out with $Na^+$ as the stabilizing ions in SPC/E (Simulations 9a - 9b) and OPC water model (Simulation 10). In all the simulations, $Na^+$ ion from Site 2 moved to Site 1 or 3 and the original ion from the site escaped into the solvent. $Na^+$ ions were present only at Sites 1 and 3 and no exchange of ion with the solvent was observed (Figures S15 and S16). Additional cation binding sites were developed just above and below the G-stem in all the simulations (Figure S15 and S16).

**Description of Simulations 16a - 16d**

The two-quartet RNA GQ represented by a monomer of 2RQJ was also a good test system for the TIP4P-D water model. It was stable in three out of four TIP4P-D Simulations 16a-16d (Figure S22). Multiple cation exchanges were observed in the Simulation 16a carried out in JC $Na^+$ ions but GQ was maintained throughout the 5 μs simulation (Figures S22a and S23a). The GQ completely unfolded in the Simulation 16b carried out with CHARMM22 $Na^+$ ions (Figure S22b). Excessive buckling of quartets was observed in this simulation from the start. One cation escaped through the first quartet at 1.8 μs and the GQ then completely unfolded within few ns (Figure S23b).

In the Simulations 16c and 16d carried out in JC and CHARMM22 $K^+$ ions, respectively the GQ was stable and no cation exchange with the solvent was observed (Figures S22c-S22d and S23c-S23d).

**Description of Simulation 17**

In the two-quartet DNA GQ with adenine flanking base and single nucleotide loops, A1 stacked over G2 while the loop bases aligned in the grooves of the GQ. As A1 moved to stack over G11 of strand 4 at 450 ns, strand 1 slipped and rotated leading to the unfolding of the GQ (Figure S24). No exchange of channel cation was observed for the stable part of the trajectory. The simulation showed that the substitution of thymine by adenine as flanking base and loops could not increase the GQ stability significantly. In other words, two-quartet DNA GQs with single nucleotide loops are unstable regardless of bases constituting the loops.

**Description of tetramolecular antiparallel RNA G-stem model (Simulations 18a - 18d)**

The model of hypothetical tetramolecular two-quartet antiparallel RNA G-stem unfolded in all the simulations in $Na^+$ and $K^+$ ions in the SPC/E water model. The quartets were not planar and the guanines were buckled from the start of the simulations. The GQ unfolded in the two $Na^+$ simulations at 10 and 50 ns by strand slippage followed by strand rotation (Figure S25a and S25b). The GQ unfolded at ~180 ns and ~510 ns in the two $K^+$ simulations (Figures S25c and S25d).

**Description of Simulations 27a-27d**

Further parallel-stranded d[GGGG]$_4$ simulations were carried out with TIP4P-D water with JC and CHARMM22 $Na^+$ and $K^+$ ions (simulation series 27). The d[GGGG]$_4$ G-stem was fully stable in only two out of four simulations in TIP4P-D water model. Strand slippage was observed in the Simulation 27a in JC $Na^+$ ions at ~3 μs (Figure S39). However, the G-stem was stable in the Simulations 27b and 27c in CHARMM22 $Na^+$ ions and JC $K^+$ ions, respectively. In the simulation with CHARMM22 $Na^+$ ions, cations were present in all the three sites and ion exchanges were observed while in the simulation with JC $K^+$ ions, cations were retained at Sites 1 and 3 while Site 2 was vacant for the majority of simulation time (Figures S40b and S40c). Disruption of 5′-quartet was observed in the Simulation 27d with CHARMM22 $K^+$ ions at ~1.3 μs as cation escaped from the top quartet (Figure S40d). The quartet reassembled at 3.6 μs and full four-quartet G-stem was formed again (Figure S39d). $Na^+$ ions were more dynamic than $K^+$ ions (Figure S40). The perturbation of quartets and excessive ion dynamics suggest that the combination of cations and TIP4P-D water parameter tested here is probably less suitable for GQ simulations than SPC/E with JC ions.

**Simulations of two-quartet parallel and antiparallel DNA G-stems using Langevin thermostat (Simulations 28a - 28f)**

Strand slippage and subsequent structure loss was observed in two independent simulations, 28a and 28b of parallel-stranded all-*anti* d$[GG]_4$ G-stem in $K^+$ ions and SPC/E water model similar to Simulations 5c and 5d (Figure S41a). The antiparallel d$[GG]_4$ G-stem was stable in all four independent simulations (Simulations 28c-28f) similarly to simulation series 7 (Figure S41b). In all four simulations of antiparallel d$[GG]_{4,}$ multiple (>10 per microsecond) cation exchange events were observed suggesting that the choice of thermostat had no systematic effect on GQ simulations and any outcome of the study (Figure S42).

**Description of simulations of 2RQJ dimer and monomer using Langevin thermostat (Simulations 29a - 29f)**

All the simulations in the series 29 were carried out with $K^+$ ions and SPC/E water model and the temperature was controlled using Langevin thermostat. Simulations 29a - 29c were carried using two-quartet RNA dimer 2RQJ with cations at different sites in the GQ in the starting structure. Simulations 29d - 29f were carried out using monomer of two-quartet RNA GQ.

The 2RQJ Simulation 29a was initiated with cations in all the three sites within the GQ equivalent to Simulation 8a with Berendsen thermostat. Cation from Site 3 moved in-plane with the fourth quartet at 440 ns followed by the movement of Site 2 cation into Site 3 at 500 ns and leading to the loss of original Site 3 cation into the solvent. No further movement of cations within the GQ was observed and Site 2 remained vacant until the end of the Simulation 29a (Figure S43b). In the Simulation 29b, cation was incorporated only at Site 1 and Sites 2 and 3 were vacant in the starting structure of 2RQJ equivalent to Simulation 8c. A cation entered into Site 3 through the fourth quartet at 50 ns of the simulation. The two cations were then retained within the GQ and no exchange of cations with the solvent was observed (Figure S43c). Simulation 29c was initiated with cations in Sites 1 and 2 within the GQ equivalent to Simulation 8e.

Cation from Site 2 moved to Site 3 at 100 ns and was retained until the end of the simulation (Figure S43d). Thus, the behaviour of cations in Simulations 29a, 29b and 29c were similar to equivalent simulations (Simulations 8a, 8c and 8e) carried out with Berendsen thermostat.

In the simulations of 2RQJ monomer, 29d - 29f, A12 at the 3′-end stacked below G11 of the second quartet during the entire simulation time. The loop bases either oriented in the solvent or aligned in the groove of the GQ and showed some dynamics. The G-stem was stable and no exchange of channel cation was observed in any of the simulations similar to the equivalent simulation series 11 with Berendsen thermostat (Figure S44).

#### *Supporting Tables*

**Table S1**. List of RNA GQ simulations carried out in the present study

| Number<br><br>Simulation | subcategory | Structure | Length of simulation (μs) | Conditions<br><br>(force field /ion/ water model, initial ion positions)[a] | Summary |
|---|---|---|---|---|---|
| **8** | a | 2RQJ | 10 | OL3/$K^+$/SPC/E | GQ with two ions, Site 2 vacant. |
| | b | | 6.8 | OL3/$K^+$/SPC/E<br>$K^+$ in none of the three sites | GQ with two ions, Site 2 vacant. |
| | c | | 6.8 | OL3/$K^+$/SPC/E<br>$K^+$ in Site 1 only | GQ with two ions, Site 2 vacant. |
| | d | | 10 | OL3/$K^+$/SPC/E<br>$K^+$ in Site 2 only | GQ with two ions, Site 2 vacant. |
| | e | | 10 | OL3/$K^+$/SPC/E<br>$K^+$ in Site 1 and 2 only | GQ with two ions, $K^+$ ion moved from Site 2 to Site 3. No ion exchange event. |
| | f | | 10 | OL3/$K^+$/SPC/E<br>$K^+$ in Site 1 and 3 only | GQ with two $K^+$ ions, Site 2 vacant. No ion exchange event. |
| **9** | a-b | 2RQJ | 5 (2 simulations) | OL3/$Na^+$/SPC/E | $Na^+$ ions only at Site 1 and 3, additional cation binding sites above and below the G-stem. |
| **10** | a | 2RQJ | 5 | OL3/$Na^+$/OPC | $Na^+$ ions only at Site 1 and 3, additional cation binding sites above and below the G-stem. |
| **11** | a | two-quartet GQ unit of 2RQJ | 10 | OL3/$K^+$/SPC/E | GQ stable; no ion exchange. |
| | b | | 10 | OL3/$K^+$/SPC/E | GQ stable; no ion exchange. |
| | c | | 10 | OL3/$K^+$/SPC/E | GQ stable; no ion exchange. |
| | d | | 10 | OL3/$K^+$/SPC/E | GQ stable; no ion exchange. |
| **12** | a-d | two-quartet GQ unit of 2RQJ without 3′-flanking base | 5 (4 simulations) | OL3/$Na^+$/SPC/E | multiple events of ion exchange but GQ is stable except for one simulation where the GQ unfolded at 4.27 μs. |
| **13** | | two-quartet GQ unit of 2RQJ without 3′-flanking base | 1 | OL3/$K^+$/SPC/E | GQ stable; no ion exchange. |

| | | | | | |
|---|---|---|---|---|---|
| **14** | a-c | two-quartet GQ unit of 2RQJ without 3′-flanking base | 1 (3 simulations) | OL3/$Na^+$/OPC | GQ was stable in one simulation and unfolded in two simulations at 400 ns and 130 ns. |
| **15** | a-c | two-quartet GQ unit of 2RQJ without 3′-flanking base | 1 (3 simulations) | OL3/$K^+$/OPC | GQ stable; no ion exchange. |
| **16** | a | two-quartet GQ unit of 2RQJ | 5 | OL3/$Na^+$/ TIP4P-D | GQ stable, exchange of channel cations. |
| | b | | 5 | OL3/CHARMM22 $Na^+$/ TIP4P-D | excessive buckling of G-quartets, cation lost through the first quartet at 1.8 µs followed by complete unfolding of GQ. |
| | c | | 5 | OL3/$K^+$/ TIP4P-D | GQ stable, no cation exchange. |
| | d | | 5 | OL3/CHARMM22 $K^+$/ TIP4P-D | GQ stable, no cation exchange. |

[a] fully occupied channel at the beginning if not mentioned otherwise. JC ions if not mentioned otherwise.

**Table S2.** List of three and four-quartet G-stem simulations carried out in the present work. Some additional simulations are also included in this Table.

| Number<br><br>Simulation | subcategory | Structure | Length of simulation (µs) | Conditions<br><br>(force field /ion/ water model)[a] | Summary |
|---|---|---|---|---|---|
| **17** | | modified DNA two-quartet GQ with adenine loops | 1 | OL15/$K^+$/SPC/E | Unstable, lost at ~ 450 ns. |
| **18** | a-d | antiparallel r[GG]$_4$ G-stem | 2 (2 simulations)<br><br>2 (2 simulations) | OL3/$Na^+$/SPC/E<br><br>OL3/$K^+$/SPC/E | Unfolded at ~10 ns and 50 ns.<br><br>Unfolded at ~180 ns and 510 ns. |
| **19** | a-b | parallel-stranded all *anti* d[GGG]$_4$ G-stem | 5 (2 simulations) | OL15/$Na^+$/SPC/E | a) Strand slippage in one strand at 1.6 µs.<br><br>b) Strand slippage at 530 ns. |
| **20** | a-b | parallel-stranded all *anti* d[GGG]$_4$ G-stem | 6<br><br>5 | OL15/$K^+$/SPC/E | a) GQ stable, one ion exchange at 4.8 µs.<br><br>b) G-stem stable, one cation exchange at 273 ns. |
| **21** | a | antiparallel d[GGG]$_4$ G-stem | 2 | OL15/$Na^+$/SPC/E | G-stem stable with few ion exchange events. |
| | b | antiparallel d[GGG]$_4$ G-stem | 2 | OL15/$K^+$/SPC/E | G-stem stable with no ion exchange event. |
| | c | antiparallel d[GGG]$_4$ G-stem | 2 | OL15/$Na^+$/OPC | Strand slippage observed. |
| | d | antiparallel d[GGG]$_4$ G-stem | 2 | OL15/$K^+$/OPC | G-stem stable with no ion exchange event. |
| **22** | a-e | parallel-stranded all *anti* d[GGGG]$_4$ G-stem | 1 (5 simulations) | OL15/$Na^+$/SPC/E | Multiple events of ion exchange in four simulations and G-stem was stable in all the simulations. |
| **23** | a-e | parallel-stranded all *anti* d[GGGG]$_4$ G-stem | 1 (5 simulations) | OL15/$K^+$/SPC/E | Ion from Site 1 was pushed over the first quartet, then frequent ion exchanges at this site with the solvent $K^+$ ions. G-stem was stable in all the simulations. |
| **24** | a-e | parallel-stranded all *anti* d[GGGG]$_4$ G- | 1 (5 simulations) | OL15/$Na^+$/OPC | G-stems were perturbed in three out of five simulations as strand slippage event was |

| | | | | | |
|---|---|---|---|---|---|
| | | stem | | | observed in these simulations. |
| **25** | a | parallel-stranded all *anti* d[GGGG]$_4$ G-stem | 1 (5 simulations) | OL15/$K^+$/OPC | Disruption of 5′-quartet at 180 ns, the other three quartets were maintained. |
| | b | | | | Disruption of 5′-quartet at 280 ns, the other three quartets were maintained. |
| | c | | | | Disruption of 5′-quartet at 350 ns, the other three quartets were maintained. |
| | d | | | | G-stem stable, Site 2 was vacant for the majority of the simulation time. |
| | e | | | | G-stem stable, all three ion sites were occupied, ion exchange at Site 3 at ~740 ns. |
| **26** | a | parallel-stranded all *anti* d[GGGG]$_4$ | 2 | OL15/$Na^+$ (LM ions) /SPC/E | Three ions stabilized the G-stem with diverse ion exchange events. |
| | b | | 2 | OL15/$K^+$ (LM ions)/SPC/E | Ion from Site 3 was lost to the solvent at 90 ns and guanine from the last quartet moved towards the solvent, corruption of the G-stem. |
| | c | | 2 | OL15/$K^+$ (LM ions)/SPC/E | G-stem stable, ion from Site 2 moved to Site 1 and Site 2 was vacant for the majority of the simulation time. |
| | d | | 2 | OL15/ $Na^+$ (HFE ions)/OPC | GQ stable but with diverse ion-exchange events. The last quartet was a bit buckled up. |
| | e | | 2 | OL15/ $K^+$ (HFE ions)/OPC | Disruption of 5′-quartet at 1.7 µs, the other three quartets were maintained. |
| | f | | 2 | OL15/ $Na^+$ (IOD ions)/OPC | Disruption of 5′-quartet at 1.0 µs, the other three quartets were maintained. |
| | g | | 2 | OL15/ $K^+$ (IOD ions)/OPC | Disruption of 5′-quartet at 400 ns, the other three quartets were maintained. |
| **27** | a | parallel-stranded all *anti* d[GGGG]$_4$ | 5 | OL15/$Na^+$/TIP4P-D | Strand slippage at ~3 µs, cation exchange observed. |
| | b | | 5 | OL15/CHARMM22 | G-stem stable, cation exchange |

| | | | | | |
|---|---|---|---|---|---|
| | | | | $Na^+$/ TIP4P-D | observed. |
| | c | | 5 | OL15/$K^+$/ TIP4P-D | G-stem stable, $K^+$ at Site 2 was vacant for the majority of the time. |
| | d | | 5 | OL15/CHARMM22 $K^+$/ TIP4P-D | Disruption of 5′-quartet at ~1.3 µs but reformed at 3.6 µs. |
| **28** | a-b | parallel-stranded all *anti* d[GG]$_4$ | 2 (2 simulations) | Langevin thermostat<br>OL15/$K^+$/ SPC/E | Strand slippage observed at ~100 ns in one and at ~200 ns in the second simulation, then the loss of the structure. |
| | c-f | antiparallel d[GG]$_4$ | 5 (4 simulations) | Langevin thermostat<br>OL15/$Na^+$/ SPC/E | Stable but buckling, ion exchange observed. |
| **29** | a | 2RQJ | 5 | Langevin thermostat OL15/$K^+$/ SPC/E | $K^+$ ion moved from Site 2 to Site 3 and original Site 3 cation was lost. GQ with two ions at Site 1 and 3. No ion exchange event; similar to Simulation 8a. |
| | b | | 5 | Langevin thermostat OL15/$K^+$/ SPC/E $K^+$ ions at Site 1 only | $K^+$ ion was taken from solvent to Site 3. GQ with two ions at Site 1 and 3. No ion exchange event; similar to Simulation 8c |
| | c | | 5 | Langevin thermostat OL15/$K^+$/ SPC/E $K^+$ ions at Site 1 and 2 only | $K^+$ ion moved from Site 2 to Site 3. GQ with two ions at Site 1 and 3. No ion exchange event; similar to Simulation 8e |
| | d - f | 2RQJ monomer | 5 | Langevin thermostat OL15/$K^+$/ SPC/E | No ion exchange in any of the simulations |

[a] fully occupied channel at the beginning if not mentioned otherwise. JC ions if not mentioned otherwise.

#### *Supporting Figures*

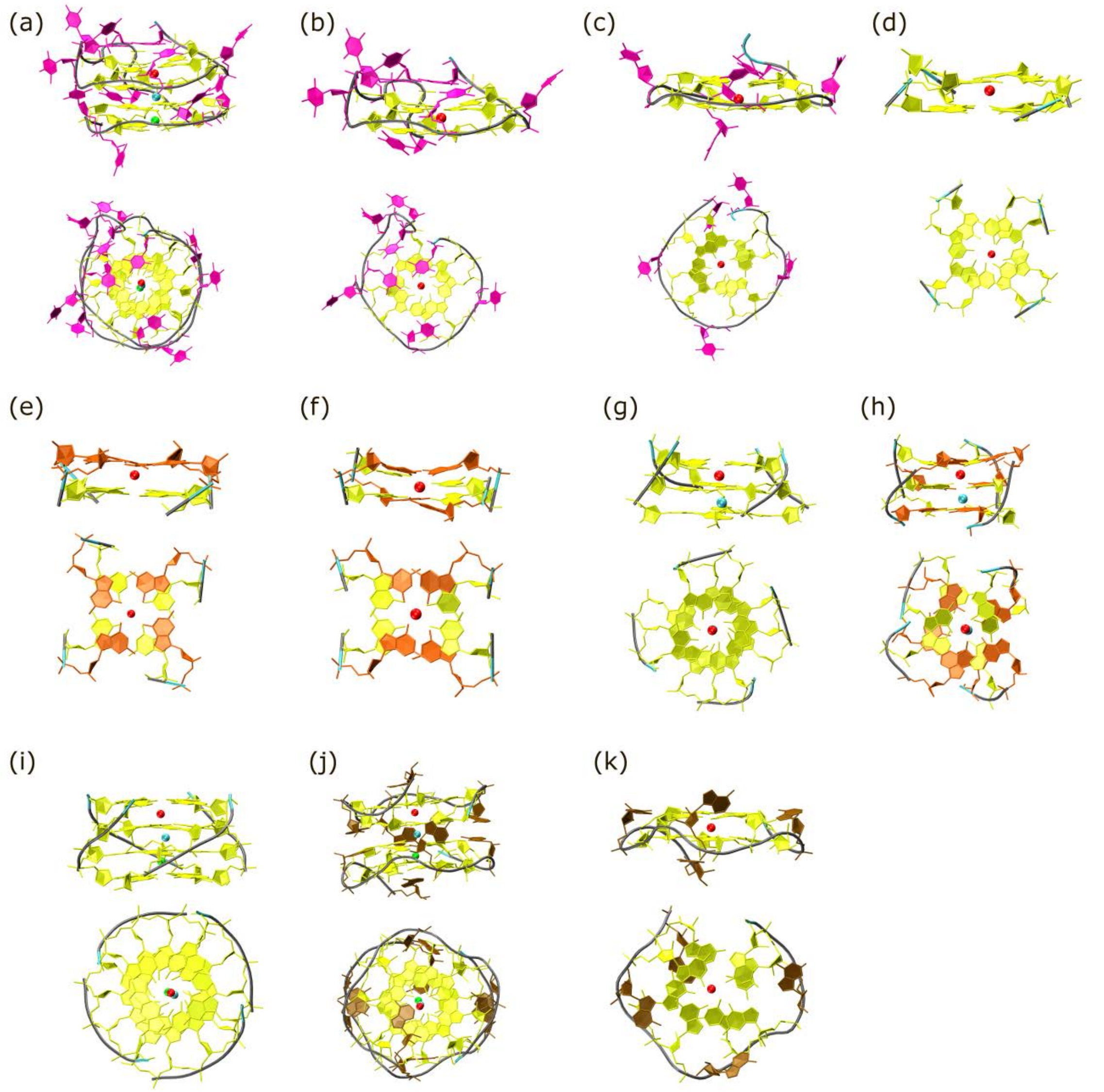


**Figure S1**. Side and top view of atomistic structures used in the simulations. (a) 2N3M GQ, (b) two-quartet GQ extracted from upper half of 2N3M, (c) two-quartet GQ extracted from lower half of 2N3M, (d) all *anti* parallel-stranded d[GG]$_4$ G-stem, (e) parallel-stranded d[GG]$_4$ with 5′-guanines in *syn* orientation, (f) antiparallel d[GG]$_4$ extracted from 148D, (g) parallel-stranded d[GGG]$_4$ from 1KF1, (h) antiparallel d[GGG]$_4$ from 143D and (i) parallel-stranded d[GGGG]$_4$ from 4R44 are shown. The RNA simulations were carried out using the dimer of (j) 2RQJ and (k) monomer of 2RQJ GQ. The *anti* and *syn* guanine nucleosides are shown in yellow and orange, respectively. The thymine and adenine nucleosides are shown in magenta and brown. The backbone is shown in grey while the 5′-end is shown in cyan. Cations within the G-stem are shown as red, cyan and green spheres.

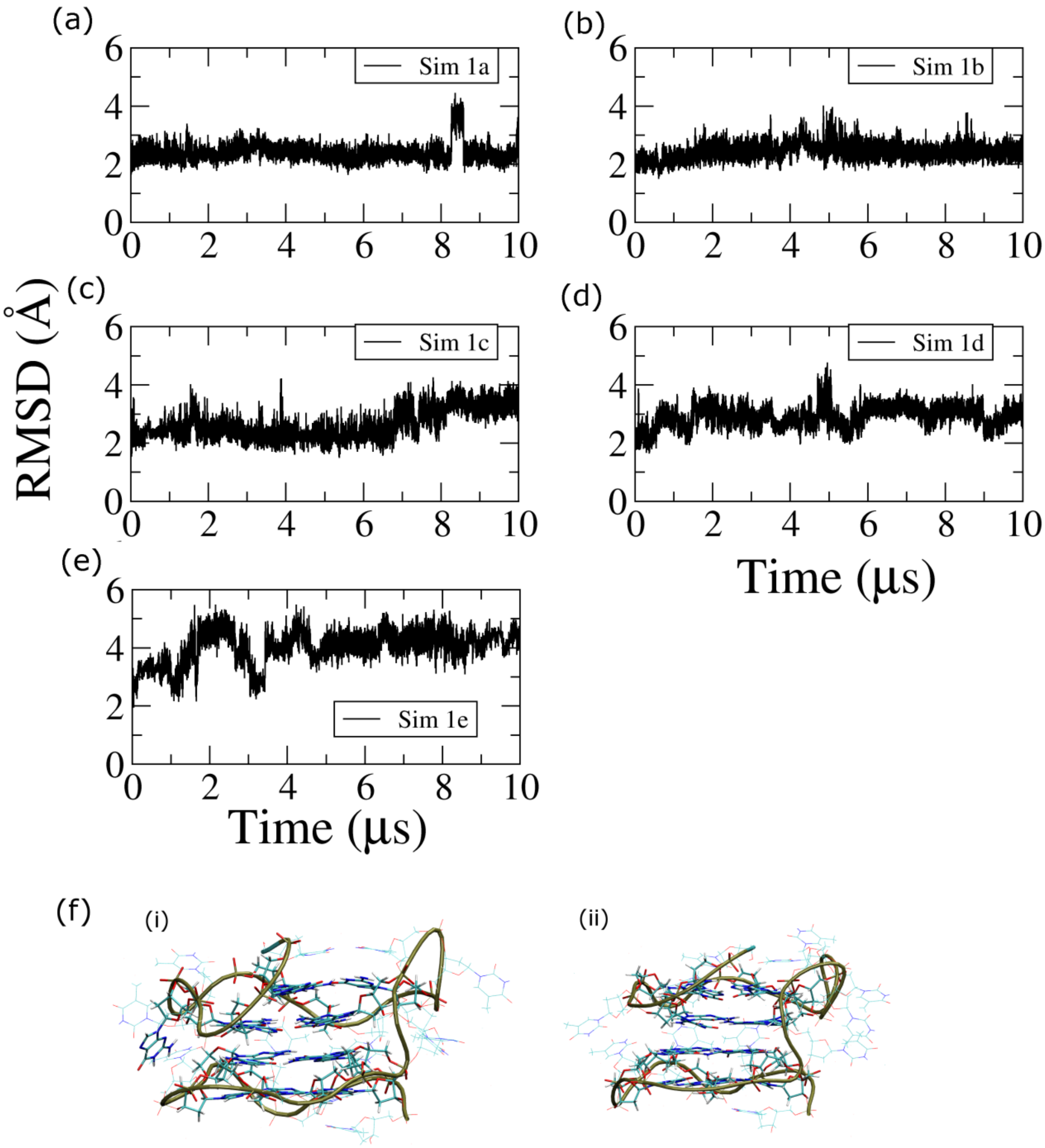


**Figure S2**. Backbone RMSD of 2N3M in the Simulations (a) 1a, (b) 1b, (c) 1c, (d) 1d and (e) 1e. The simulations in SPC/E (1a - 1b) show generally lower RMSD than those in the OPC water model. Two simulations were carried out in $K^+$ and OPC water model as the quartet at 5′-end was disrupted in the first $K^+$ and OPC simulation (Simulation 1d). (f) 2N3M at the end of Simulations (i) 1d and (ii) 1e in $K^+$ and OPC water model. Guanine and adenine nucleotides are shown in licorice and lines, respectively while the backbone of GQ is shown in tan tube. The 5′ end of the GQ backbone is shown in cyan. Panel (i) shows disruption of the first quartet in Simulation 1d as G5 moved towards the solvent. GQ was stable in Simulation 1e and is shown in the panel (ii).

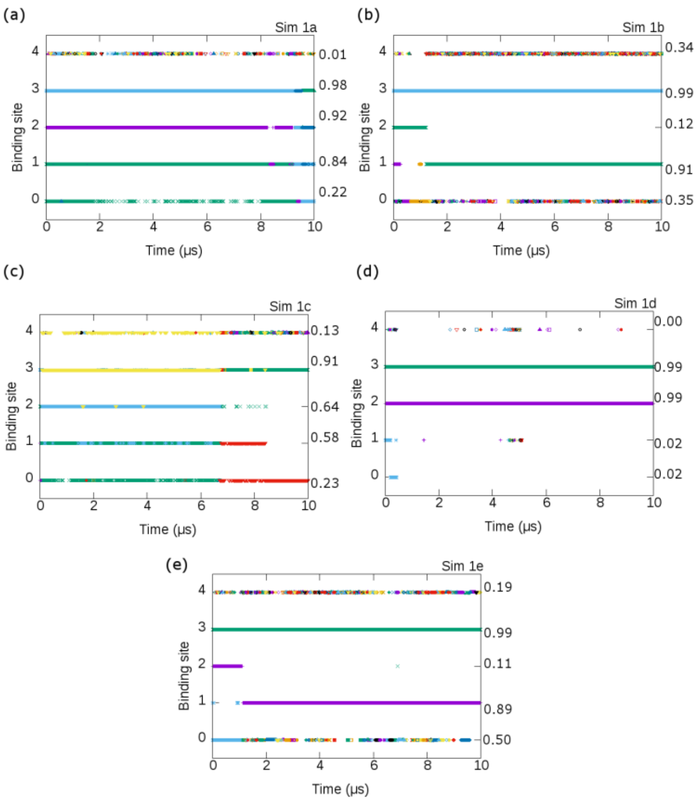


**Figure S3**. Cation binding to the GQ in the Simulations 1a - 1e. The cation binding Sites 1, 2 and 3 correspond to the typical position of cations in the channel of GQ between the two quartets starting from the 5′-end. The Sites 0 and 4 stand for the positions over the 5′-quartet and below the 3′-quartet, respectively. See panel (i) of Figure 1 for the representation of these sites. Distinct ions are represented by different colors and symbols, and any change in the color/symbol indicates an exchange of an ion in the site. The numbers on the right show occupancies of the respective sites. In the simulations carried out in SPC/E water model, (a) three $Na^+$ ions stabilized the GQ and two ion-exchange events were observed after 9 μs at Site 3 causing concerted linear movement of cations while (b) two $K^+$ ions were retained in the channel of 2N3M in the Simulation 1b. In the simulation in $Na^+$/OPC, (c) three $Na^+$ ions were present till 6.71 μs and Site 2 was empty in the remaining part of the Simulation 1c. In the simulations in $K^+$/OPC (d) two $K^+$ ions were present throughout the Simulation 1d as the first quartet was disrupted at 1.45 μs and (e) in the Simulation 1e, $K^+$ ion from Site 1 escaped into the solvent at 1.1 μs and subsequently ion from Site 2 moved to occupy Site 1. The GQ was stabilized by two $K^+$ ions for the remaining part of the simulation. Our algorithm analyzes geometrical parameters even if one quartet is lost or if the GQ is disrupted, but the results of such analyses are already less relevant as in Simulation 1d. Note that apparent simultaneous presence of cations at two binding sites (cavities) with the used visualization method indicates the presence of cation within the plane of a G-quartet; see the explanation in the "Visualization of cation binding sites" section at the beginning of the Supporting Information. The situation is relevant for some simulations in the presence of $Na^+$ ions as they tend to stay in G-quartet planes. The ion is then present in the plane of the G-quartet that is shared by the two ion-binding cavities. Presence of an ion in an outer G-quartet may be in the plots occasionally manifested as occupation of the outer sites above and below the G-stem. The text description is always accurate as it is based on the utilization of also other descriptors and on visual monitoring.

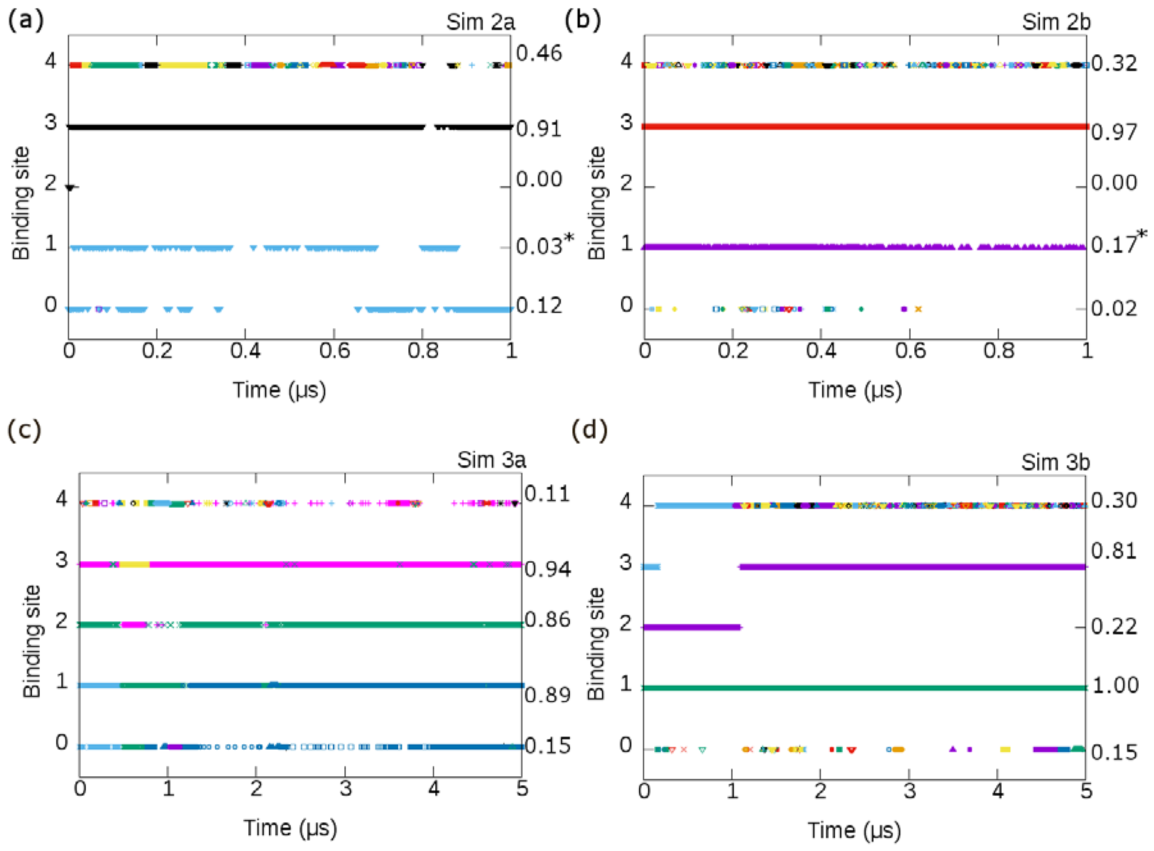


**Figure S4**. Cation movement in the SPC/E Simulations (a) 2a, (b) 2b, (c) 3a and (d) 3b. (a) and (b) Simulations 2a and 2b of 2N3M were carried out with $Na^+$ and $K^+$ ions initially present only in the solvent, i.e., absent in the channel. The $Na^+$ and $K^+$ entered the G-stem from top or bottom of the G-stem at Sites 1 and 3. Site 2 was vacant throughout the simulations. The numbers on the right show occupancies of the respective sites. In the sites marked by an asterisk (*), the occupancies were underestimated by the algorithm due to the presence of a water molecule which increases O6-ion distance. The presence of cation has been visually confirmed in these cases. Each color/symbol represents a different cation in the simulations. Note that the initial cation entry is so fast that it is not resolved on the figures in panels a and b. (c) and (d) The starting structure in the Simulations 3a and 3b was 2N3M dimer without any bond between the GQ units. In the Simulation 3a carried out with $Na^+$, cation exchange events were observed until 2.28 μs. Two $K^+$ ions stabilized the GQ in the Simulation 3b. Sites 1 and 2 were occupied while Site 3 was vacant till 1.08 μs after which $K^+$ ion from Site 2 moved to Site 3. For more details see the legends to Figure S3.

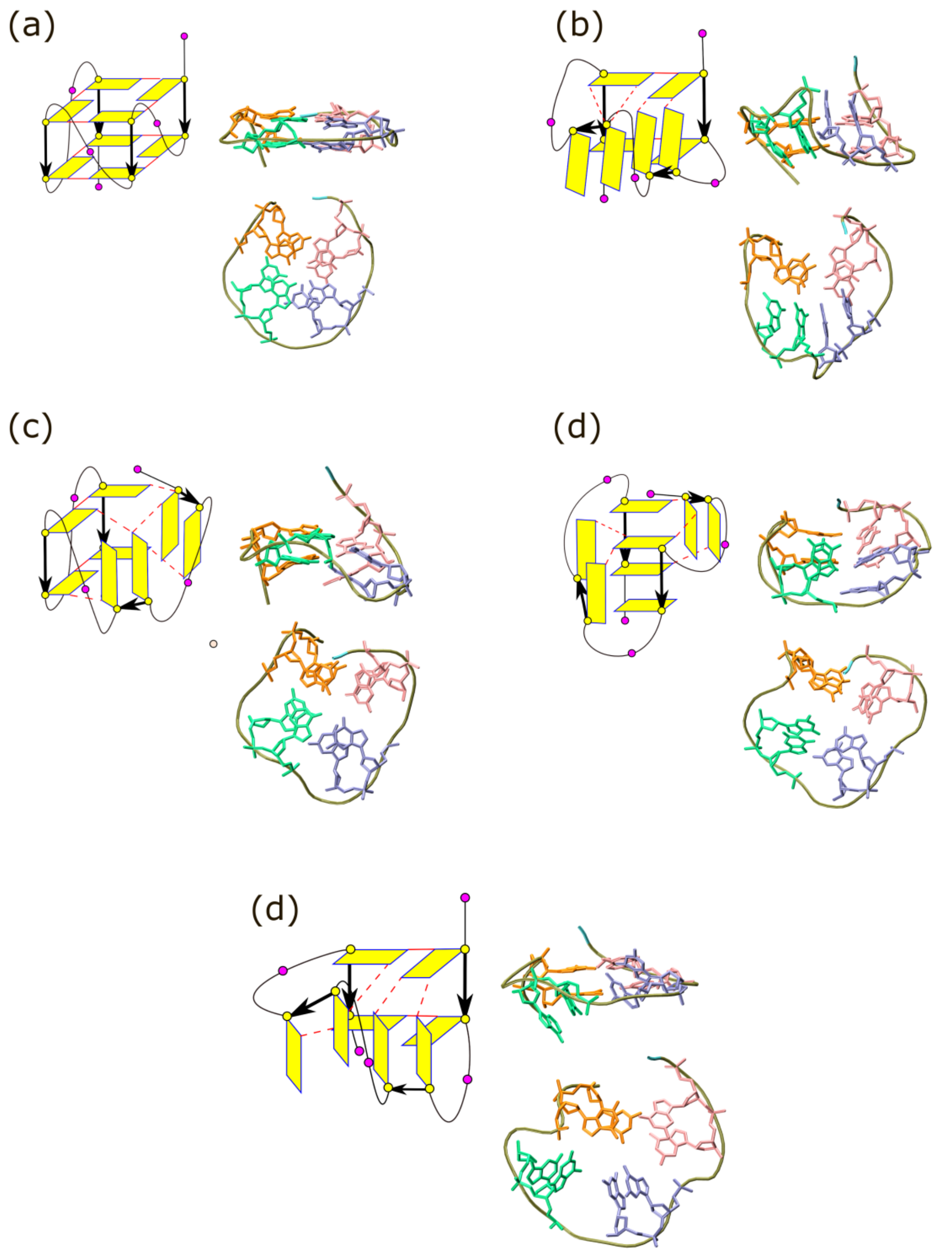


**Figure S5**. Simulations of the second GQ unit of 2N3M (Simulations 4e-4h). The two-quartet structure unfolded in all the simulations similar to the first GQ unit (cf. Figure 3 in the main text). Schematic and atomistic representations (both side and top view) of (a) starting structure and main transient structure occurring during the unfolding in (b) $Na^+$/SPC/E, (c) $K^+$/SPC/E, (d) $Na^+$/OPC and (e) $K^+$/OPC are shown. See legend to Figure 1 in the Main manuscript for further details of the schematic representations. The solid red lines represent characteristic Hoogsteen base pairing while dash red lines indicate any other hydrogen bonds. In the atomistic representations, the backbone of GQ is shown as tan tube with the 5′-end shown in cyan. The first, second, third and fourth strands of the GQ are shown in pink, blue, green and orange sticks, respectively.

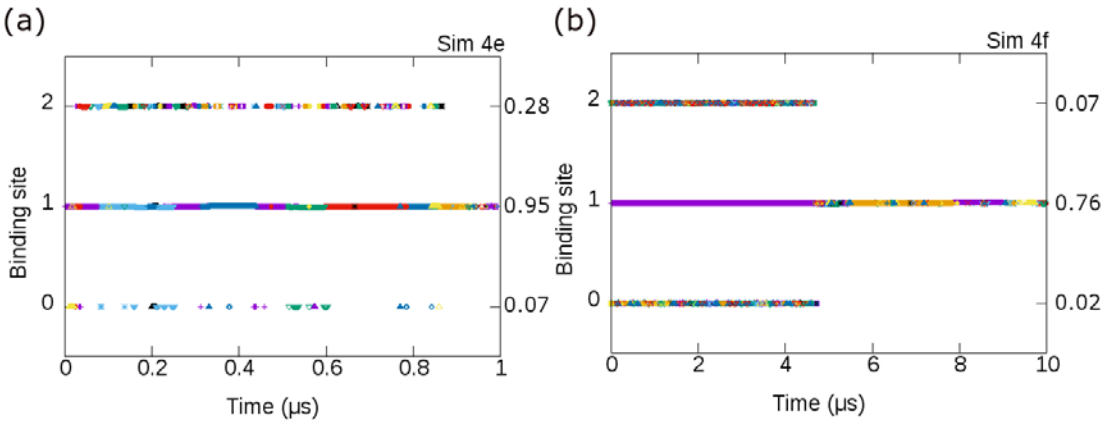


**Figure S6**. Cation movement in the Simulations 4e - 4f of second GQ unit of 2N3M in the SPC/E water model. (a) In the Simulation 4e carried out with $Na^+$, frequent ion-exchange events were observed until the GQ disrupted at 0.88 μs. (b) The $K^+$ ions were stable in between the two quartets in the Simulation 4f until the GQ disrupted at ~4.65 μs. The cation binding Site 1 corresponds to the typical position of cations in the channel of GQ between the two quartets. Sites 0 and 2 stand for the positions over the 5′-quartet and below the 3′-quartet, respectively. The numbers on the right show occupancies of the respective sites. See panel (d) of Figure 1 in the Main manuscript for the representation of these sites. Distinct ions are represented by different colors and symbols, and any change in the color/symbol indicates an exchange of an ion in the site. Note that our visualization algorithm, due to the used definition, detects ions even if one quartet is lost or if the GQ is disrupted, but the results of such analyses are already less relevant. Disruptions are always specified in the text.

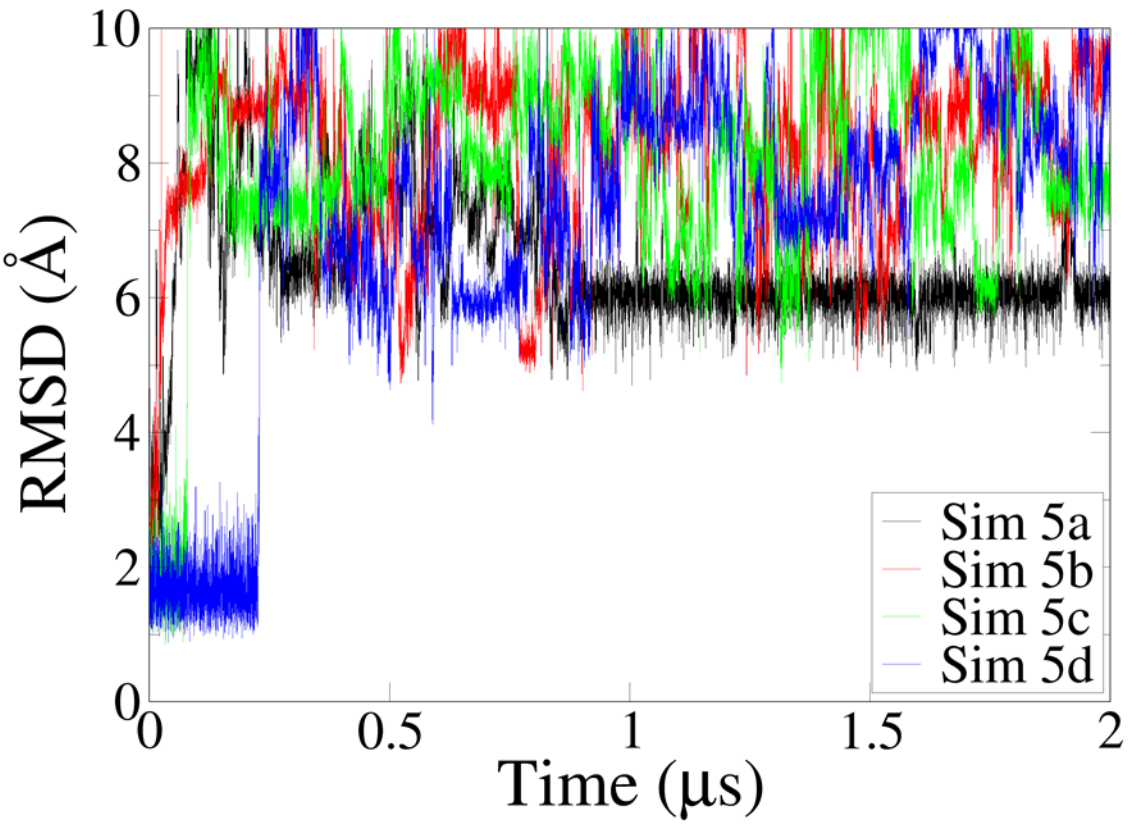


**Figure S7**. Backbone RMSD of all *anti* d[GG]$_4$ G-stem in the Simulations 5a - 5d. A significant change in backbone RMSD indicates unfolding of the G-stem. All the simulations were carried out in the SPC/E water model. Two simulations were carried out for both $Na^+$ and $K^+$ ions. The G-stem unfolded at 5 ns and 6 ns in $Na^+$ ions shown in black and red (Simulation 5a - 5b). The G-stem unfolded at 76 ns and 228 ns in the $K^+$ ion simulations and is shown in green and blue, respectively (Simulations 5c and 5d).

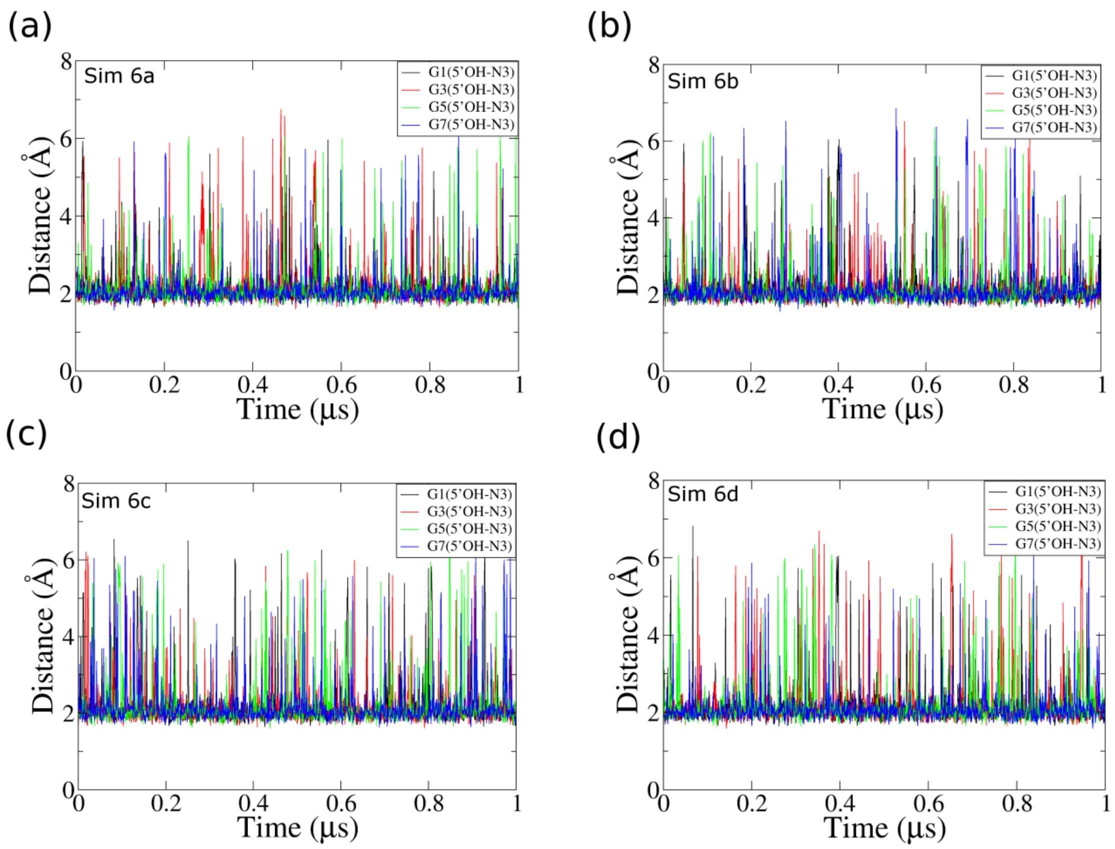


**Figure S8**. Distance plots monitoring the terminal *syn*-specific 5′-OH – G(N3) H-bonds in the Simulations 6a-6d of d[GG]$_4$ parallel-stranded G-stem. The 5′-bases of all the strands were in *syn* orientation. The hydrogen bonds between 5′-OH – G(N3) were sampled for the major time in Simulations (a) 6a, (b) 6b, (c) 6c and (d) 6d. The Simulations 6a and 6b were carried out in $Na^+$ and SPC/E water model while the Simulations 6c and 6d were carried out in $K^+$ and SPC/E water model.

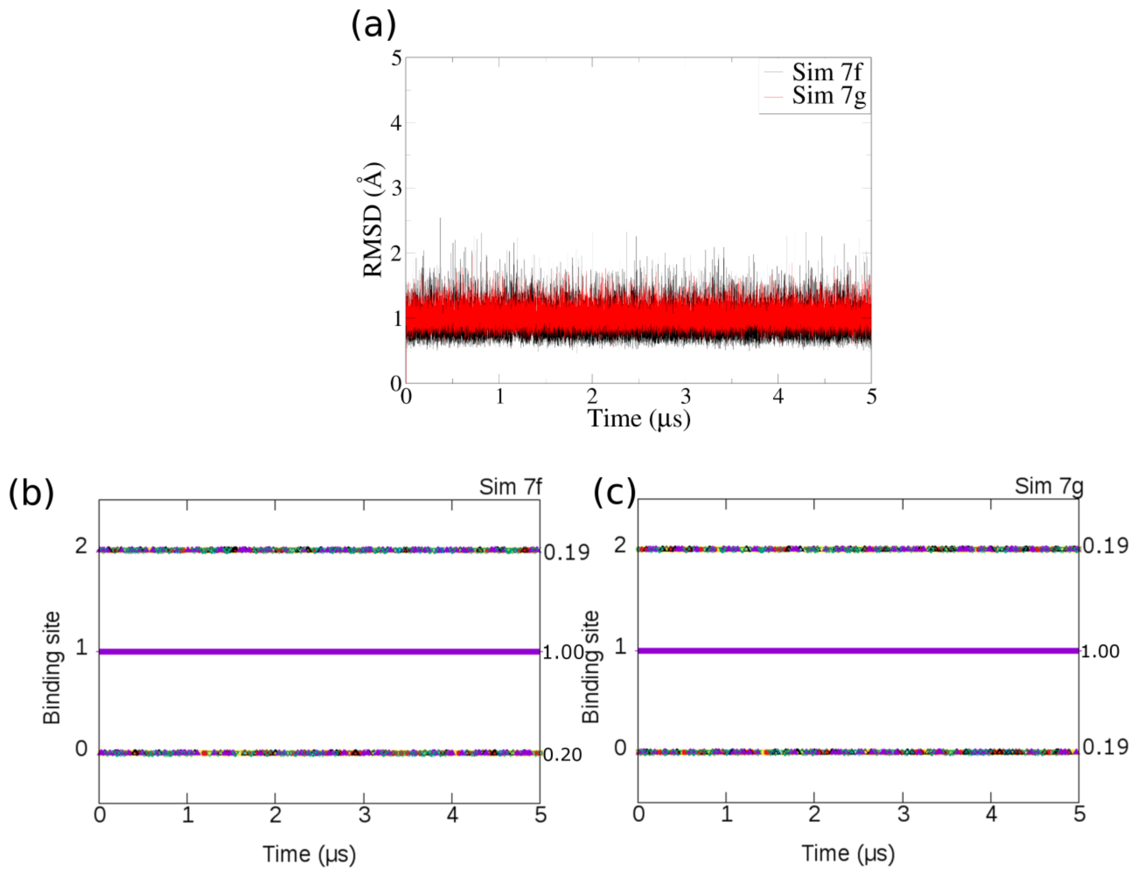


**Figure S9**. Backbone RMSD (a) and plots for monitoring cation movement (b-c) in the simulations of antiparallel d[GG]$_4$ G-stem from PDB 148D carried out in $K^+$ ions and SPC/E water model (Simulations 7f and 7g). (a) Backbone RMSD plots showed that G-stem was stable in both the Simulations 7f and 7g, shown in black and red, respectively. Cation binding plots of simulations (b) 7f and (c) 7g showed that there were no ion exchange movements and one cation stabilized the G-stem. The cation binding Site 1 corresponds to the typical position of cations in the channel of GQ between the two quartets. Sites 0 and 2 stand for the positions over the 5′-quartet and below the 3′-quartet, respectively. In the panels b and c, the numbers on the right show occupancies of the respective sites For full details, see the legend to Figure 4 of the Main manuscript.

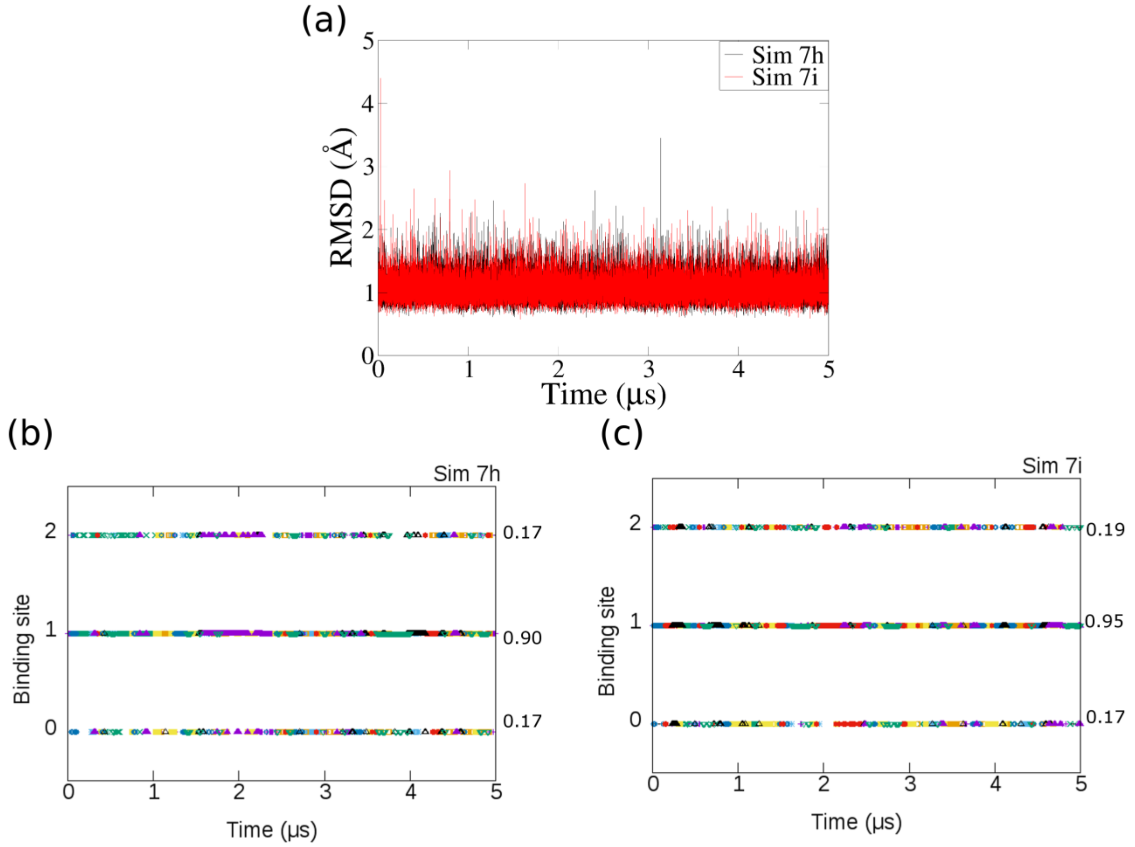


**Figure S10**. Backbone RMSD (a) and plots for monitoring cation movement (b-c) in the simulations of antiparallel d[GG]$_4$ G-stem from PDB 148D carried out in $Na^+$ ions and OPC water model (Simulations 7h and 7i). (a) Backbone RMSD plots showed that G-stem was stable in both the Simulations 7h and 7i, shown in black and red, respectively. Cation binding plots of the simulations (b) 7h and (c) 7i showed that there were very frequent ion-exchange movements between the G-stem and the solvent. The cation binding Site 1 corresponds to the typical position of cations in the channel of GQ between the two quartets. Sites 0 and 2 stand for the positions over the 5′-quartet and below the 3′-quartet, respectively. In the panels b and c, the numbers on the right show occupancies of the respective sites. For full details, see the legend to Figure 4 of the Main manuscript.

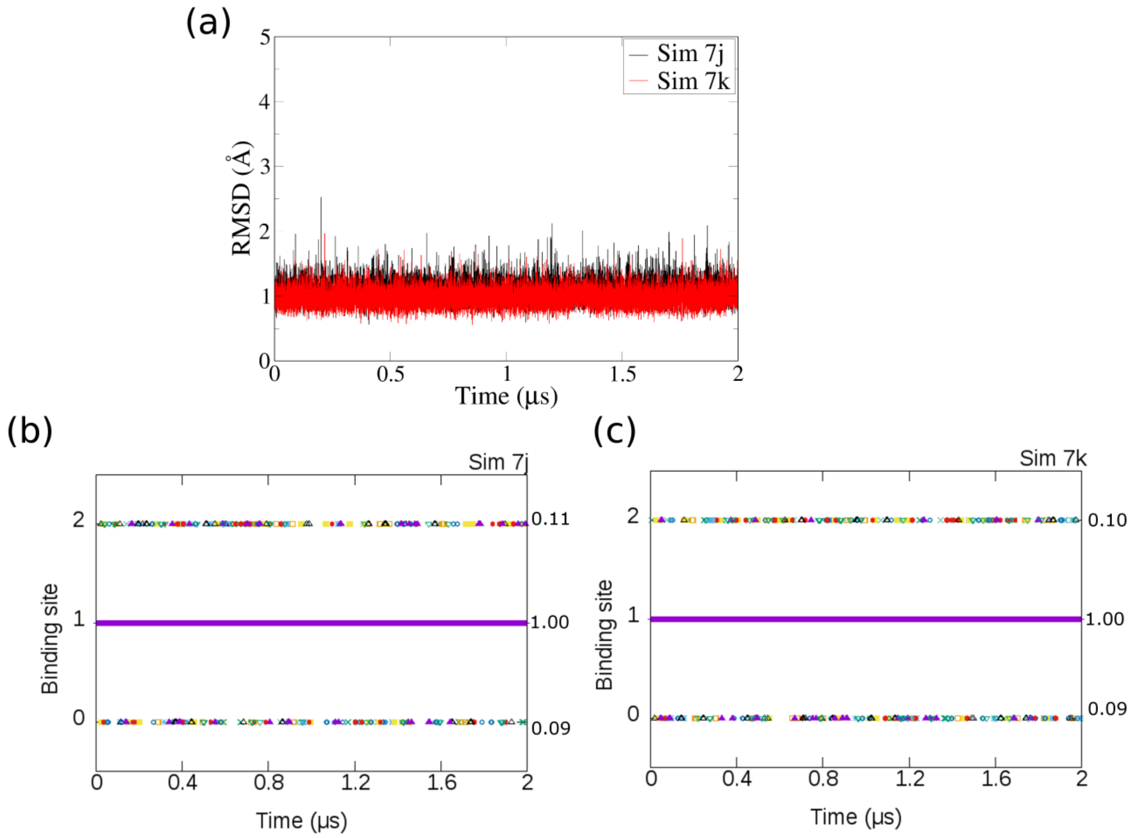


**Figure S11**. Backbone RMSD (a) and plots for monitoring cation movement (b-c) in the simulations of antiparallel d[GG]$_4$ G-stem from PDB 148D carried out in $K^+$ ions and OPC water model (Simulations 7j and 7k). (a) Backbone RMSD plots showed that G-stem was stable in both the Simulations 7j and 7k, shown in black and red, respectively. Cation binding plots of the simulations (b) 7j and (c) 7k showed that there was no cation exchange movement and one cation stabilized the G-stem. The cation binding Site 1 corresponds to the typical position of cations in the channel of GQ between the two quartets. Sites 0 and 2 stand for the positions over the 5′-quartet and below the 3′-quartet, respectively. In the panels b and c, the numbers on the right show occupancies of the respective sites. For full details, see the legend to Figure 4 of the Main manuscript.

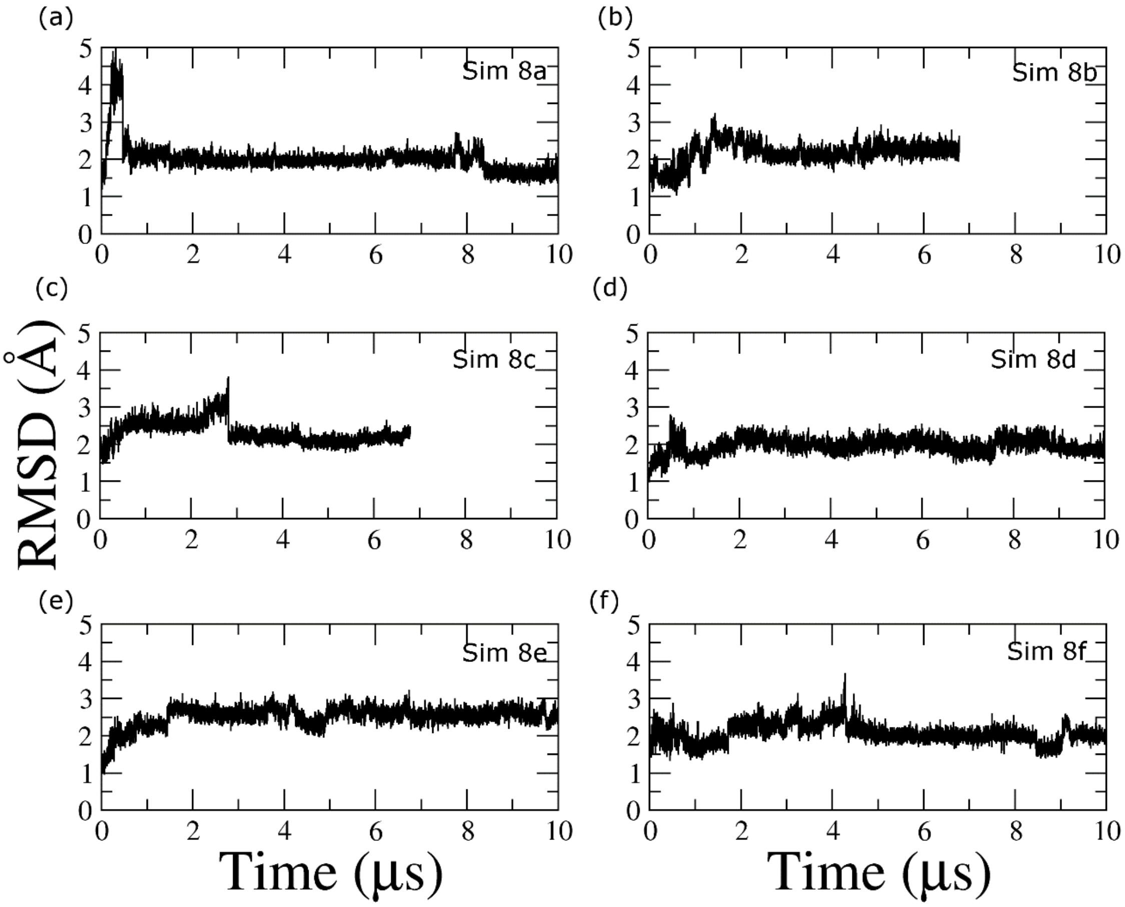


**Figure S12**. Backbone RMSD plots of dimer of two-quartet RNA GQs in the Simulations (a) 8a, (b) 8b, (c) 8c, (d) 8d, (e) 8e and (f) 8f carried out in $K^+$ ions placed at different sites in the starting structure and SPC/E water model. The GQ dimer did not show any major RMSD fluctuations and was maintained in all the simulations.

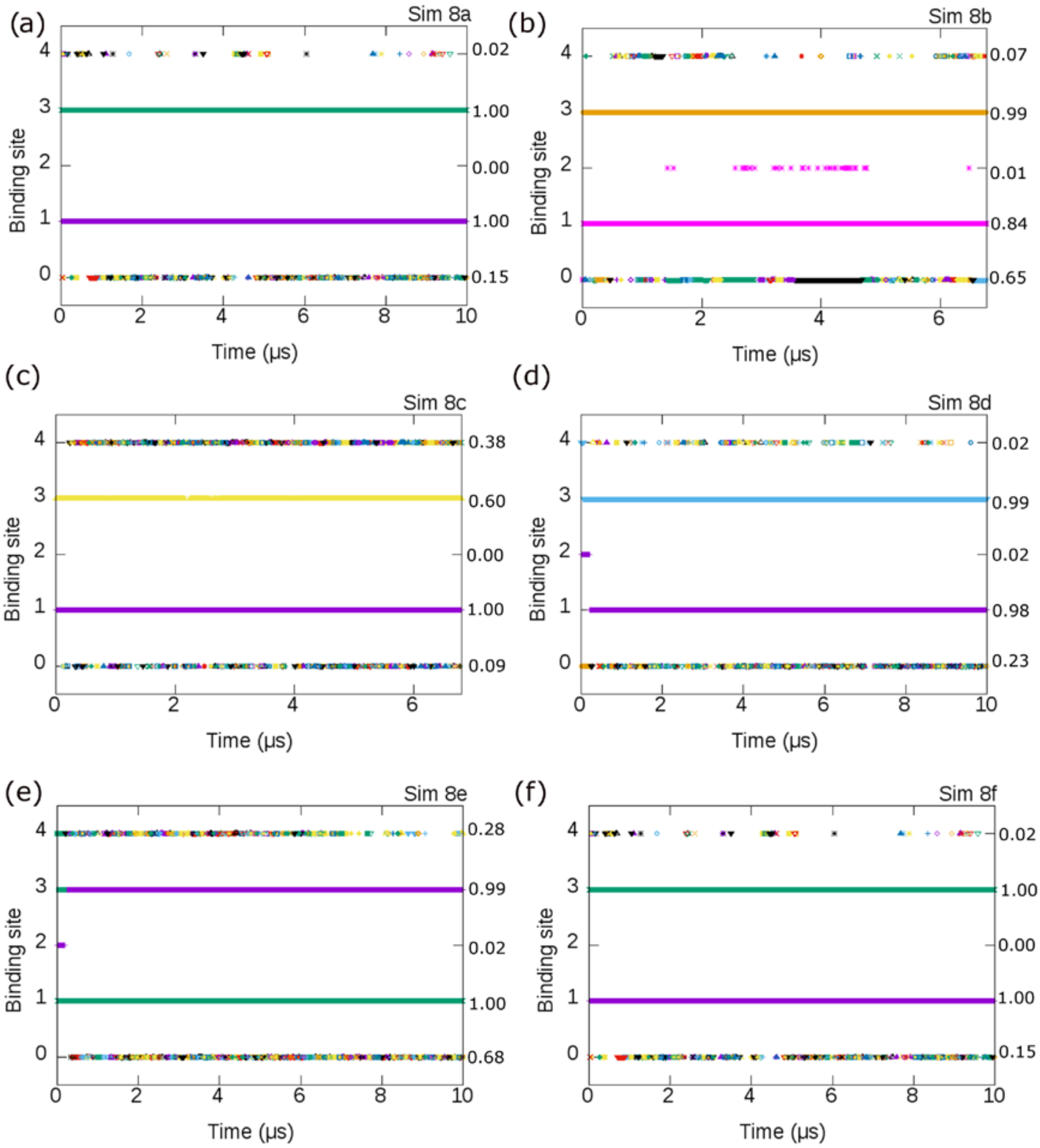


**Figure S13**. Cation binding to the G-stem in the Simulations 8a - 8f of two-quartet RNA dimer in the SPC/E water model. $K^+$ ions were placed at different sites in the starting structure (see Table S1 for details). Cation binding is shown in the Simulations (a) 8a, (b) 8b, (c) 8c, (d) 8d, (e) 8e and (f) 8f. The numbers on the right show occupancies of the respective sites. Site 2 was vacant for the majority of the time in all the simulations. The apparent transient simultaneous occupancy of Sites 1 and 2 in the Simulation 8b (b) means transient cation binding in-plane of the second quartet. For full details, see the legend to Figure S3.

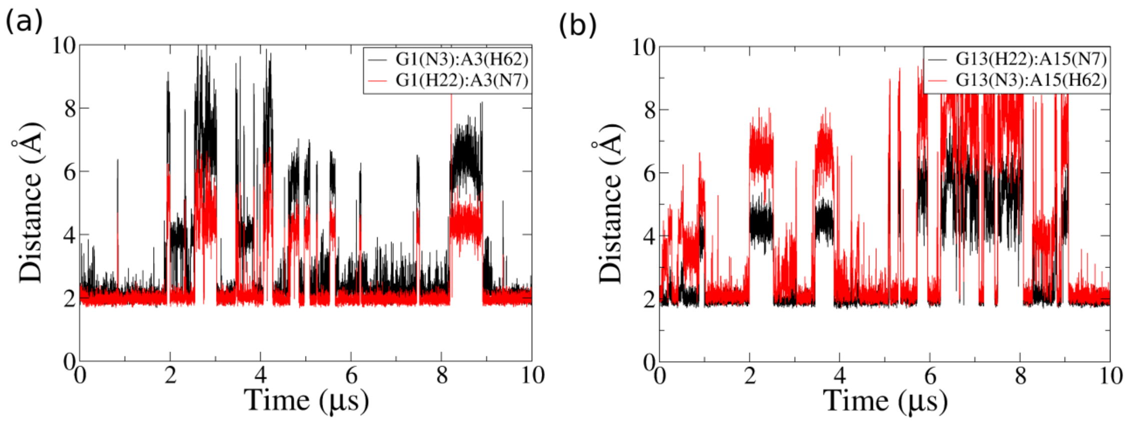


**Figure S14**. Distance plots showing the interactions of loop nucleotides with the G-stem of 2RQJ in the Simulation 8d carried out in the presence of $K^+$ ions and SPC/E water model. (a) A3 formed base pair with G1 and (b) A15 formed base pair with G13 during the simulation.

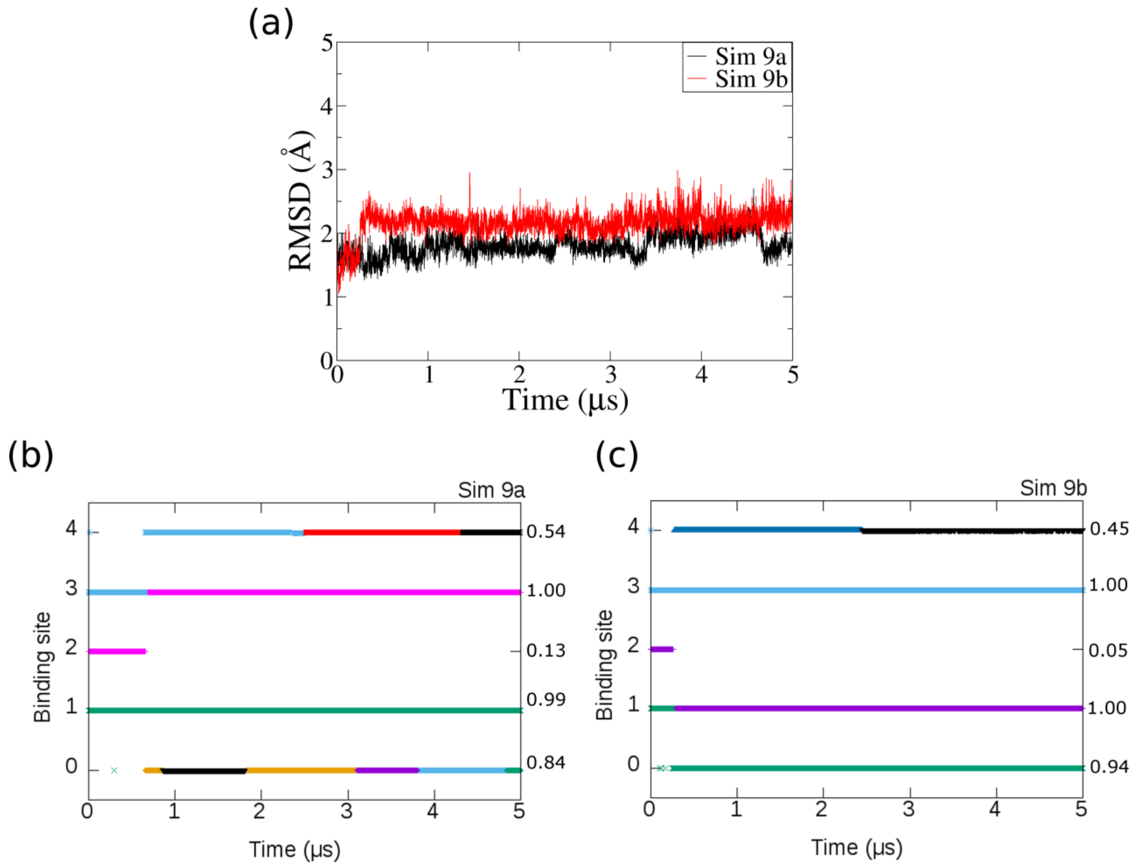


**Figure S15**. Backbone RMSDs (a) and plots for monitoring cation binding (b - c) in the simulations of two-quartet RNA GQ dimer represented by PDB 2RQJ (Simulations 9a and 9b). The simulations were carried out using $Na^+$ ions and SPC/E water model. Panel a shows backbone RMSD of the GQ in Simulations 9a and 9b in black and red, respectively. No major fluctuation in backbone RMSD values was observed indicating the stability of the simulations. In the panels b and c, the numbers on the right show occupancies of the respective sites. Site 2 was vacant for the majority of the time in both the simulations. For full details, see the legend to Figure S3.

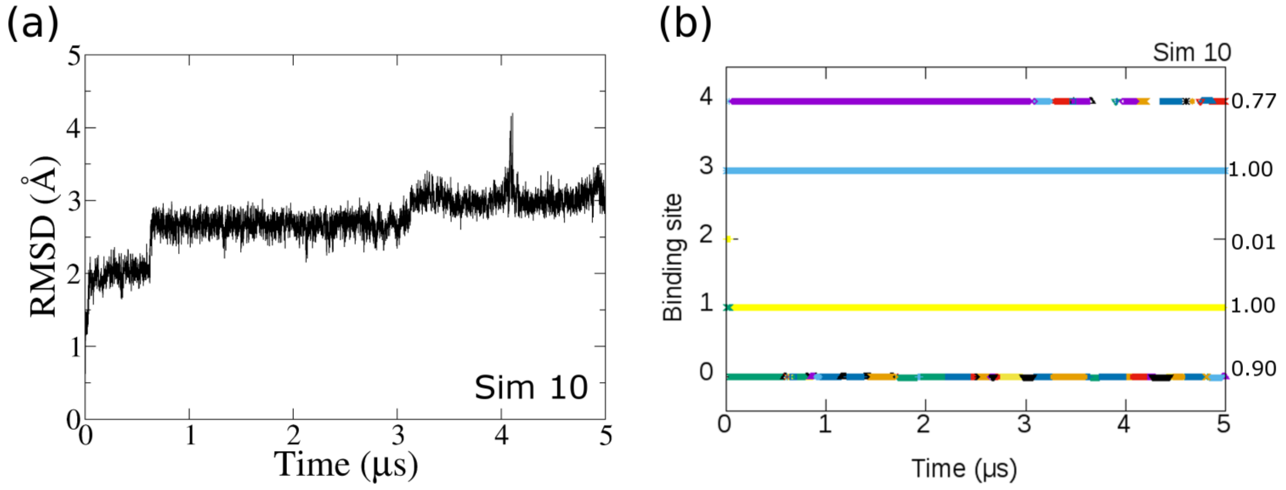


**Figure S16**. Backbone RMSD and plot for monitoring cation binding in the simulation of two-quartet RNA GQ dimer represented by PDB 2RQJ (Simulation 10). The simulation was carried out using $Na^{+}$ ions and OPC water model. Panel (a) shows backbone RMSD of the GQ. No major fluctuation in backbone RMSD was observed indicating the stability of the simulation. The slight RMSD movements around 0.8 and 4.1 μs are due to the loop rearrangements. Panel (b) shows cation movement within the GQ. Just after the start of the simulation, the "yellow" cation moved from Site 2 to Site 1 pushing the original Site 1 "green" ion over the 5′-quartet. No further cation exchange was observed during the simulation. Distinct ions are represented by different colors and symbols, and any change in the color/symbol indicates an exchange of an ion in the site. The numbers on the right show occupancies of the respective sites. For full details, see the legend to Figure S3.

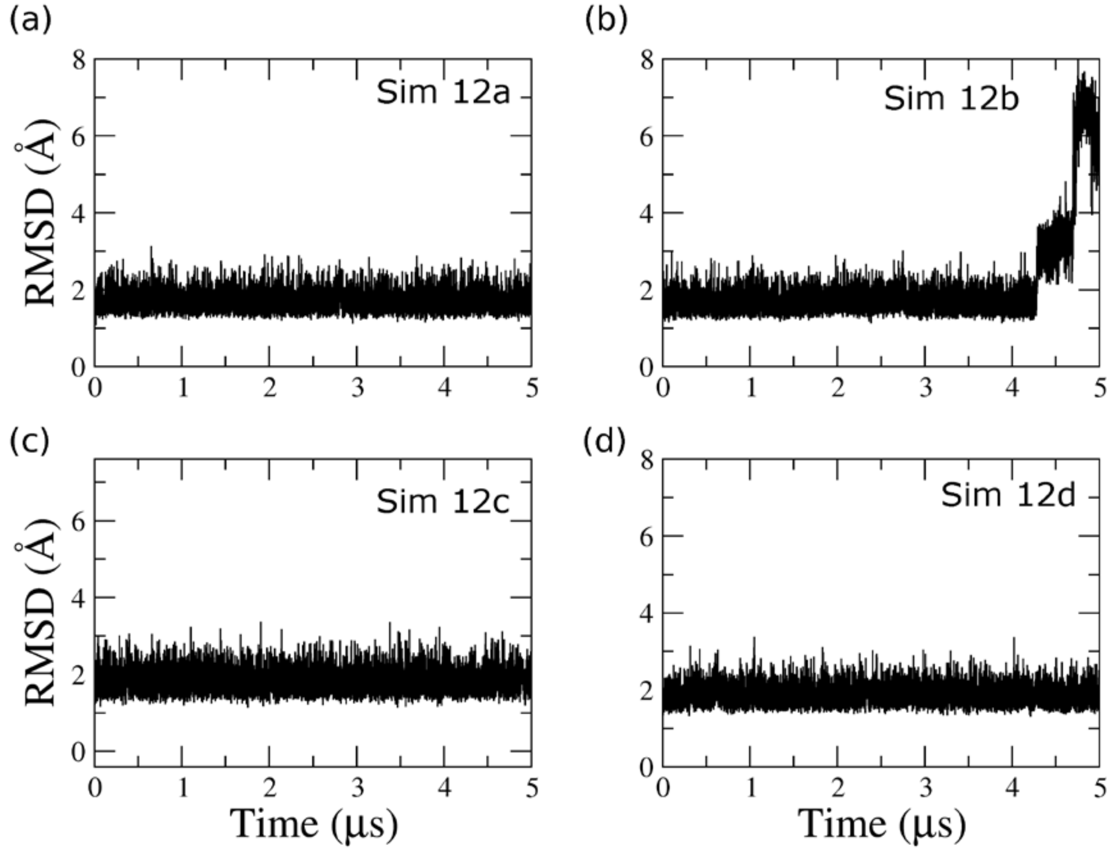


**Figure S17**. Backbone RMSD of two-quartet RNA GQ in the Simulations (a) 12a, (b) 12b, (c) 12c and (d)12d. The simulations were carried out in $Na^+$ ions and SPC/E water model. The two-quartet RNA GQ unfolded in the Simulation 12b at 4.27 μs shown in the panel b and was stable in the other three simulations shown in the panels a, c and d.

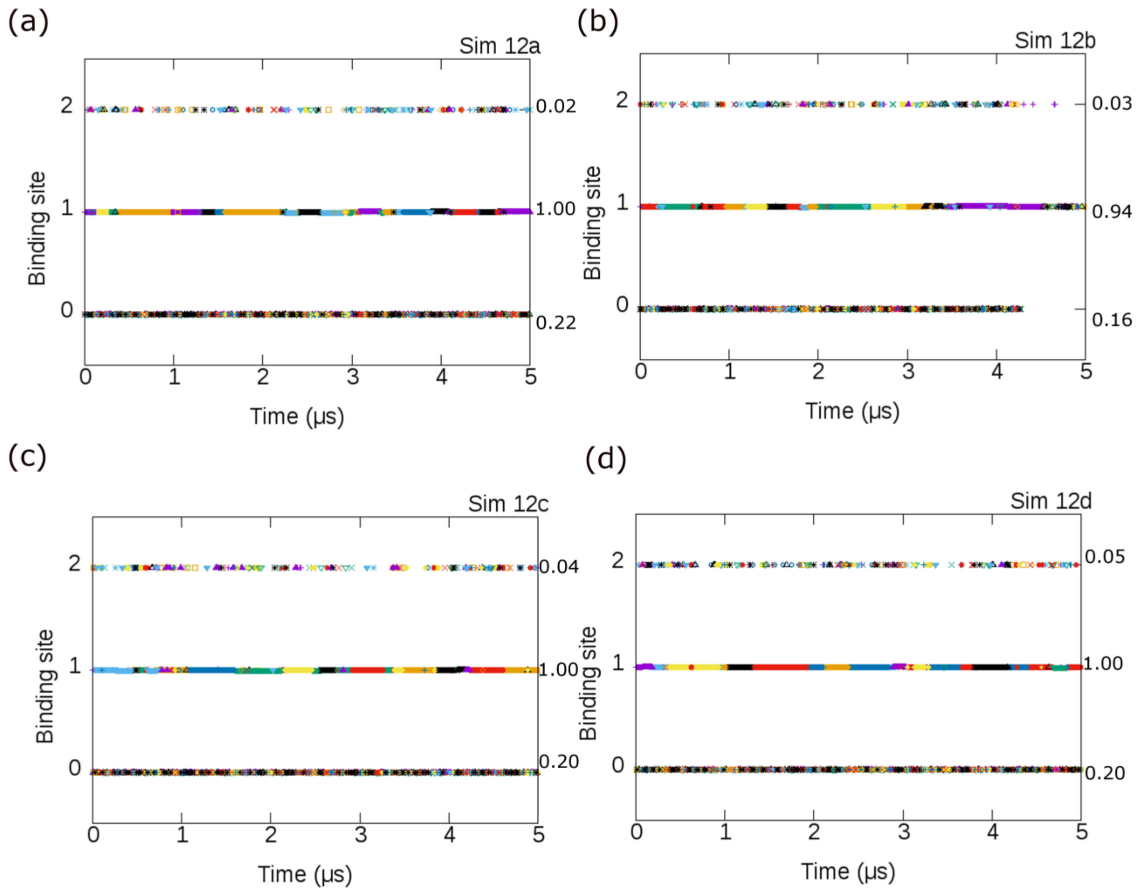


**Figure S18**. Movement of $Na^+$ ions in two-quartet RNA GQ (panels a - d) in the Simulations 12a-12d carried out in SPC/E water model. The GQ unfolded in the Simulation 12b at 4.27 µs. The cation binding Site 1 corresponds to the typical position of cations in the channel of GQ between the two quartets. Sites 0 and 2 stand for the positions over the 5′-quartet and below the 3′-quartet, respectively. Frequent ion exchanges between the channel and the solvent were observed in all the simulations. The numbers on the right show occupancies of the respective sites. For full details, see the legend to Figure 4 of the Main manuscript.

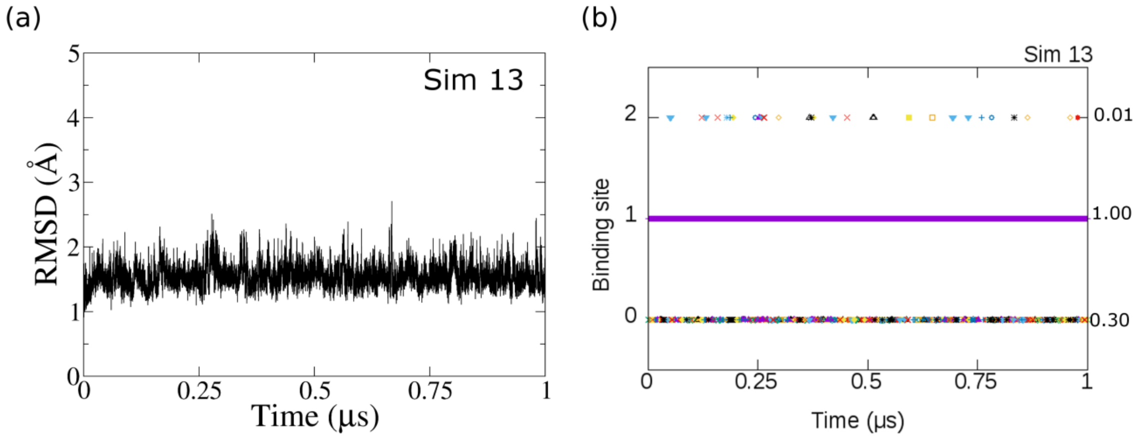


**Figure S19**. (a) Backbone RMSD of GQ and (b) plot showing cation binding in the GQ in the Simulation 13. The simulation of two quartet RNA GQ from PDB 2RQJ without 3′-flanking base was carried out in $K^+$ ion with the SPC/E water model. The backbone RMSD plot shows that GQ was stable as no significant deviation from the starting structure was observed. The cation binding Site 1 corresponds to the typical position of cations in the channel of GQ between the two quartets. Sites 0 and 2 stand for the positions over the 5′-quartet and below the 3′-quartet, respectively. There was no exchange of channel $K^+$ ion with the solvent as is evident from the graph in panel b. In panel b, the numbers on the right show occupancies of the respective sites. For full details, see the legend to Figure 4 of the Main manuscript.

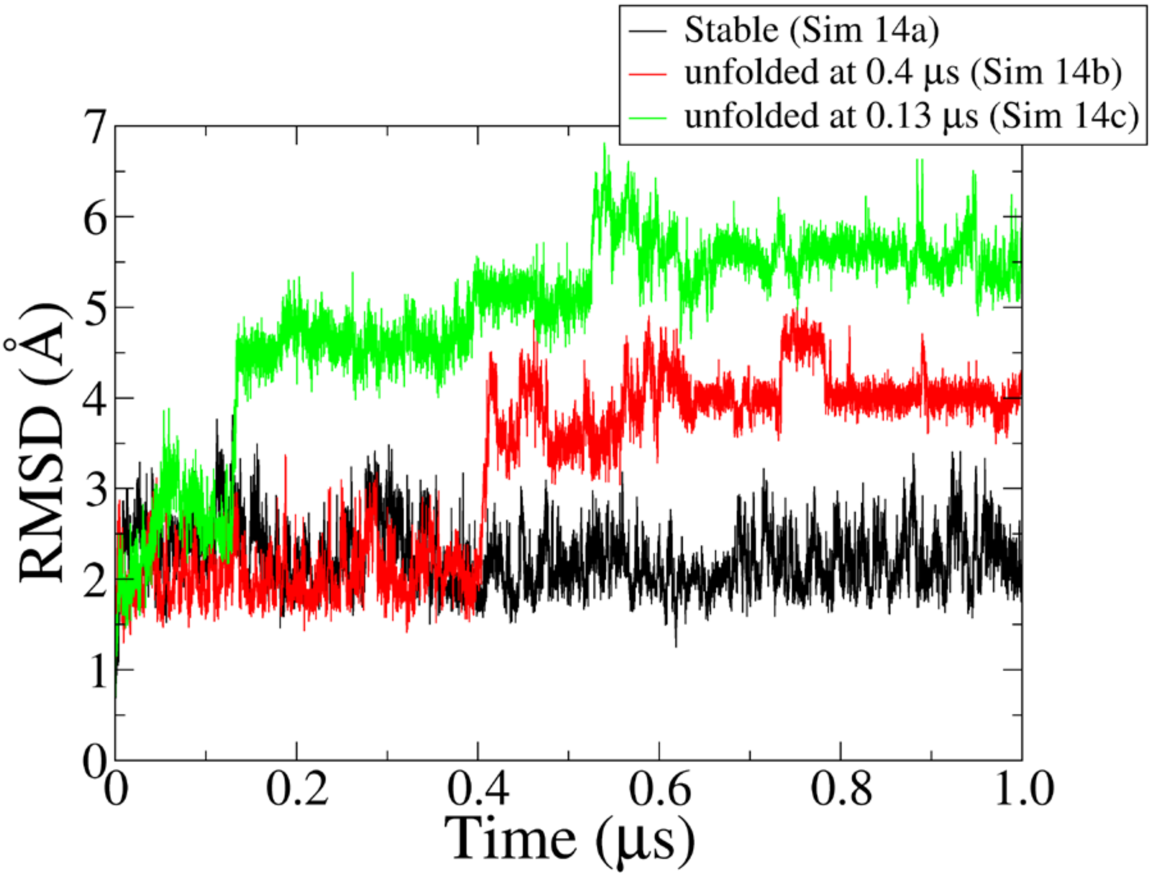


**Figure S20**. Backbone RMSD of two-quartet RNA GQ in $Na^+$ ions and OPC water model in the Simulations 14a - 14c. The GQ was stable in the Simulation 14a shown in black. The GQ unfolded at 0.4 μs and 0.13 μs in the Simulations 14b (shown in red) and 14c (shown in green), respectively.

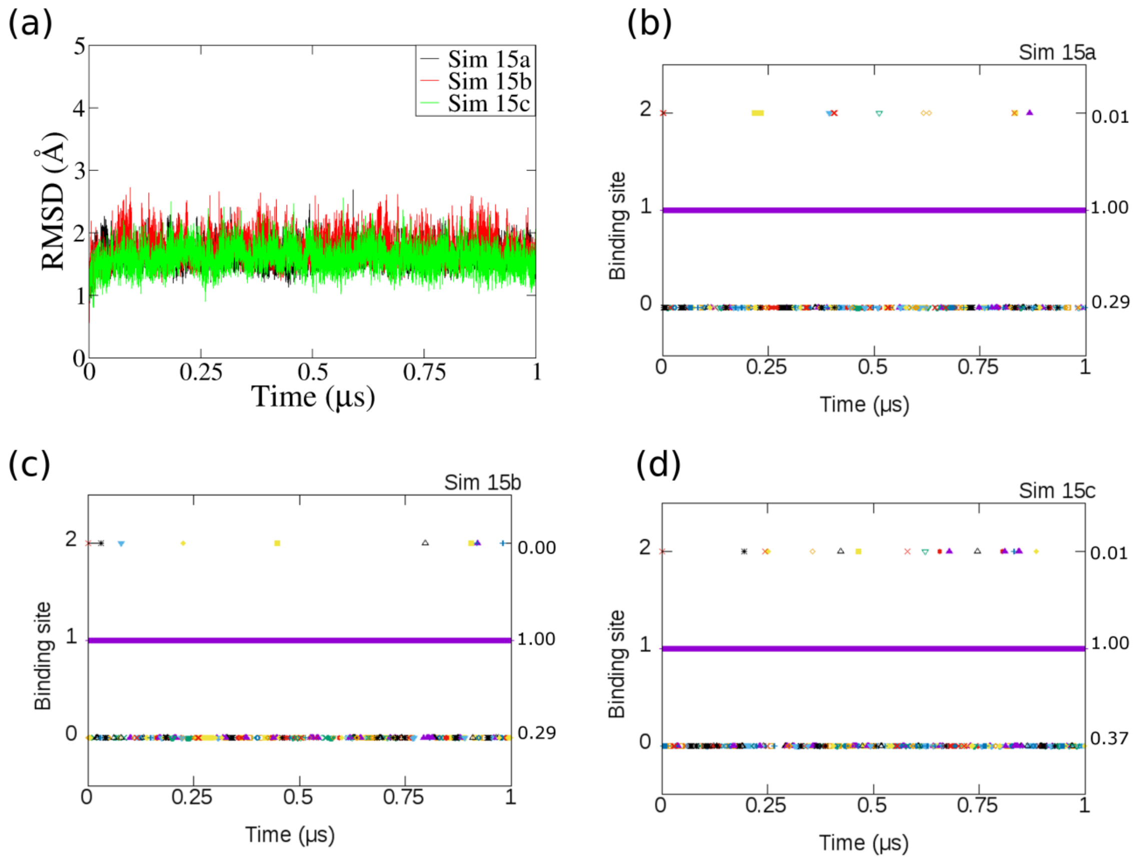


**Figure S21**. (a) Backbone RMSD of GQ, (b), (c) and (d) movements of cations in the GQ in the Simulations 15a-15c. The Simulations 15a-15c of two quartet RNA GQ from PDB 2RQJ without 3′-flanking base were carried out in $K^+$ ion with OPC water model. The backbone RMSD plot shows that the GQ was stable. In panels b, c and d, the cation binding Site 1 correspond to the typical position of cations in the channel of GQ between the two quartets. Sites 0 and 2 stand for the positions over the 5′-quartet and below the 3′-quartet, respectively. There was no exchange of channel $K^+$ ion with the solvent in any of the three simulations as is evident from the graphs in panels b, c and d; each color/symbol represents different cation in the simulations. The numbers on the right show occupancies of the respective sites. For full details, see the legend to Figure 4 of the Main manuscript.

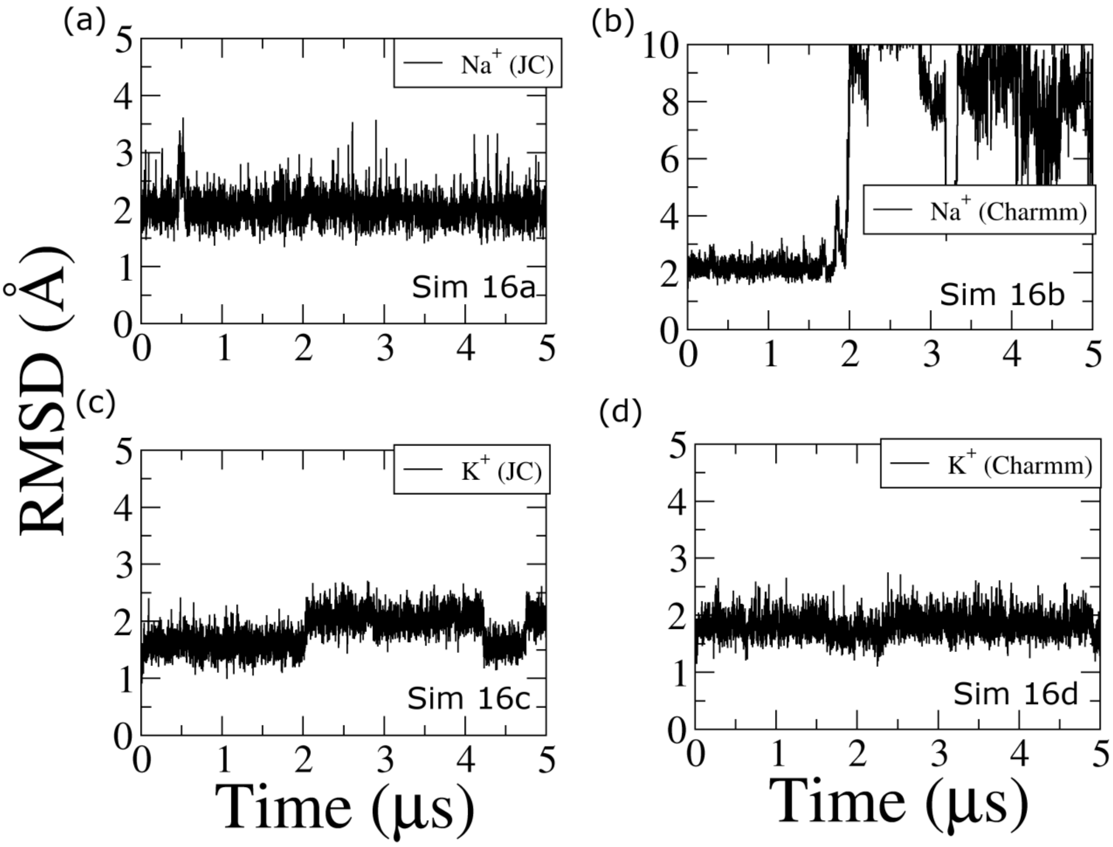


**Figure S22**. Backbone RMSD of two-quartet RNA GQ represented by a monomer of 2RQJ in the Simulations 16a - 16d. The simulations were carried out in TIP4P-D water and (a) Joung and Cheatham (JC) $Na^+$, (b) CHARMM22 $Na^+$, (c) JC $K^+$ and (d) CHARMM22 $K^+$ ions. The GQ was stable in JC $Na^+$, JC $K^+$ and CHARMM22 $K^+$ ions (Simulations 16a, 16c and 16d). GQ completely unfolded in the simulation in CHARMM22 $Na^+$ ions (Simulation 16b).

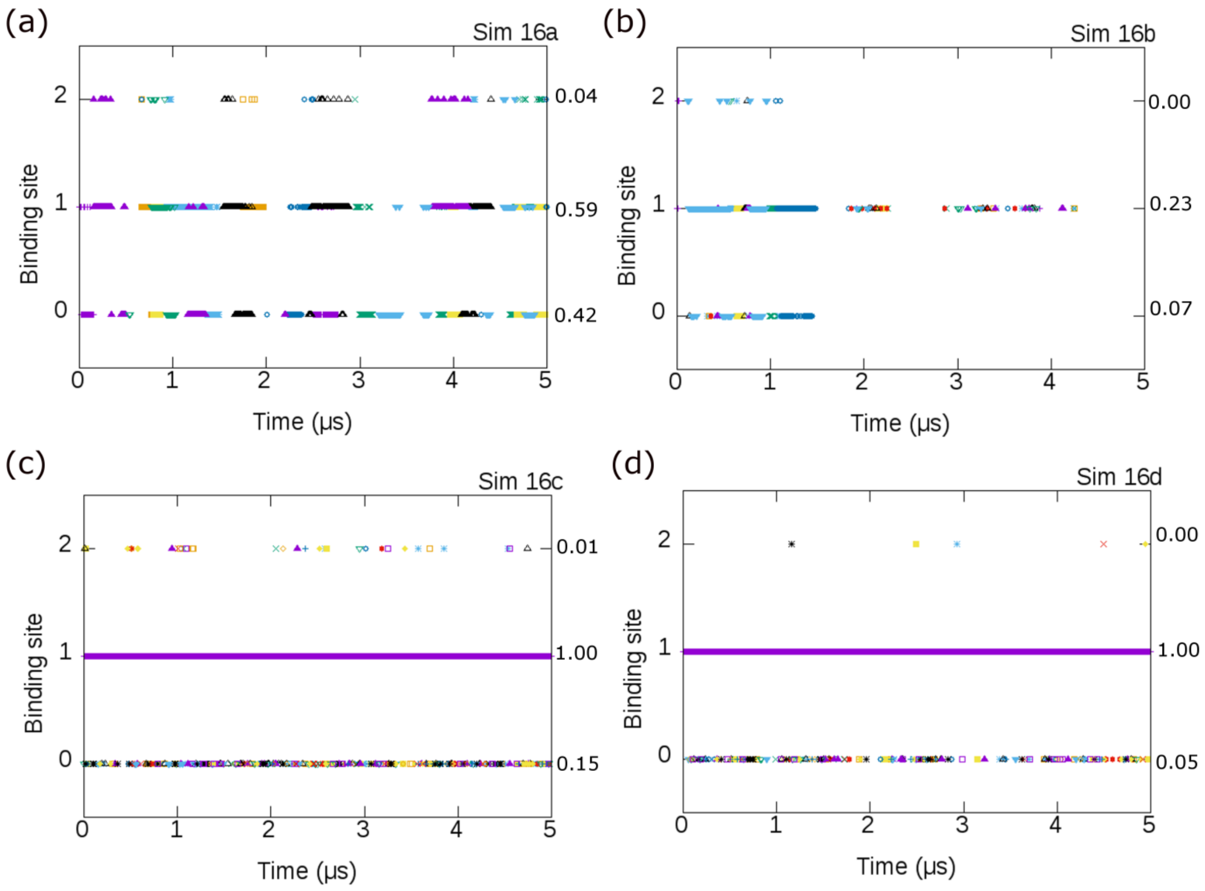


**Figure S23**. Plots monitoring cation binding in GQ in the simulations of 2RQJ monomer in the Simulations 16a - 16d. The simulations were carried out in TIP4P-D water and (a) JC $Na^+$, (b) CHARMM22 $Na^+$, (c) JC $K^+$ and (d) CHARMM22 $K^+$ ions. The cation binding Site 1 corresponds to the typical position of cations in the channel of GQ between the two quartets. Sites 0 and 2 stand for the positions over the 5′-quartet and below the 3′-quartet, respectively. Multiple cation exchanges were observed in the simulation in JC $Na^+$ ions while there was no cation exchange in both $K^+$ simulations. The GQ completely unfolded in the simulation in CHARMM22 $Na^+$ ions (Simulation 16b). Each color/symbol in the panels represents different cation in the simulations. The numbers on the right show occupancies of the respective sites. For full details, see the legend to Figure 4 of the Main manuscript.

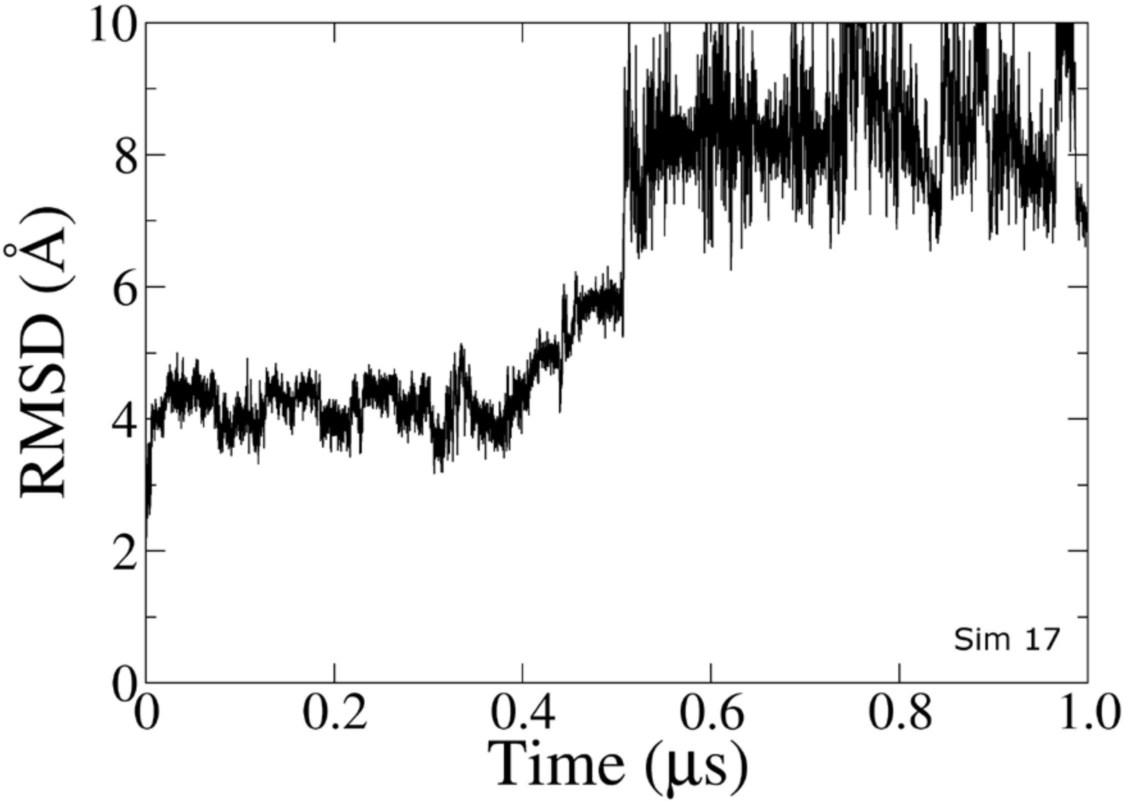


**Figure S24**. Backbone RMSD of two-quartet DNA GQ with adenine loops in the Simulation 17. The simulation was carried out with $K^+$ ions and SPC/E water model. The significant change in RMSD at 450 ns indicates unfolding of the GQ.

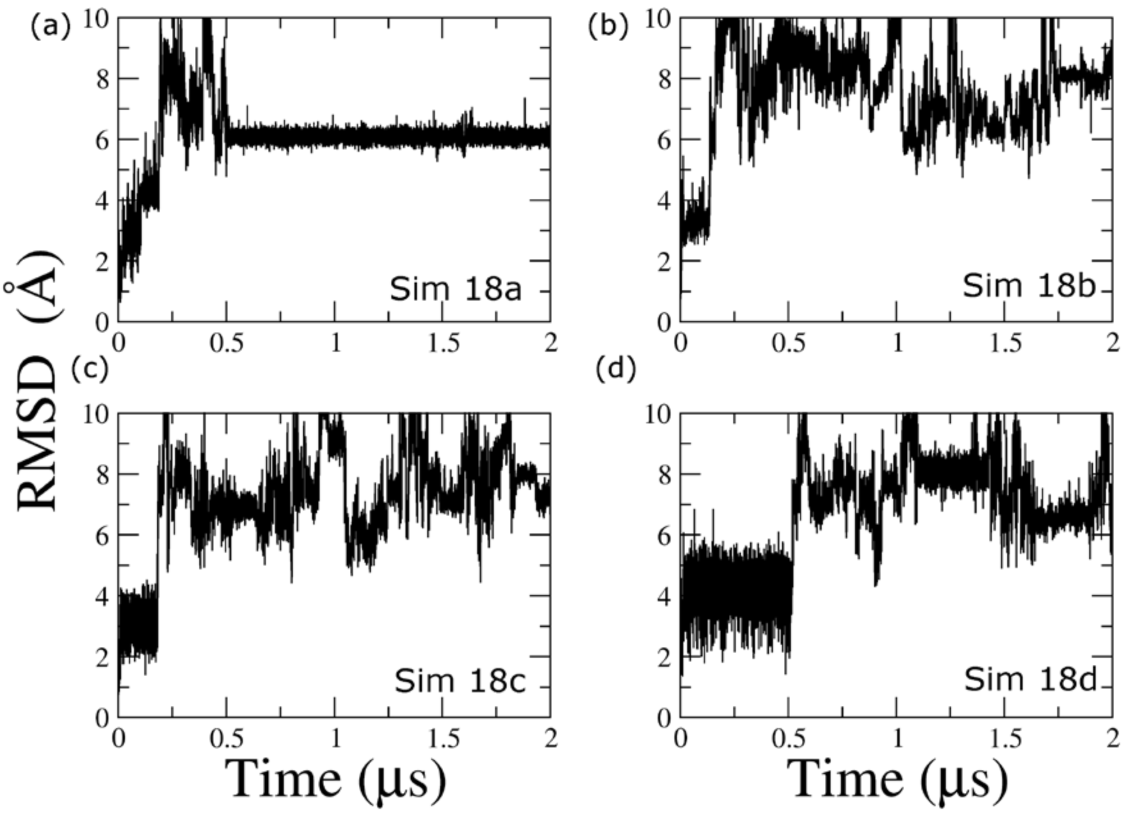


**Figure S25**. Backbone RMSD of antiparallel r[GG]$_4$ model G-stem in the Simulations 18a-18d. The simulations were carried out in (a), (b) $Na^+$ or (c), (d) $K^+$ and in SPC/E water model. The G-stem unfolded in all the simulations which is evident from significant RMSD changes at 10 ns, 50 ns, 180 ns and 510 ns in the Simulations 18a-18d, respectively.

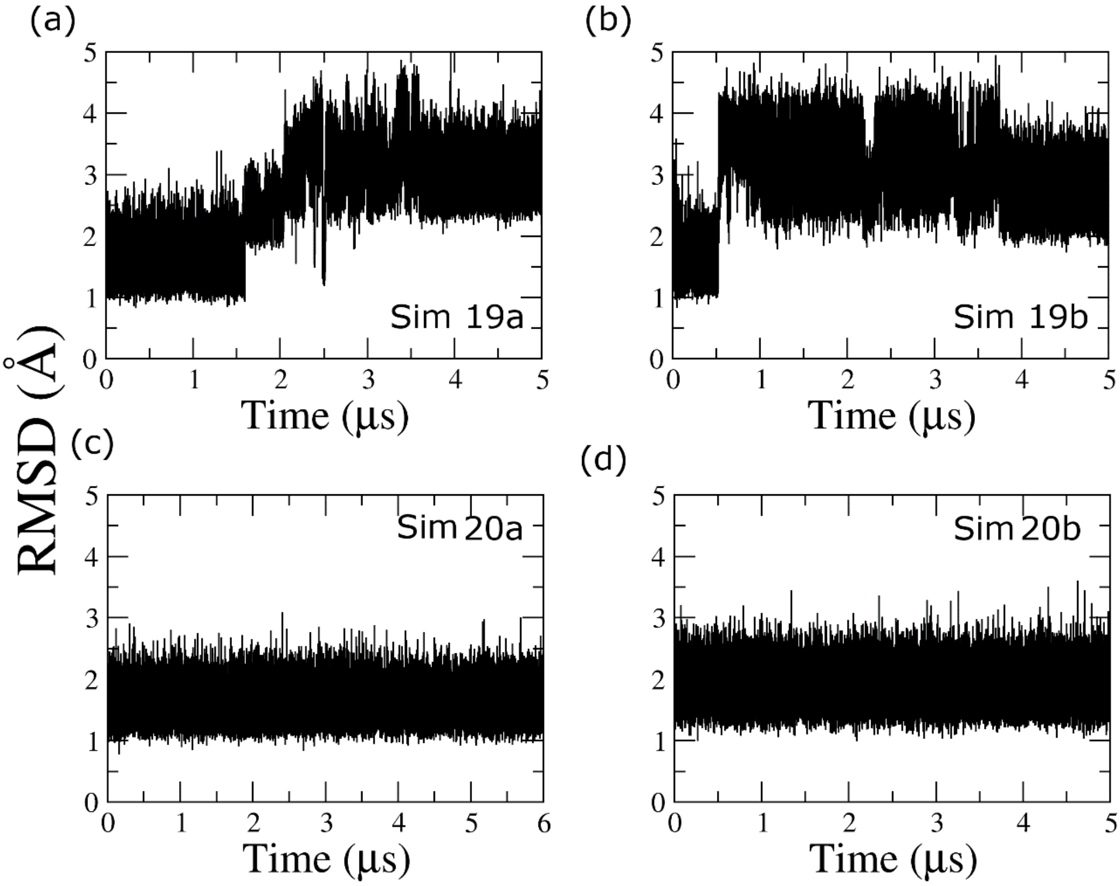


**Figure S26**. Backbone RMSD of all *anti* parallel-stranded d[GGG]$_4$ G-stem in the Simulations 19a-19b and 20a-20b. The simulations were carried out in (a), (b) $Na^+$ (Simulations 19a-19b) or (c), (d) $K^+$ (Simulations 20a-20b) and in SPC/E water model. The significant change in RMSD indicates perturbation of the G-stem by vertical strand slippage. Strand slippage was observed in both the simulations in $Na^+$ and is evident from significant RMSD changes at 1.6 and 0.53 μs in panels a and b. The G-stem was stable in the simulations in $K^+$ ions shown in the panels c and d.

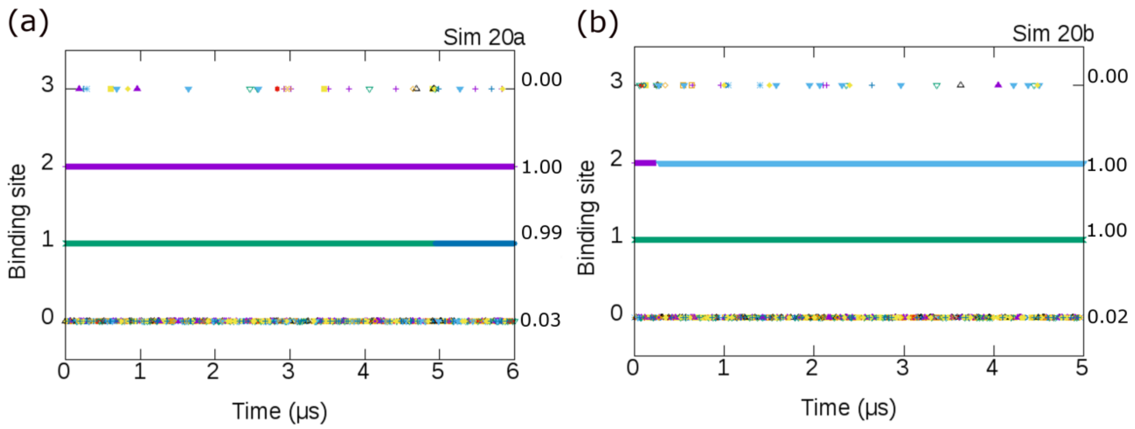


**Figure S27**. Plots monitoring cation binding in the all *anti* parallel-stranded d[GGG]$_4$ G-stem in the Simulations 20a-20b. The simulations were carried out in $K^+$ and in the SPC/E water model. The cation binding sites 1 and 2 correspond to the typical position of cations in the channel of GQ between the two quartets. Sites 0 and 3 stand for the positions over the 5′-quartet and below the 3′-quartet, respectively. See panel (g) of Figure 1 for the representation of these sites. Distinct ions are represented by different colors and symbols, and any change in the color/symbol indicates an exchange of an ion in the site. The numbers on the right show occupancies of the respective sites. Two $K^+$ ions stabilized the three-quartet G-stem. A channel cation was lost in the Simulation 20a at 4.8 μs followed, however, by quick entry of new cation from the bulk. In the Simulation 20b, a cation exchange event was observed at 0.27 μs.

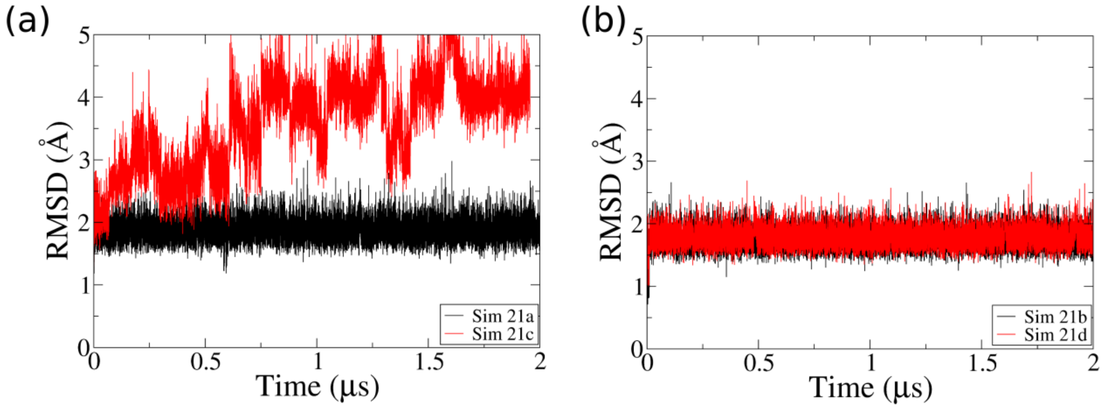


**Figure S28**. Backbone RMSD of antiparallel d[GGG]$_4$ derived from PDB 143D in the Simulations 21a-21d. The simulations were carried out in $Na^+$ or $K^+$ ions and in SPC/E or OPC water model. In the panel (a) $Na^+$ simulations in SPC/E (Simulation 21a) and OPC (Simulation 21c) water models are shown in black and red, respectively. In the panel (b) $K^+$ simulations in SPC/E (Simulation 21b) and OPC (Simulation 21d) water models are shown in black and red, respectively. The significant change in RMSD in panel (a) shown in red indicates quartet disruption at 0.185 μs in the $Na^+$ ions and OPC water model (Simulation 21c).

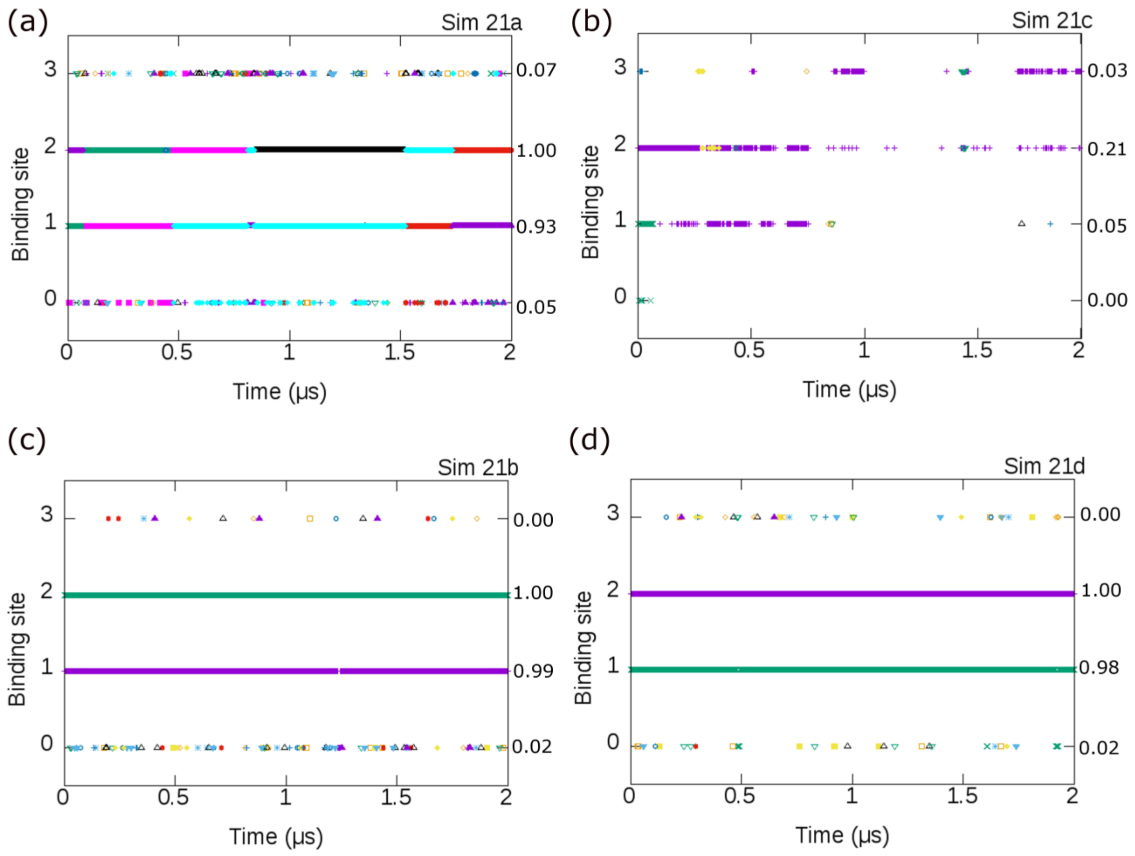


**Figure S29**. Plots monitoring cation binding in the antiparallel d[GGG]$_4$ G-stem derived from PDB 143D in the Simulations 21a-21d. The simulations were carried out in (a) $Na^+$ and SPC/E, (b) $Na^+$ and OPC, (c) $K^+$ and SPC/E and (d) $K^+$ and OPC conditions. The numbers on the right show occupancies of the respective sites. Two $Na^+$ or $K^+$ ions stabilized the three-quartet G-stem. Multiple cation exchange events were observed in the Simulation 21a and two $Na^+$ ions stabilized the G-stem as shown in panel a. The first quartet was disrupted in the Simulation 21c at 0.185 μs and cation binding is not of much significance in such a case. Panels c and d show that two $K^+$ ions were stably retained in the simulations in both SPC/E and OPC water model. See legend to Figure S27 for details.

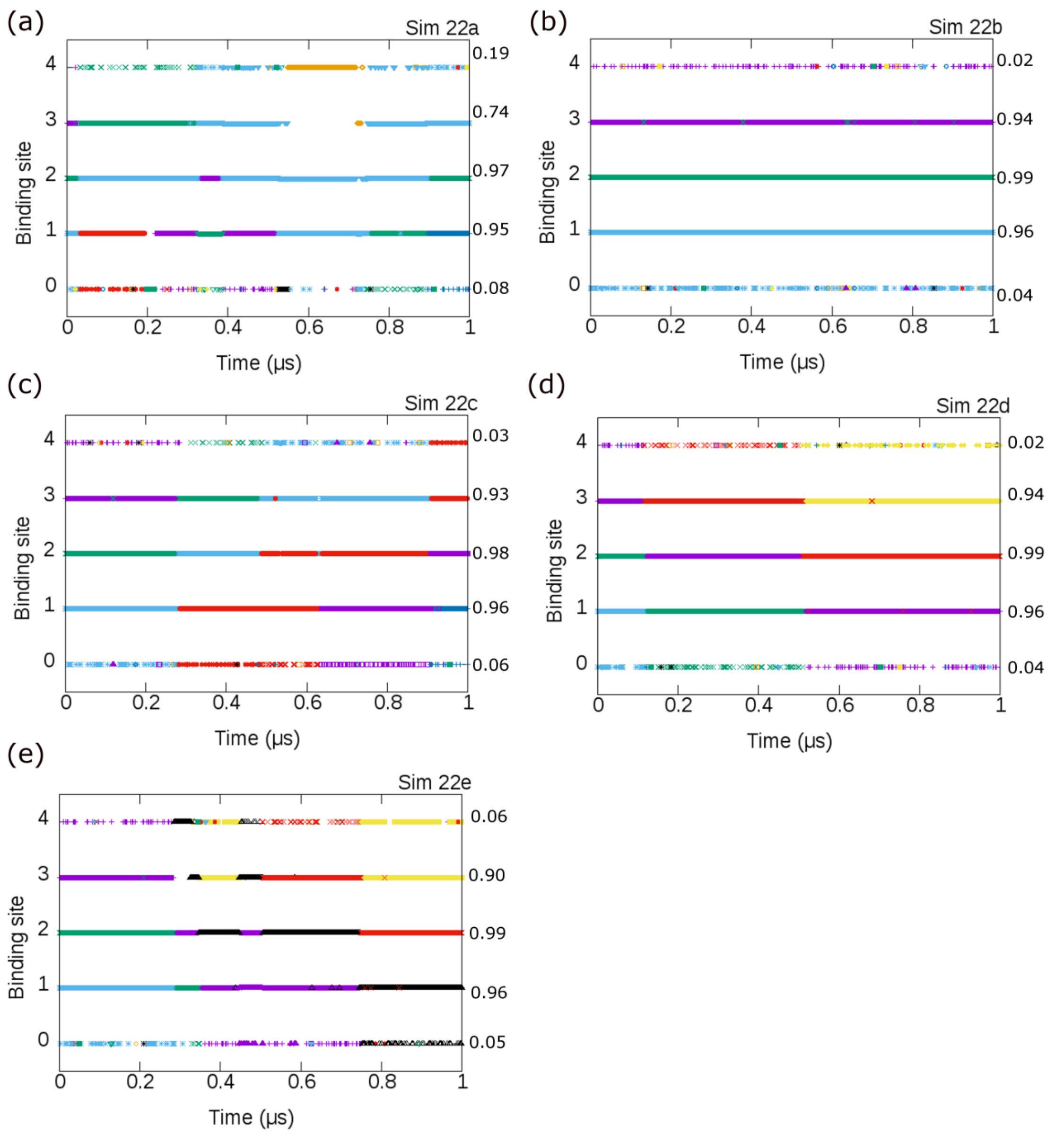


**Figure S30**. Plots monitoring cation binding in the parallel-stranded all-*anti* d[GGGG]$_4$ G-stem in the Simulations 22a - 22e. Simulations (a) 22a, (b) 22b, (c) 22c, (d) 22d and (e) 22e were carried out in $Na^+$ ions and SPC/E water model. Distinct ions are represented by different colors and symbols, and any change in the color/symbol indicates an exchange of an ion in the site. The numbers on the right show occupancies of the respective sites. See legend to Figure S3 for details. Multiple cation exchange events were observed in the Simulations 22a (panel a) and 22c-22e (panel c - e). Apparent simultaneous occurrence of a cation at two sites means the cation is in-plane of the quartet between the two sites; see the explanation in the “Visualization of cation binding sites” paragraph at the beginning of the Supporting Information. No cation exchange was observed in the Simulation 22b (panel b).

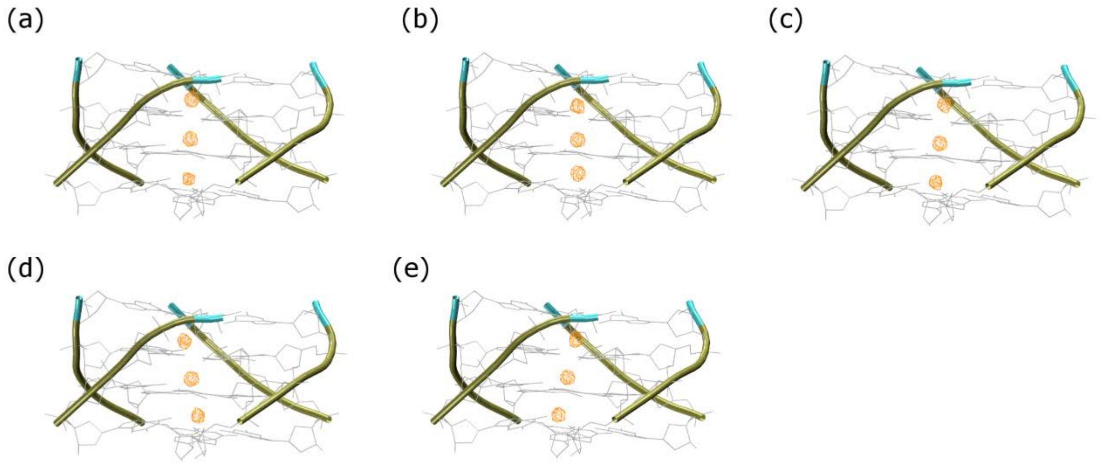


**Figure S31**. Cation binding sites observed in the atomistic simulations of parallel-stranded all *anti* d[GGGG]$_4$ four-quartet G-stem in the Simulations (a) 22a, (b) 22b, (c) 22c, (d) 22d and (e) 22e. The simulations were carried out in $Na^+$ ions and SPC/E water model. Three $Na^+$ ions stabilized the G-stem in the simulations and were present close or closer to the quartet plane at the 3'-end side of the ion. The $Na^+$ binding sites are shown in orange. The backbone is shown in tan tube and the guanine nucleotides are shown as silver sticks. The backbone of the 5′-end of the strands is shown in cyan.

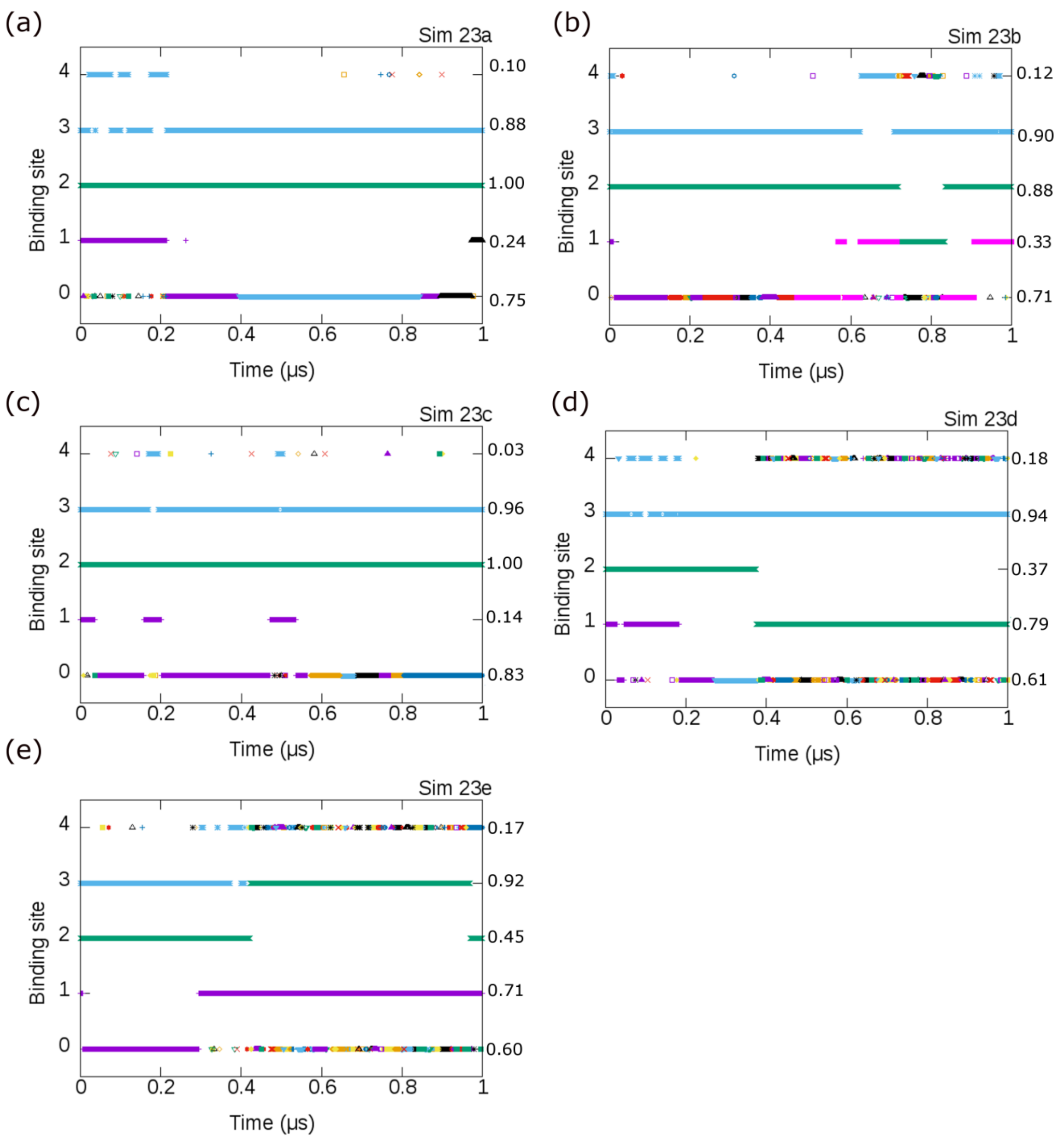


**Figure S32**. Plots monitoring cation binding in the parallel-stranded all *anti* d[GGGG]$_4$ four-quartet G-stem in the Simulations 23a - 23e. Simulations (a) 23a, (b) 23b, (c) 23c, (d) 23d and (e) 23e were carried out in $K^+$ ions and SPC/E water model. Distinct ions are represented by different colors and symbols, and any change in the color/symbol indicates an exchange of an ion in the site. The numbers on the right show occupancies of the respective sites. See legend to Figure S3 for details.

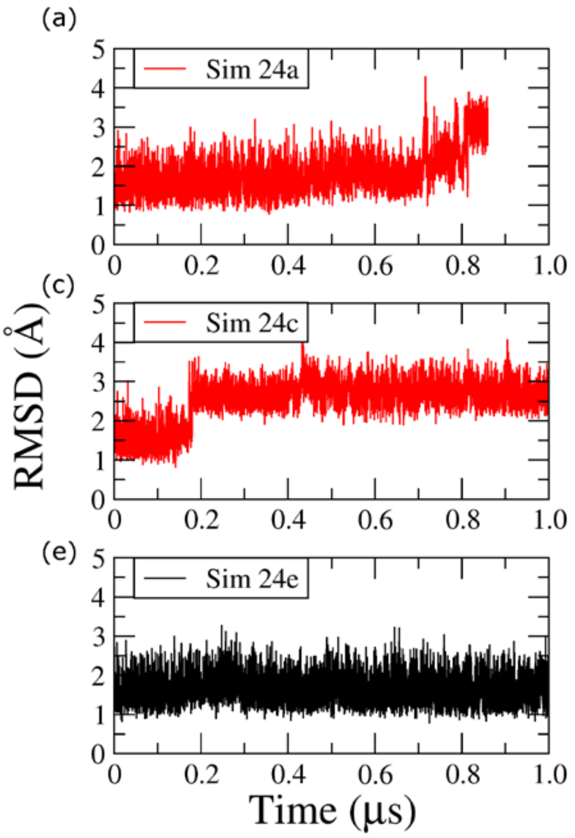


**Figure S33**. Backbone RMSD of parallel-stranded all *anti* d[GGGG]$_4$ four-quartet G-stem in the Simulations (a) 24a, (b) 24b, (c) 24c, (d) 24d and (e) 24e. Five independent simulations were carried out in $Na^+$ ions and OPC water model. A significant change in the RMSD values indicated significant perturbation of the G-stem, as described in the main text. The RMSD plots in stable and perturbed G-stem simulations are shown in black and red, respectively. Strand slippage was observed in three out of five simulations.

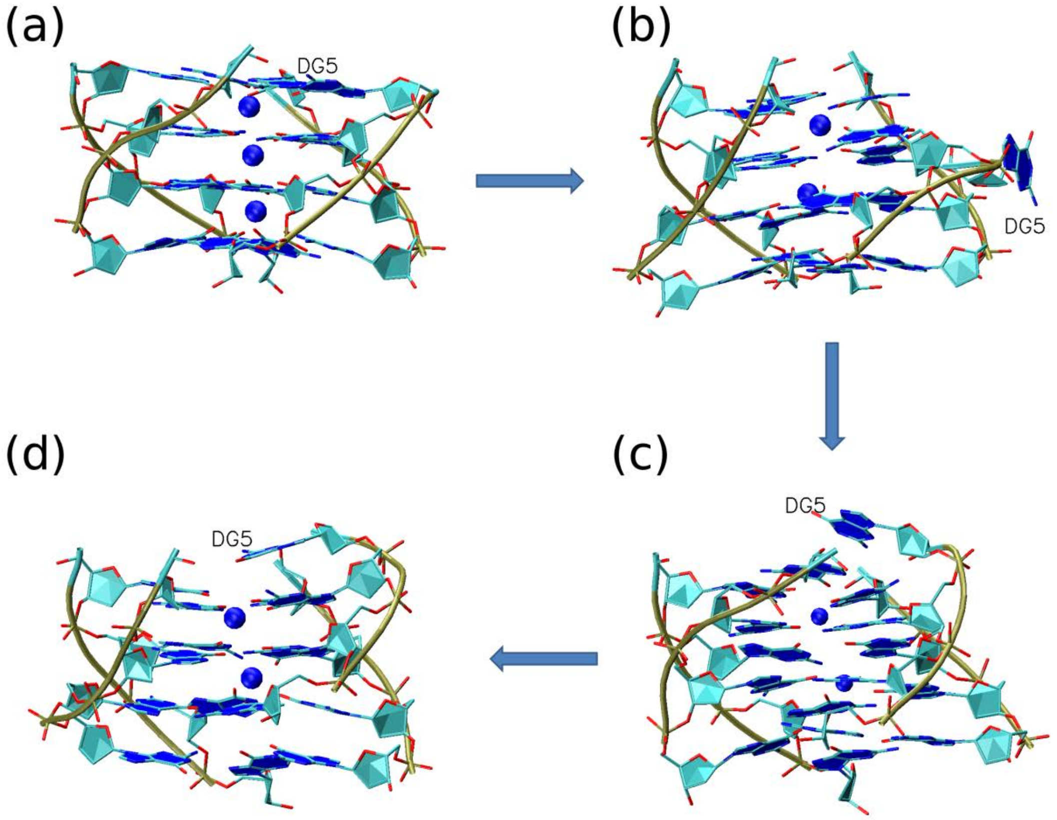


**Figure S34.** Sequence of events leading to strand slippage in parallel-stranded all-*anti* d[GGGG]$_4$ G-stem in the Simulation 24a carried out in $Na^+$ ions and OPC water model. The structure of G-stem at (a) start of the simulation, (b) 724 ns, (c) 804 ns and (d) 821 ns. Guanine 5 (DG5) of the G-stem moved towards the solvent (shown in panel b) and then stacked on the remaining terminal G-triad (shown in panel c). The movement of DG5 eventually enforced strand slippage (shown in panel d). The backbone of the G-stem is shown as tan tube with 5′ends in cyan tube. The guanine bases and $Na^+$ ions within the G-stem are shown in blue.

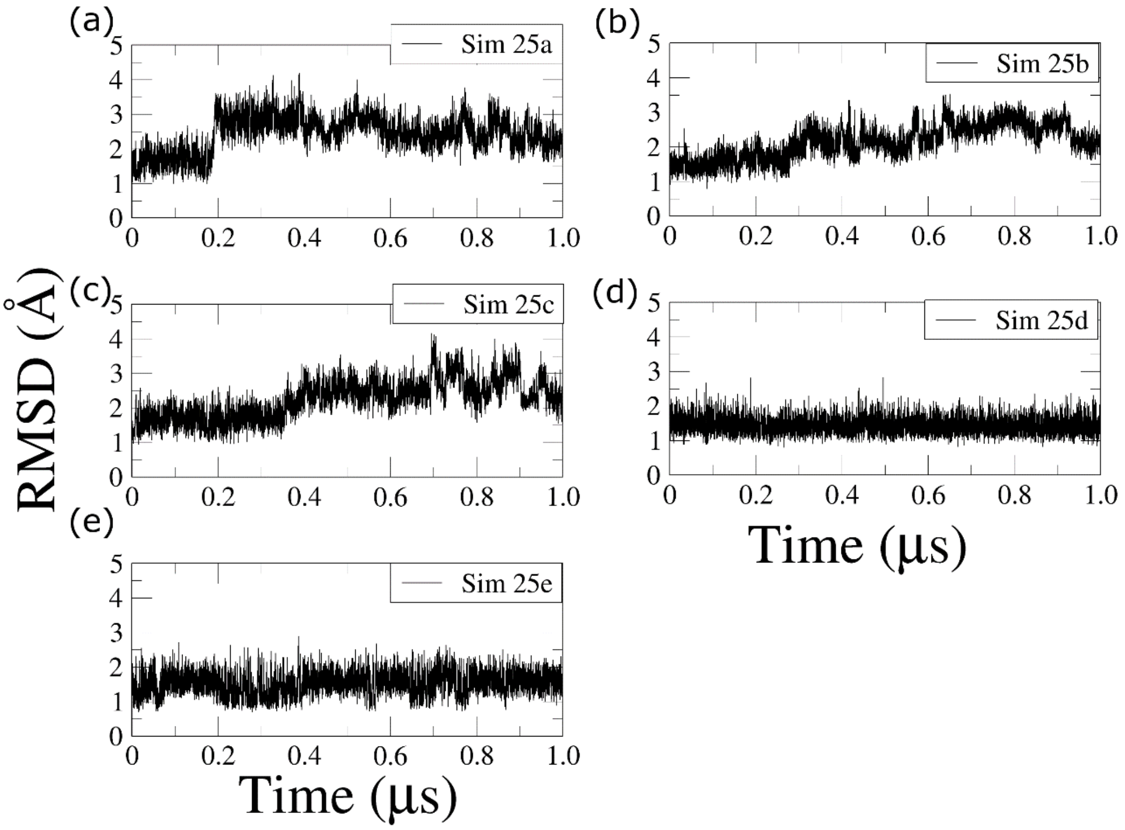


**Figure S35**. Backbone RMSD of parallel-stranded all *anti* d[GGGG]$_4$ four-quartet G-stem in the Simulations (a) 25a, (b) 25b, (c) 25c, (d) 25d and (e) 25e. Panels (a) - (e) show five independent simulations carried out in $K^+$ ions and OPC water model. The 5′-quartet was disrupted in the Simulations 25a- 25c but all the four quartets were maintained in the Simulations 25d and 25e.

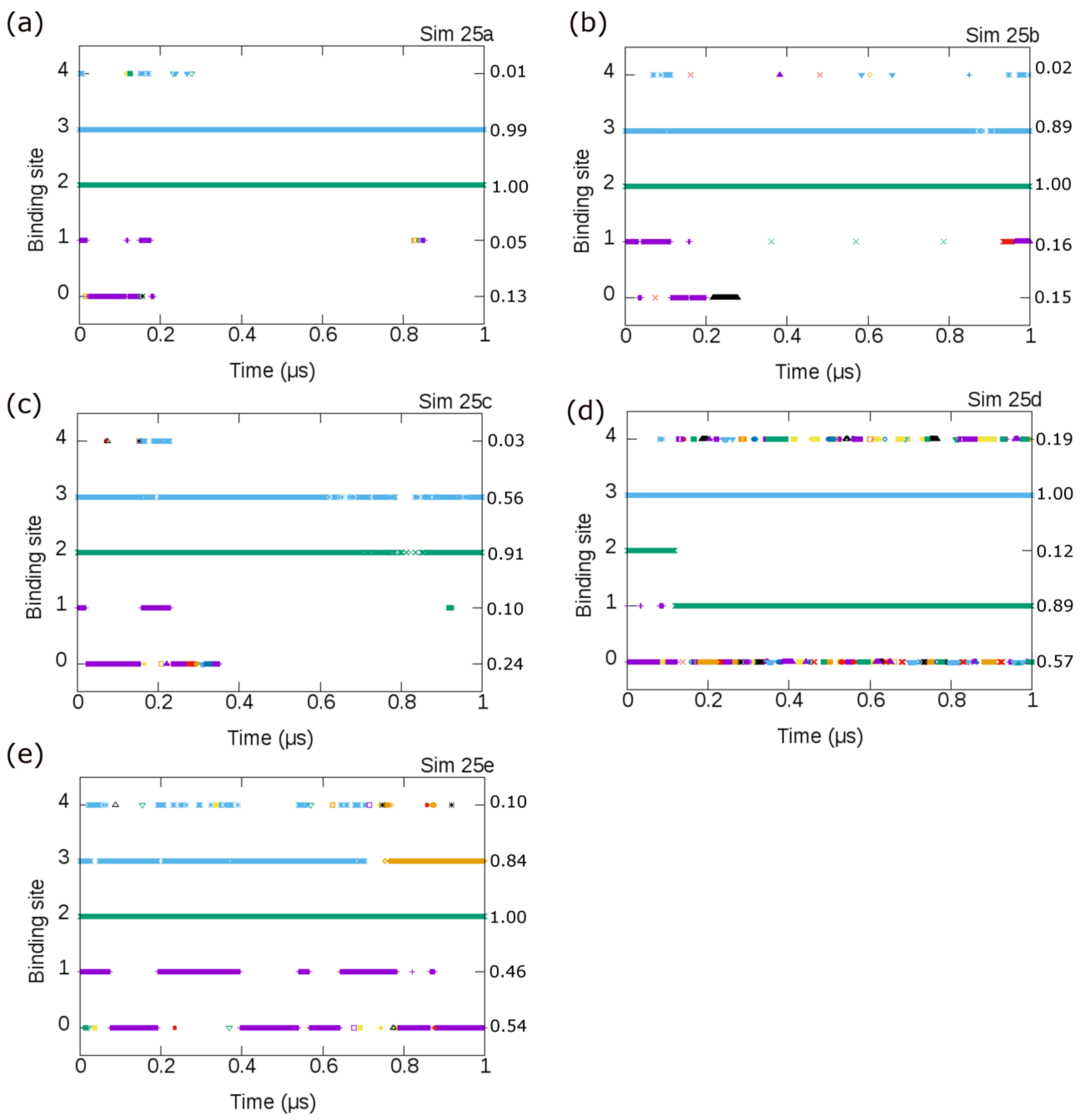


**Figure S36**. Cation exchanges in parallel-stranded all *anti* d[GGGG]$_4$ G-stem in the Simulations 25a - 25e. Simulations (a) 25a, (b) 25b, (c) 25c, (d) 25d and (e) 25e were carried out in $K^+$ ions and OPC water model. Distinct ions are represented by different colors and symbols, and any change in the color/symbol indicates an exchange of an ion in the site. The numbers on the right show occupancies of the respective sites. See legend to Figure S3 for details. In the Simulations 25a - 25c, Site 1 ion was pushed above the first quartet and eventually the first quartet of G-stem was disrupted. In the Simulation 25d, Site 2 was vacant while in the Simulation 25e the "purple" ion from Site 1 was pushed above the first quartet and entry of this cation back into the Site 1 pushed the "light blue" cation from Site 3 to transiently sample the position at the bottom of the third quartet, Site 4. Note, the simultaneous transient appearance of cations at two sites indicates the transient presence of cation within the plane of G-quartet shared by the corresponding sites.

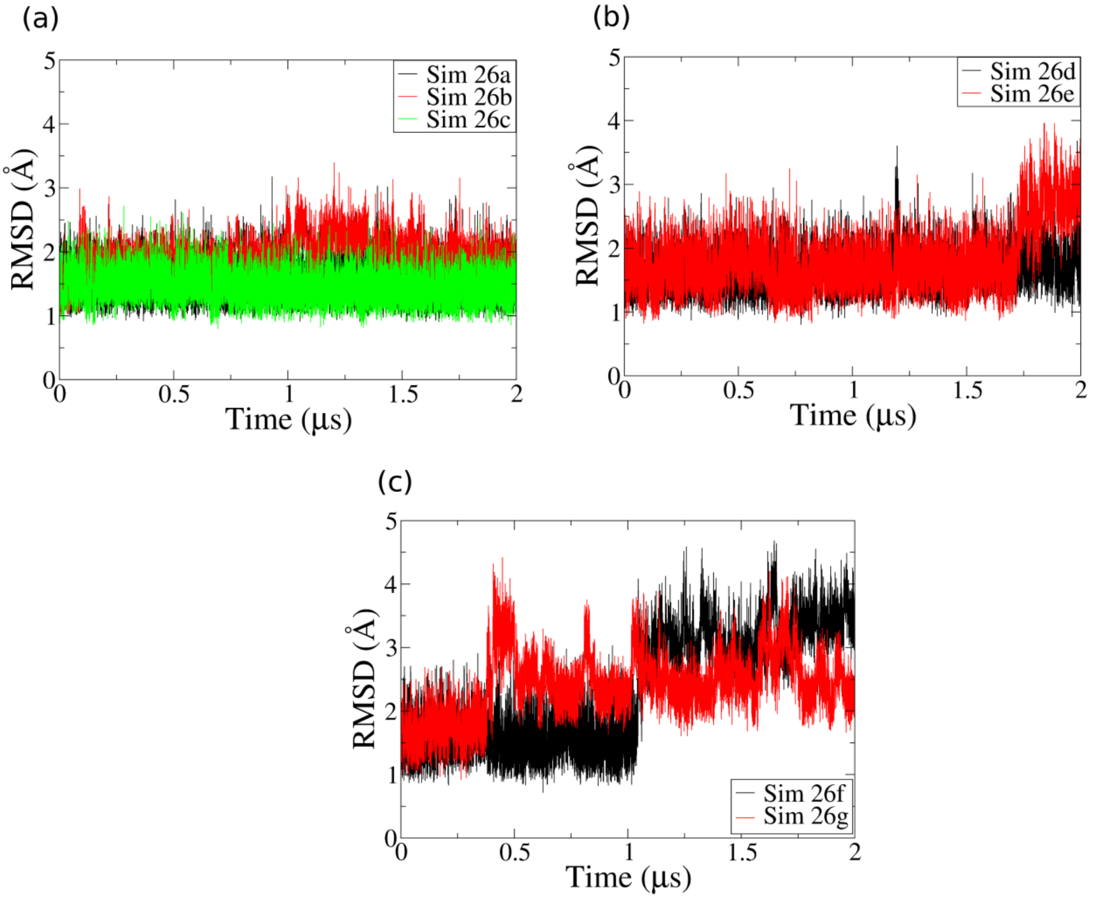


**Figure S37**. Backbone RMSD of parallel-stranded all *anti* d[GGGG]$_4$ four-quartet G-stem derived from PDB 4R44 in the Simulations 26a - 26g. Panels (a) Li/Merz $Na^+$ (black), $K^+$ ions (Simulation 26b and 26c in red and green, respectively) and SPC/E water model, (b) HFE $Na^+$ (black) and $K^+$ (red) ion and OPC water model and (c) IOD $Na^+$ (black) and $K^+$ (red) ion and OPC water model. The G-stem was stable in Li/Merz $Na^+$ ion simulation in SPC/E water model and HFE $Na^+$ ions and OPC water model. The fourth quartet was disrupted at 90 ns in Li/Merz $K^+$ ion simulation in SPC/E water model (shown in red in panel a) but was maintained in the second simulation in the same conditions (shown in green in panel a). Disruption of 5′-quartet was observed at 1.7 μs in the simulation with HFE $K^+$ ions and OPC water model (shown in red in panel b). 5′-quartet was also disrupted at 1 μs and 0.4 μs in IOD $Na^+$ and $K^+$ ions in the OPC water model, respectively as shown in panel c.

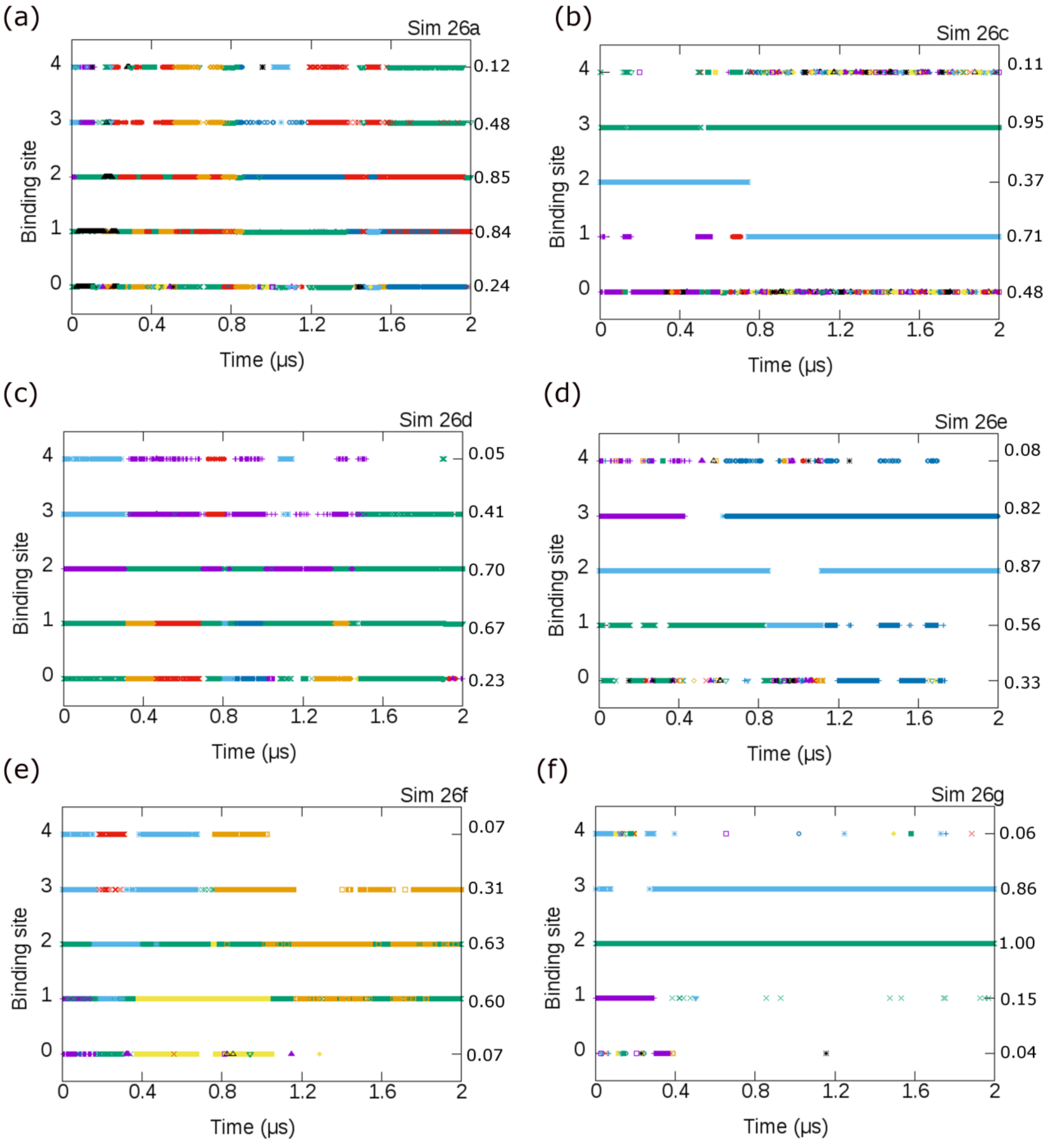


**Figure S38**. Plots monitoring the cation binding in parallel-stranded all *anti* d[GGGG]$_4$ G-stem in the Simulations 26a and 26c - 26g. Simulations were carried out in (a) Li/Merz $Na^+$ and SPC/E (Simulation 26a), (b) Li/Merz $K^+$ ion and SPC/E (Simulation 26c), (c) HFE $Na^+$ and OPC (Simulation 26d), (d) HFE $K^+$ ion and OPC (Simulation 26e), (e) IOD $Na^+$ and OPC (Simulation 26f) and (f) IOD $K^+$ ion and OPC (Simulation 26g) water model. In the two simulations carried out in $K^+$ Li/Merz ions and SPC/E water model, the fourth quartet disrupted in Simulation 26b (not shown) but was stable in 26c; the cation dynamics in Simulation 26c is shown in the figure (panel b). Distinct ions are represented by different colors and symbols, and any change in the color/symbol indicates an exchange of an ion in the site. The numbers on the right show occupancies of the respective sites. See legend to Figure S3 for details. G-stem was fully maintained only in the Simulations 26a, 26c and 26d. More frequent cation exchanges were observed in the stable simulations in $Na^+$ ions (panels a and c) than in the $K^+$ ions (panel b).

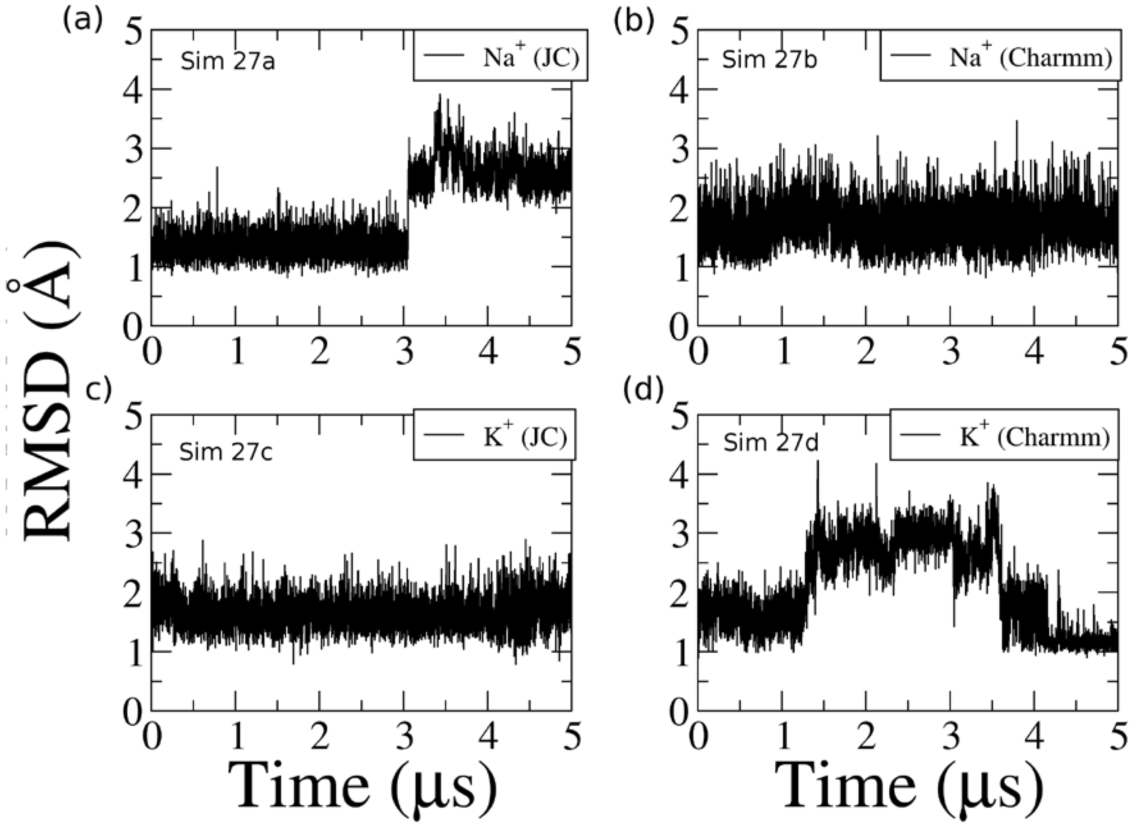


**Figure S39**. Backbone RMSD of parallel-stranded all *anti* d[GGGG]$_4$ G-stem in the Simulations (a) 27a, (b) 27b, (c) 27c and (d) 27d. The simulations were carried out in TIP4P-D water and JC $Na^+$, CHARMM22 $Na^+$, JC $K^+$ and CHARMM22 $K^+$ ions. The G-stem was stable in CHARMM22 $Na^+$ and JC $K^+$ ions (Simulations 27b and 27c). G-stem was perturbed in the simulation in JC $Na^+$ ions by vertical strand slippage (Simulation 27a). 5′-quartet was disrupted in the simulation in CHARMM22 $K^+$ ions at ~1.3 µs though the structure repaired spontaneously at 3.6 µs (Simulation 27d).

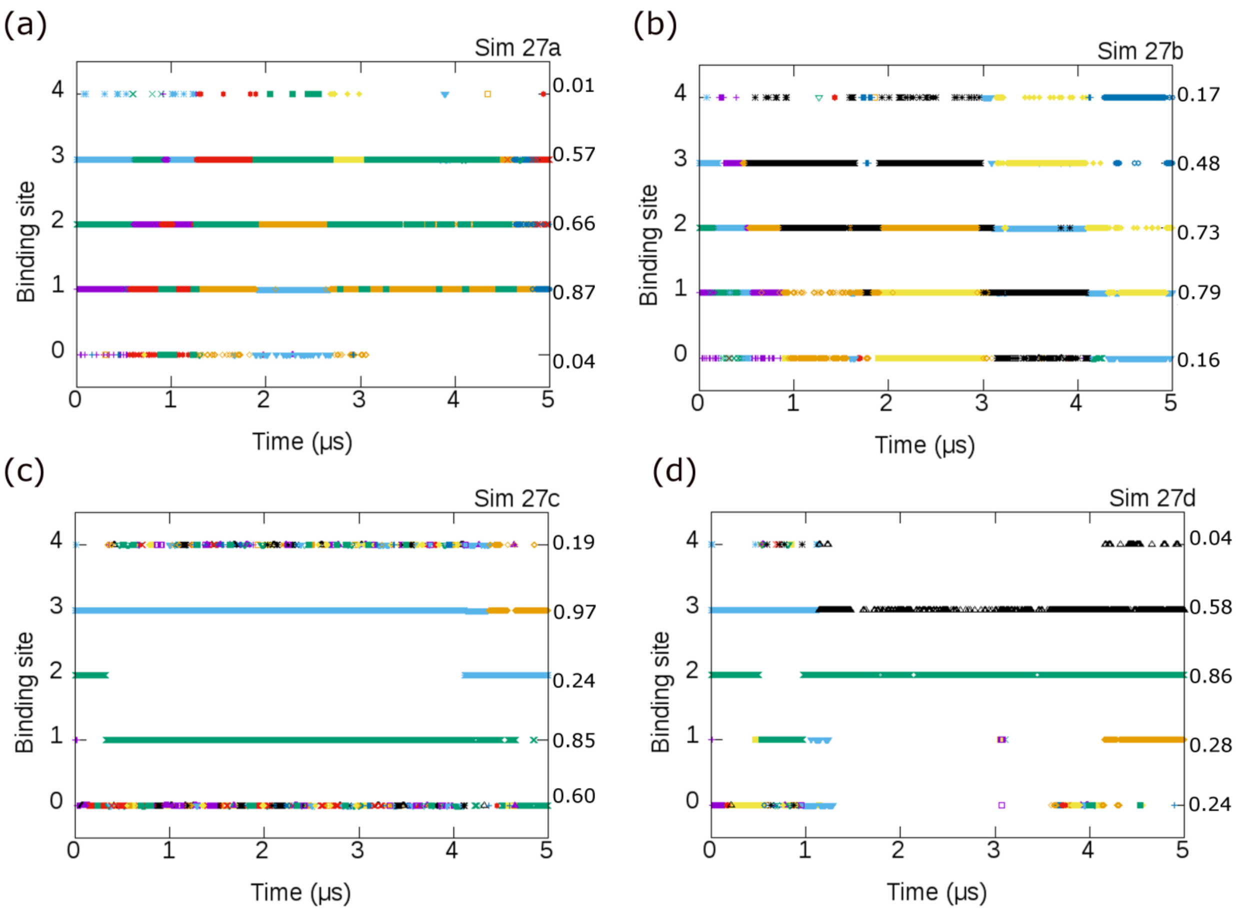


**Figure S40**. Plots monitoring cation binding in the simulations of parallel-stranded all *anti* d[GGGG]$_4$ G-stem in the Simulations 27a - 27d. The simulations were carried out in TIP4P-D water and (a) JC $Na^+$, (b) CHARMM22 $Na^+$, (c) JC $K^+$ and (d) CHARMM22 $K^+$ ions. Multiple cation exchanges were observed in the simulations in $Na^+$ ions with both the parameters. G-stem was stable only in Simulations 27b (panel b) and 27c (panel c). Distinct ions are represented by different colors and symbols, and any change in the color/symbol indicates an exchange of an ion in the site. The numbers on the right show occupancies of the respective sites. See legend to Figure S3 for details.

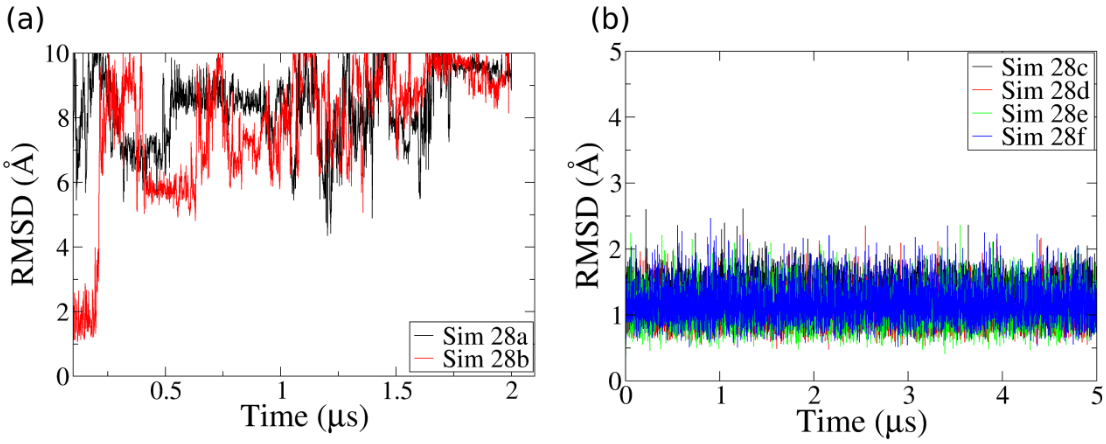


**Figure S41.** Backbone RMSD of the two-quartet d[GG]$_4$ (a) parallel-stranded and (b) antiparallel G-stem in the Simulations 28a - 28f. The temperature in all the simulations of this series was maintained using Langevin thermostat. The two independent simulations of parallel-stranded all *anti* d[GG]$_4$ were carried out in $K^+$ ions with the SPC/E water model. The four independent simulations of antiparallel d[GG]$_4$ were carried out in $Na^+$ ions with the SPC/E water model. Strand slippage and G-stem unfolding was observed in both the simulations of parallel-stranded d[GG]$_4$ as is shown in panel a by the significant change in RMSD at 0.1 and 0.2 μs. The antiparallel d[GG]$_4$ G-stem was stable in all the four simulations.

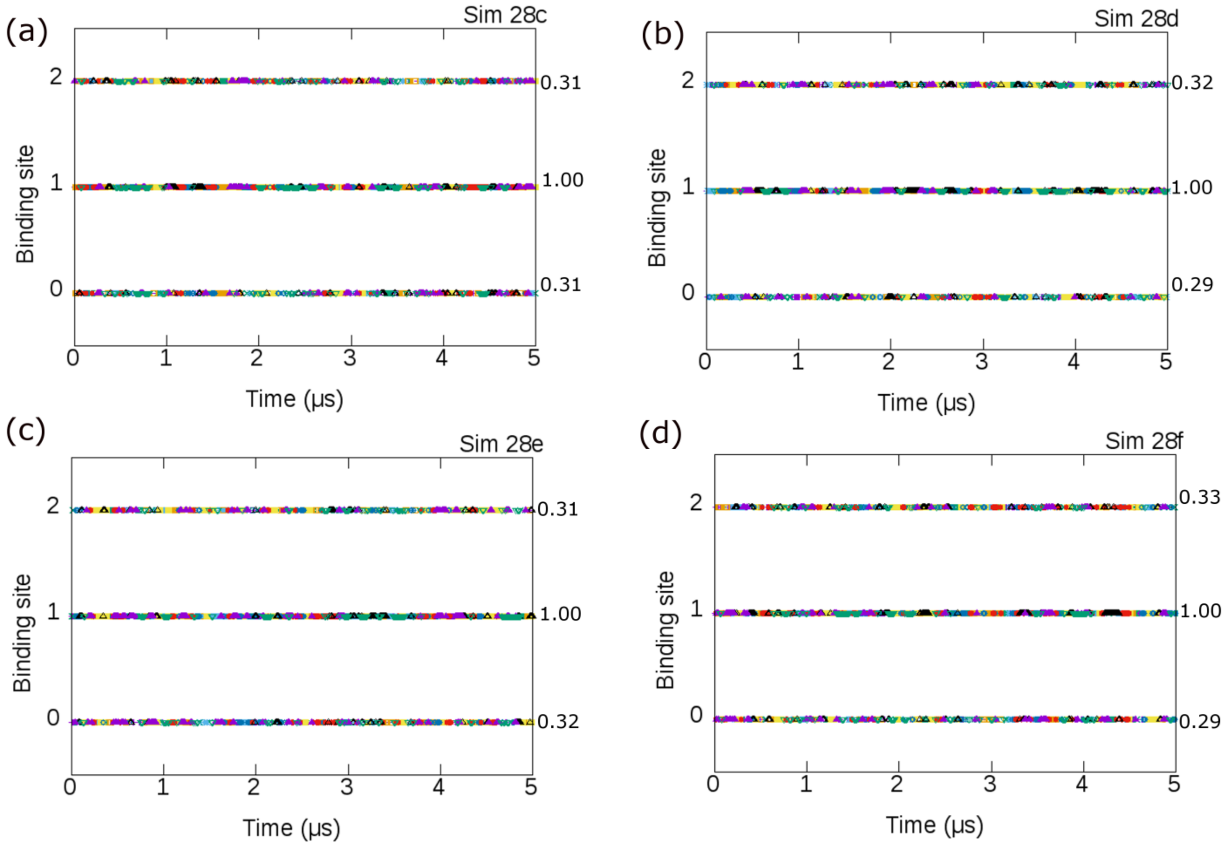


**Figure S42**. Plots monitoring cation movement in the simulations of antiparallel d[GG]$_4$ Simulations (a) 28c, (b) 28d, (c) 28e and (d) 28f. The simulations were carried out in $Na^+$ ions and SPC/E water model and the temperature was maintained using the Langevin thermostat. Distinct ions are represented by different colors and symbols, and any change in the color/symbol indicates an exchange of an ion in the site. The numbers on the right show occupancies of the respective sites. See legend to Figure 4 in the Main manuscript for details. The plots showed that there were very frequent ion-exchange movements between the G-stem and the solvent in all the four simulations similar to the equivalent simulations carried out with Berendsen thermostat (Simulations 7a-7e).

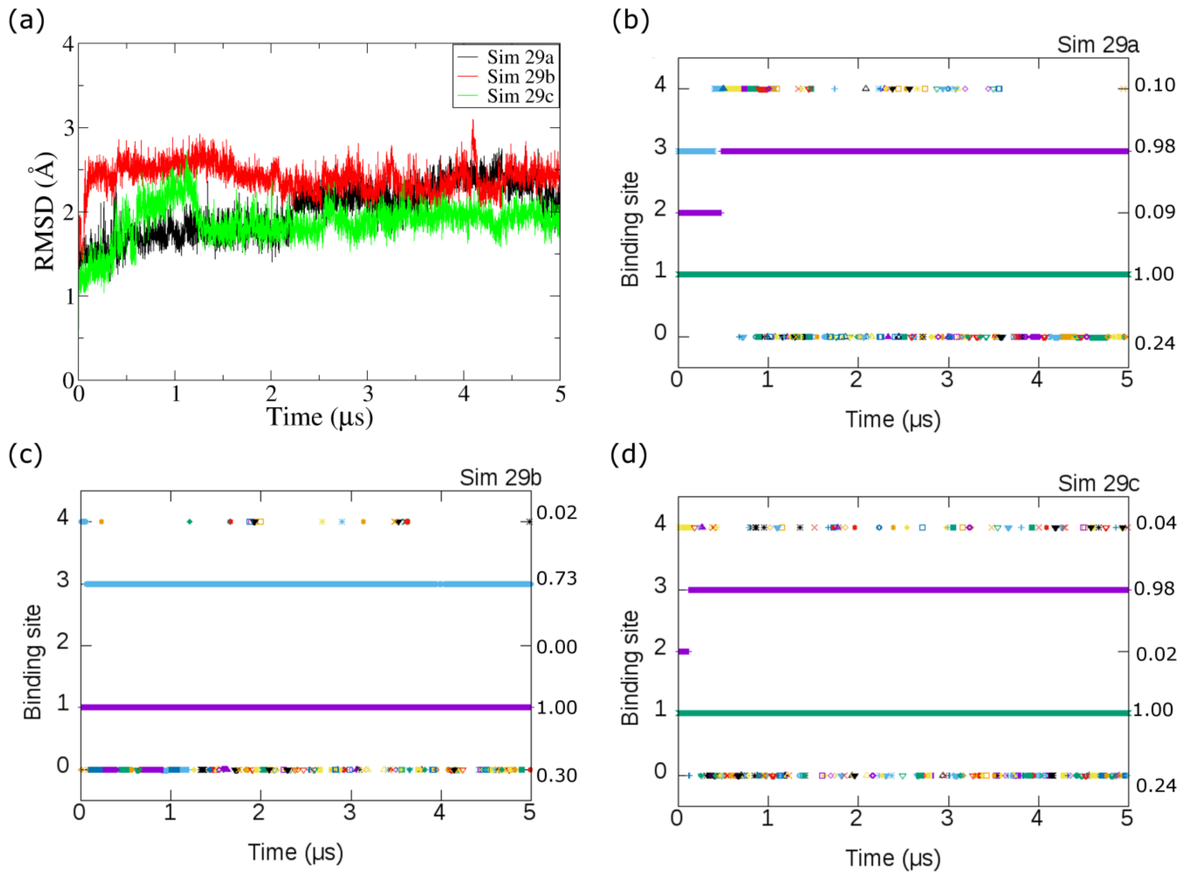


**Figure S43**. Backbone RMSD (a) and plots monitoring cation movement in the Simulations (b) 29a, (c) 29b, and (d) 29c of complete 2RQJ. The simulations were carried out in $K^+$ ions and SPC/E water model and various initial ion positions were considered. $K^+$ ions were placed in all the three sites in the Simulation 29a while it was placed in only Site 1 in the Simulation 29b and Sites 1 and 2 in the Simulation 29c. In the panels b - d, the numbers on the right show occupancies of the respective sites. The temperature was maintained using the Langevin thermostat in all the simulations. (a) No major backbone RMSD changes showed that the GQ was stable in the simulations. (b - d) The plots showed that eventually, only two cations (at Sites 1 and 3) stabilized the GQ similar to the equivalent simulations carried out with Berendsen thermostat (see Figure S13). See legend to Figure S3 for details.

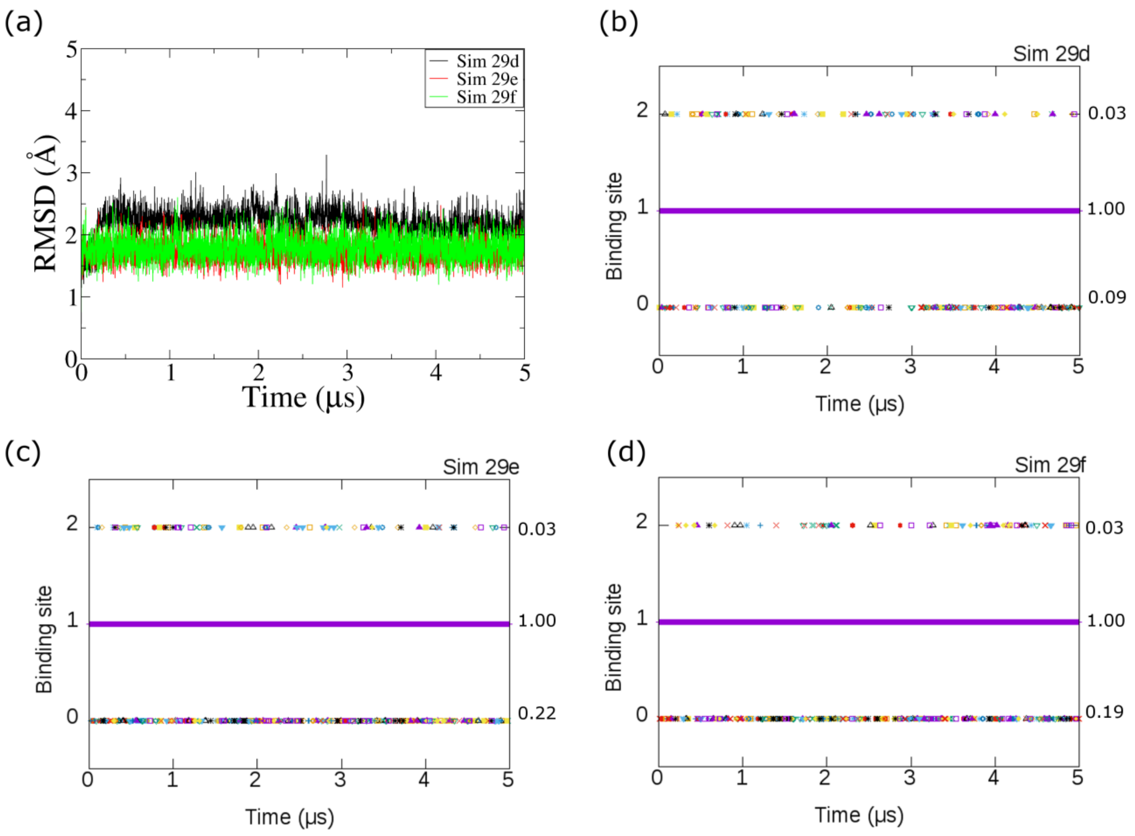


**Figure S44**. Backbone RMSD (a) and plots monitoring cation movement in the simulations of two quartet RNA GQ monomer from PDB 2RQJ in the Simulations (b) 29d, (c) 29e, and (d) 29f. The simulations were carried out in $K^+$ ions and SPC/E water model and the temperature was maintained using the Langevin thermostat. (a) No major backbone RMSD changes showed that the GQ was stable in the simulations. (b - d) The plots showed that one cation was retained between the two quartets and there was no exchange with the solvent similar to the equivalent simulation carried out with Berendsen thermostat (see Figure S19). The numbers on the right show occupancies of the respective sites. See legend to Figure 4 in the Main manuscript for details.

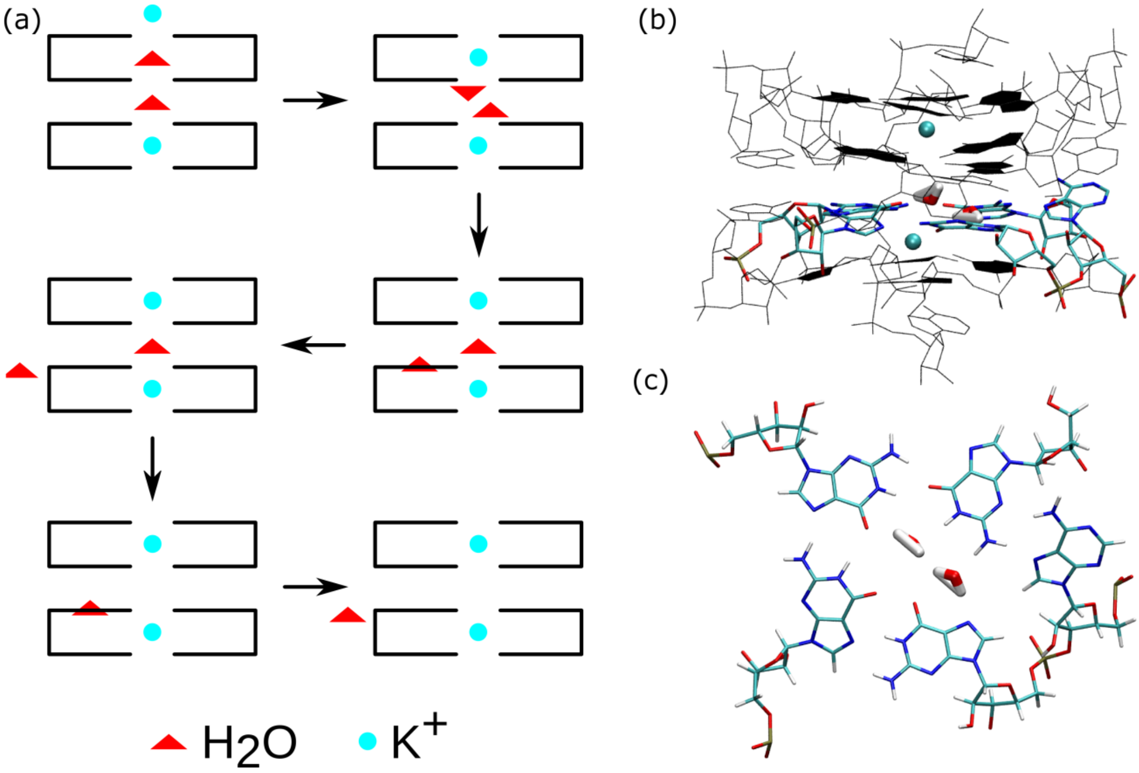


**Figure S45**. Water expulsion from the channel of a dimeric RNA GQ (2RQJ). (a) A sequence of events starting with one cation and two water molecules in the channel and an incoming cation, ending by both water molecules repelled from the channel into the groove. (b) Snapshot of the intermediate with cations (cyan) in Sites 1 and 3, one water molecule in Site 2 and one water molecule embedded into one of the middle quartets, which is also hydrogen-bonded with a loop adenine (shown in the licorice representation). (c) View of the quartet-water-adenine arrangement. The snapshot is taken from Simulation 8b.

**REFERENCES**

1. Stadlbauer, P.; Krepl, M.; Cheatham, T. E., 3rd; Koča, J.; Šponer, J., Structural dynamics of possible late-stage intermediates in folding of quadruplex DNA studied by molecular simulations. *Nucleic Acids Res.* **2013**, 41, 7128-7143.

2. Stefl, R.; Cheatham, T. E., 3rd; Spacková, N.; Fadrná, E.; Berger, I.; Koca, J.; Sponer, J., Formation pathways of a guanine-quadruplex DNA revealed by molecular dynamics and thermodynamic analysis of the substates. *Biophysical journal* **2003**, 85, 1787-1804.
